# Efficient reliability analysis of spatially resolved transcriptomics at varying resolutions using SpaSEG

**DOI:** 10.1101/2022.11.16.516728

**Authors:** Yong Bai, Xiangyu Guo, Keyin Liu, Bingjie Zheng, Yingyue Wang, Qiuhong Luo, Jianhua Yin, Liang Wu, Yuxiang Li, Yong Zhang, Ao Chen, Xun Xu, Xin Jin

## Abstract

Spatially resolved transcriptomics (SRT) for characterizing cellular heterogeneities and activities requires systematic analysis approaches to decipher gene expression variations in physiological contexts. Here we develop SpaSEG, an unsupervised convolutional neural network-based model for multiple SRT analysis tasks by jointly learning the transcriptional similarity of spots and their spatial dependence. SpaSEG adopts an edge strength constraint to encourage spatial domain coherence and allows integrative analysis by automatically aligning the spatial domains across multiple adjacent sections. It also enables the detection of domain-specific gene expression patterns and the inference of intercellular interactions and colocalizations within a tissue. In an invasive ductal carcinoma sample analysis, SpaSEG facilitates the unraveling of intratumor heterogeneity and the understanding of immunoregulatory mechanisms. Through comprehensive evaluation over a collection of SRT datasets generated by different platforms at various resolutions, SpaSEG shows superior reliability and computational efficiency over existing methods, endowing it with a great potential for the exploration of tissue architectures and pathological biology.

## Introduction

Coordinated activities and functions of diverse cells within a tissue or organ context that underlie their complex communications with surroundings and sophisticated biological processes can be characterized by gene expression patterns organized in spatial territories^1–3^. Emerging spatially resolved transcriptomics (SRT) technologies have enabled unbiased profiling of transcriptome-wide gene expression with physical capture spots, offering spatial snapshots of cellular heterogeneity across tissue sections^4, 5^. Recent years have witnessed tremendous progress in various SRT experimental methods^6^, including the imaging-based transcriptomics approaches like MERFISH^7^ and seqFISH^8^, and the next-generation sequencing-based approaches such as Slide-seqV2 (ref.^9^), 10x Visium^10^ and Stereo-seq^11^. Although these SRT technologies have reached an unprecedented resolution from multicellular to single-cell or even subcellular level with high gene throughput, they inevitably bring new challenges for data analysis^12^.

A pivotal task in SRT data analysis is to identify spatial domains defined as regions with coherence in gene expression and spatial dependence. The spatial domains serve uncovering tissue structures and functions, facilitating the characterization of cell type compositions and transcriptomic profiles, as well as the inference of intercellular interactions within the tissue^6, 13^. Conventional approaches to arranging spots into distinct spatial domains resort to non-spatial clustering methods such as Leiden^14^ that only take into account gene expressions of spots^15^. To incorporate gene expression with spatial information, several methods have been developed, including SEDR^16^, BayesSpace^17^, SpaGCN^18^, stLearn^19^, and Giotto^20^, to name a few. Despite the promising performance, these spatial clustering methods are only applied on the small SRT dataset with limited spots and thus, presumably fragile for the large assayed tissue size at high capture resolution. Besides, the spatial clustering accuracy is often restricted by the discontinuity in spatial domains owing to the lack of boundary smoothness constraints and thus susceptible to noise.

Integrating SRT datasets for joint analysis of multiple adjacent tissue sections is of advantage as it promises the characterization of variations in gene expression from different sections. A critical issue in such integrative analysis is the comparison of tissue structures, which requires the correspondence and consistency of spatial domains across adjacent sections^21^. Such cross-domain correspondence allows spatial registration of tissue organization from adjacent sections, simultaneous investigation of regions of interest (ROI), and reconstruction of an organ transcriptome atlas in a 3-dimensional (3D) space along an axis. Nonetheless, due to the lack of methods, several studies involving multiple adjacent sections either examine them individually^22, 23^, or simply aggregate gene expression of spots together without regard of spatial coordinates that define the rigid structure of a tissue in the SRT data^15^.

Understanding biological functions associated with spatial domains necessitates pinpointing genes that exhibit spatial expression variations and patterns known as spatially variable genes^24^ (SVGs). A handful of methods such as trendsceek^25^, SpatialDE^26^ and SPARK^27^ have been pursued to detect SVG primarily based on statistical modeling. These methods establish the dependence between the gene expressions and spatial locations, and subsequently perform statistical tests to screen significantly differentially expressed genes. By contrast, SpaGCN^18^ detects SVGs by investigating differential gene expressions between spatial domains using a deep graph convolutional network. Moreover, spatial variations in gene expression across spatial domains related to regional functions imply cell-cell interactions^13, 24^ (CCIs). Nevertheless, the majority of existing studies analyzing CCIs using SRT data rely on the approaches originating from scRNA-seq data^15, 28^. This would hamper the interrogation of the spatial CCI map and cell colocalization within an intact tissue due to the absence of examining the spatial proximity of cells^3, 29^.

Here, we developed SpaSEG, a simple yet powerful unsupervised convolutional neural network (CNN)-based method for SRT analysis by jointly learning the similarity of gene expressions and their spatial dependence. In brief, SpaSEG iteratively identifies spatial domains based on an image-like tensor derived from SRT data. It guarantees the continuity of spatial domains by minimizing the edge strength of the image-like tensor. By extensively analyzing a collection of SRT datasets generated by a range of platforms at various resolutions, we demonstrate that SpaSEG exhibits superior performance over the existing state-of-the-art methods. In addition, SpaSEG can automatically align spatial domains across multiple adjacent sections, leading to the correspondence to and consistency with their tissue structures and thereby, allowing the integration of multiple SRT datasets for deciphering tissue structures in 3D space. Based on spatial domains, SpaSEG detects SVGs for elucidating domain-specific gene expression patterns and examines spot-wise CCI map and cell colocalization within the tissue. In an invasive ductal carcinoma analysis, SpaSEG illustrates inter- and intra-tumor heterogeneities and facilitates understanding of immunoregulatory mechanisms. Through comprehensive evaluation, we show that SpaSEG is remarkably computationally efficient and platform-agnostic, and hence well applicable to diverse SRT data for multiple analysis tasks, serving as an appealing tool to explore tissue architecture and cellular characterization for different assayed tissue sizes at various resolutions.

## Results

### Overview of SpaSEG

SpaSEG started with preprocessing of raw spatial transcriptomic data by excluding low-quality genes and poor spots, followed by log-normalization of gene expression profiles, principal component analysis (PCA) and *z*-score scaling, yielding a low-dimensional feature vector ***s***_*n*_ ∈ ℝ^*d*^ for each spot *n* (Fig. 1a). SpaSEG converted the feature vectors into an image-like tensor, whereby spots were analogous to the image pixels while the corresponding feature vectors to channels. SpaSEG responsible for spatial domain identification was a CNN-based network model that consisted of a batch normalization layer, two stacking convolutional blocks and a refinement module, mapping each feature vector ***s***_*n*_ to a latent representation ***y***_*n*_ ∈ ℝ^*d*^ (Fig.1b). After pre-training SpaSEG for the initialization of parameters using the mean squared error loss between ***s***_*n*_ and ***y***_*n*_, we optimized a joint loss of weighted sum of the cross-entropy loss ℒ_seg_ with L2-norm regularization and the edge strength loss ℒ_edge_ in the subsequent training iterations to progressively enhance the spot-wise classification while promoting the coherence of domains (Methods). Consequently, spots that presented similar gene expressions along with spatially continuous coordinates were clustered into the same domain. The downstream analyses such as multiple adjacent sections integration, SVG detection and spatial CCI inference can be further investigated based on the identified spatial domains (Fig.1c).

**Fig. 1.**
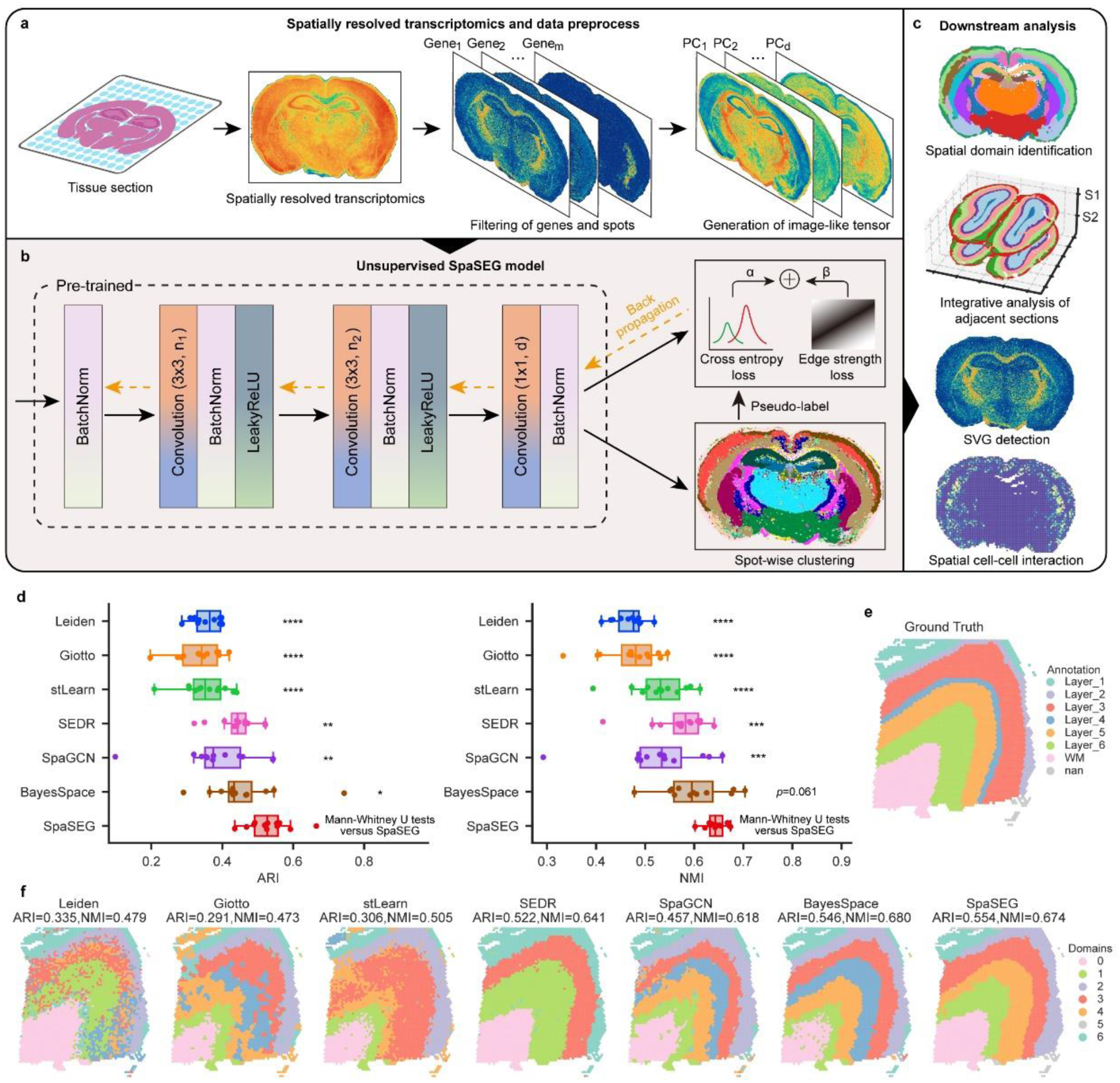
Overview of SpaSEG. **a,** Spatially resolved transcriptomics data preprocessing and image-like tensor preparation. **b**, Unsupervised SpaSEG neural network with two loss functions used to jointly learn transcriptional similarity and spatial dependence iteratively. **c,** Downstream analysis tasks including spatial domain identification, integrating multiple adjacent sections, SVG detection, and spatial cell-cell interaction inference. **d**, Boxplots of ARI and NMI evaluation metrics showing the accuracy of SpaSEG in identifying spatial domains in all 12 sections of the 10x Visium DLPFC dataset against competing methods. * p<0.05, ** p<0.01, *** p<0.001, **** p<0.0001. **e**, Ground truth annotation of the DLPFC section 151673. **f**, Visualization of spatial domains identified by SpaSEG and competing methods for section 151673. Corresponding ARI and NMI values are shown.

### SpaSEG improves spatial domain identification in the human dorsolateral prefrontal cortex dataset

To evaluate SpaSEG on the spatial domain identification, we first exploited the 10x Visium human dorsolateral prefrontal cortex^22^ (DLPFC) as a benchmark dataset. This manually annotated dataset consisted of 12 sections collected from three subjects that covered six neuron layers and white matter (Supplementary Table 1). We chose the commonly used non-spatial clustering method Leiden, and other five recently published state-of-the-art methods including Giotto, stLearn, SEDR, SpaGCN, and BayesSpace for comparison. Apart from the qualitative visualization analysis, two widely used clustering evaluation metrics of adjusted rand index (ARI) and normalized mutual information (NMI) were used to quantify the clustering accuracy.

With a low computational expense (Supplementary Fig. 1), SpaSEG generally significantly outperformed the competing methods on all 12 DLPFC sections in terms of the highest ARI (0.527 ± 0.061; p<0.05) and NMI (0.643 ± 0.021; p<0.0001 except BayesSpace) (Fig. 1d, Supplementary Tables 2, 3), as well as the neatest spatial domains with clear boundaries in large agreement to tissue structures (Supplementary Fig. 2). For example, SpaSEG-identified spatial domains in a representative section 151673 displayed an excellent consistency with the annotations (ARI=0.554 and NMI=0.674; Figs. 1e, f). All methods struggled to discern layer 4 from others presumably because of an extremely small fraction of spots that may have genes expressed similarly to the adjacent layers. Despite acceptable clustering accuracy, SpaGCN and BayesSpace appeared to improperly separate the white matter into two domains with ragged boundaries, while SEDR incorrectly merged layers 4, 5 and 6 into a single domain. The spatial domains resulting from Leiden, stLearn and Giotto massively mixed many unexpected outliers, leading to rough tissue structures and poor performance.

### SpaSEG enables spatial domain identification for SRT datasets generated by different platforms

Next, we assessed the reliability of SpaSEG for identifying spatial domains in discrepant SRT datasets. To do this, we collected four published SRT datasets generated by different platforms embracing a series of technological and biological characteristics (Methods). These SRT datasets captured a broad range of spots and genes, from 6412 to 53,208 and 161 to 25,879, respectively (Supplementary Table 1). Considering both clustering accuracy and concordance of spatial domains achieved above, we benchmarked SpaSEG with SpaGCN and BayesSpace, as well as Leiden serving as a baseline method in this analysis.

On the mouse hemibrain Stereo-seq data at CellBin resolution^11^, SpaSEG well depicted the spatial distribution of neurons and non-neuronal cells compared to Leiden and SpaGCN (Fig. 2a, Supplementary Fig. 3a). Specifically, it clearly outlined EX CA (domain 6), GN DG (domain 2), EX thalamus (domain 9) and Oligo (domain 0) that were marked by *Neurod6*, *Spink8*, *Prox1*, *Lrrtm4*, *Prkcd*, and *Mbp* (Fig. 2b, Supplementary Fig. 4a). BayesSpace did not yet successfully conduct spatial clustering on this dataset probably due to its inadaptation to the Stereo-seq data with a large number of spots in a non-lattice arrangement. In the mouse somite-stage embryo seqFISH data^30^, SpaSEG accurately uncovered the spatial regions in contrast to other methods (Fig. 2c, Supplementary Fig. 3b), including forebrain (domain 9, Fig. 2d), midbrain (domain 20), hindbrain (domain 0) and spinal cord (domains 10 and 11) that were marked by *Six3*, *Lhx2*, *Otx2* and *Pou3f1*, as well as *Sox2* (ref.^30^, Supplementary Fig. 4b). Likewise, using the mouse hypothalamic preoptic region MERFISH data^31^, we found that SpaSEG better delineated the spatial distribution of different cell types than other methods (Fig. 2e, Supplementary Fig. 3c), such as ependymal cells marked by *Cd24a* (domain 4, Fig. 2f, Supplementary Fig. 4c), mature OD by *Mbp* (domain 3), excitatory cells by *Slc17a6* (domain 2), and inhibitory cells by *Gad1* (domains 5 and 7). Results of ARI showed the superior clustering accuracy of SpaSEG over the competing methods (Fig. 2i). We further applied SpaSEG on the mouse hippocampus Slide-seqV2 data^9^. As expected, SpaSEG can better outline the topology of the tissue along with the neat spatial domains and sharp boundaries (Fig. 2g, Supplementary Fig. 3d). For instance, in addition to cortical layers marked by *Mef2c* (domains 10 and 15, Fig. 2h, Supplementary Fig. 4d), SpaSEG clearly outlined the Ammon’s horn marked by *Neurod6*, *Spink8* and *Hs3st4* (domain 2 and 9), DG-sg by *Lrrtm4* and *Prox1* (domain 1), V3 by *Ttr* (domain 4), and MH and LH by *Nwd2* (domain 8). We used the local inverse Simpson’s index (LISI) to measure the clustering accuracy on such SRT data without manual annotations. The significantly low LISI value reached by SpaSEG highlighted its optimal spatial clustering performance (p<0.001, Fig. 2j). Altogether, these results demonstrated the great reliability and accuracy of SpaSEG in the identification of spatial domains for SRT datasets generated by different platforms, suggesting its platform-agnostic capability.

**Fig. 2.**
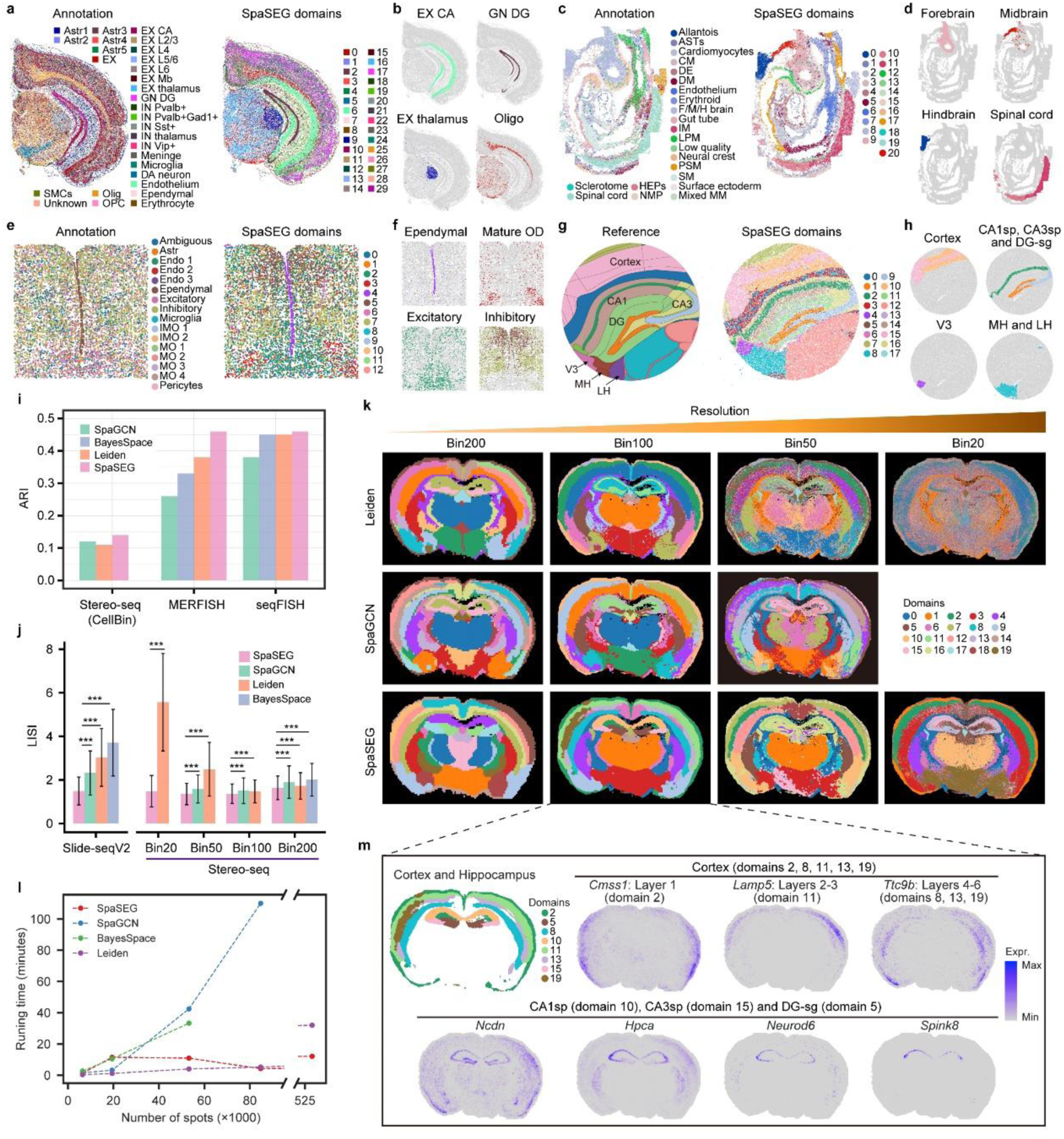
The identification of spatial domains in diverse SRT datasets generated by different SRT platforms. **a-h**, Annotations and SpaSEG-identified spatial domains, as well as domain examples in the mouse hemibrain Stereo-seq data at CellBin resolution (**a, b**), mouse somite-stage embryo seqFISH data (**c, d**), mouse hypothalamic preoptic region MERFISH data (**e, f**), and mouse hippocampus Slide-seqV2 data (**g, h**). We employed the annotations from Allen Brain Atlas^33^ as a reference for the mouse hippocampus. Astr: astrocyte, EX: excitatory, IN: inhibitory, CA: cornu ammonis, Mb: midbrain, GN DG: granule cell of dentate gyrus, DA: dopaminergic, Oligo: oligodendrocyte, OPC: oligodendrocyte precursor cell, SMCs: smooth muscle cells, ASTs: anterior somitic tissues, CM: cranial mesoderm, DE: definitive endoderm, F/M/H brain: fore/mid/hind brain, IM: intermediate mesoderm, LPM: lateral plate mesoderm, PSM: presomitic mesoderm, SM: splanchnic mesoderm, Mixed MM: mixed mesenchymal mesoderm, HEPs: haematoendothellal progenitors, NMP: neuromesodermal progenitors, IMO: immature oligodendrocytes; MO: mature oligodendrocytes, V3: third ventricle, MH: medial habenula, LH: lateral habenula. **i**, Comparison of ARI values for SpaGCN, BayesSpace, Leiden and SpaSEG in the SRT datasets generated by Stereo-seq (CellBin), MERFISH and seqFISH. **j**, LISI values showing the accuracy of spatial domains identified by SpaSEG, SpaGCN, Leiden, and BayesSpace for mouse hippocampus Slide-seqV2 data (Left panel) and whole adult mouse brain Stereo-seq data at various resolutions (Right panel). *** p<0.001. **k**, Illustration of spatial domains identified by Leiden, SpaGCN, and SpaSEG for the whole adult mouse brain Stereo-seq data at resolutions of Bin200, Bin100, Bin50, and Bin20. **l**, Running time for spatial domain identification used by SpaSEG, SpaGCN, BayesSpace and Leiden against varying numbers of spots. The algorithm is terminated if its runtime exceeds 6 hours. **m**, Spatial expression patterns of selected markers for the cerebral cortex and hippocampus in the adult mouse brain Stereo-seq data at Bin100 resolution.

### SpaSEG efficiently reveals tissue structures in large sections at high resolution

Next, we examined the scalability and efficiency of SpaSEG on the large tissue section at high resolution. To do this, we analyzed a published whole adult mouse brain Stereo-seq data^32^ assayed in a large area of 1cm × 1cm. To facilitate our analysis at different resolutions, we intentionally diversified this data into four datasets at resolutions of Bin200 (100μm diameter), Bin100 (50μm diameter), Bin50 (25μm diameter) and Bin20 (10μm diameter) (Methods), spanning the number of spots from 5420 to 526,716 (Supplementary Table 1). We used the coronal section from Allen Brain Atlas^33^ as a reference in our analysis (Supplementary Fig. 5a).

SpaSEG well characterized mouse brain structures such as cortical layers and hippocampus (particularly the subfields of CA1sp, CA3sp and DG-sg) at all resolutions (Fig. 2k). In contrast, Leiden mixed domains with other spots at Bin20 and Bin50 resolutions while failing to separate CA3sp and DG-sg at Bin100 and Bin200 resolutions. SpaGCN was unable to disentangle the Bin20 dataset due to the substantial number of spots and running out of memory. Neither did it yield continuous or neat spatial domains at any resolutions nor uncovered CA3sp and DG-sg separately at Bin50 and Bin200 resolutions. BayesSpace cannot perform spatial clustering on this large Stereo-seq data except at the low Bin200 resolution (Supplementary Fig. 6), partly because of the extremely large number of spots. The significantly low LISI values achieved by SpaSEG further demonstrated its superior performance in the identification of spatial domains concerning different resolutions (p<0.001; Fig. 2j). Most importantly, SpaSEG displayed exceedingly high computational efficiency together with strikingly little memory consumption for the large SRT dataset at high resolution, approximately 26-fold faster than SpaGCN over the Bin50 dataset (84,724 spots) and 2.5-fold than Leiden over the Bin20 dataset (526,716 spots) (Fig. 2l, Supplementary Fig. 7). To validate the SpaSEG-identified spatial domains, we investigated the spatial expressions of known marker genes for cortex layers and hippocampus in the Bin100 dataset (Fig. 2m). As expected, marker genes of *Cmss1* (domain 2), *Lamp5* (domain 11) and *Ttc9b* (domains 8, 13 and 19) were enriched in cortical layers while *Ncdn*, *Hpca*, *Neurod6* and *Spink8* were highly expressed in CA and DG-sg (domains 5, 10 and 15). Moreover, we found that marker genes of *Tcf7l2*, *Zic1* and *Prkcd* were uniquely expressed in domain 1 (thalamus, Supplementary Fig. 8). Collectively, these results showed that SpaSEG enabled the identification of spatial domains in large assayed tissue size at high resolution with considerable scalability.

### SpaSEG allows integrative analysis of SRT datasets from multiple adjacent tissue sections

To investigate the ability of SpaSEG to integrate SRT datasets, we first exploited two annotated adjacent sections of mouse olfactory bulb Stereo-seq datasets^11^ at bin14 resolution (Supplementary Table 1). We benchmarked SpaSEG with Harmony^34^ and LIGER^35^ in that they have been reported as the most optimal integration methods for scRNA-seq data^36^. Consequently, spatial domains identified and aligned simultaneously by SpaSEG from the two sections exhibited better consistency and correspondence than Harmony and LIGER (Fig.3a), making an advantage to reconstructing a stacked 3D tissue map (Fig. 3b). Notably, the SpaSEG-identified domains, together with the highest clustering accuracy (Fig.3c), also predominantly preserved the laminar structures in tissue sections such as GCL in domain 0 and SEZ in domain 1 (Fig. 3d). UMAP analysis demonstrated that, using the latent representations, SpaSEG tended to generate compact distinct domain-specific clusters that nearly evenly mixed with different tissue sections and laminar structures compared favorably to Harmony and LIGER (Fig.3e). This result suggested that SpaSEG had the potential to produce a biologically meaningful latent space that decoupled the intrinsically biological variations across adjacent sections from nuisance factors such as the batch effect ^36, 37^. Furthermore, SpaSEG achieved significantly higher scores of F1_*LISI*_ (0.889 ± 0.250) and F1_*SC*_ (0.651 ± 0.020) than other methods, suggesting its best efficacy in SRT data integration (p<0.0001, Fig.3f, Methods).

**Fig. 3.**
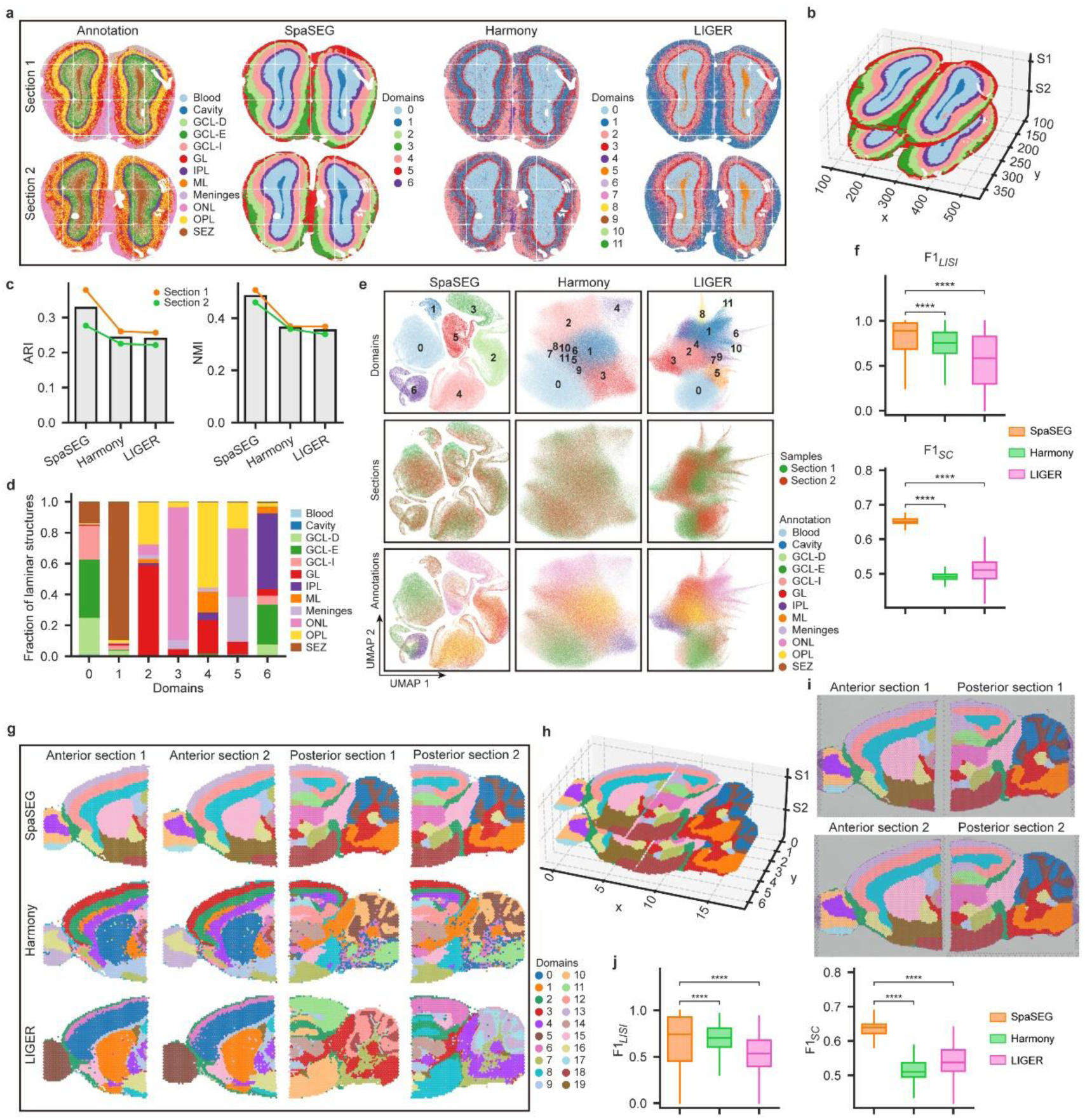
Integrative analysis of SRT datasets from multiple adjacent tissue sections. **a**, Annotations of two adjacent sections of mouse olfactory bulb Stereo-seq datasets at bin14 resolution (Left), Spatial domains in the two sections identified and aligned by SpaSEG (Second left), Harmony (Second right), and LIGER (Right). ONL: olfactory nerve layer, OPL: outer plexiform layer, GL: glomerular layer, GCL-D: granular cell layer deep, GCL-E: granular cell layer externa, GCL-I: granular cell layer internal, IPL: internal plexiform layer, ML: mitral layer, SEZ: subependymal zone. **b**, Visualization of 3D tissue structure map with SpaSEG-identified spatial domains. **c**, Results of ARI and NMI showing the spatial domain identification accuracy by SpaSEG, Harmony, and LIGER on the mouse olfactory bulb Stereo-seq datasets. Bar plots depict corresponding mean values. **d**, Faction of annotated laminar structures in each SpaSEG-identified spatial domain after data integration. **e**, UMAP plots illustrating qualitative evaluation of the SRT data integration by SpaSEG (Left column), Harmony (Middle column), and LIGER (Right column). Spatial domains (Top row), tissue sections (Middle row), and laminar structure annotations (Bottom row) were colored separately. **f**, Scores of *F*1_*LISI*_ (Top) and *F*1_*SC*_ (Bottom) showing the integration accuracy by SpaSEG, Harmony, and LIGER on the mouse olfactory bulb Stereo-seq datasets. **g**, Spatial domains identified by SpaSEG (Top row), Harmony (Middle row), and LIGER (Bottom row) for the four sections from the mouse brain 10x Visium datasets. **h**, Visualization of 3D tissue structure map after stitching anterior and posterior brain sections with SpaSEG-identified spatial domains. **i**, SpaSEG-identified spatial domains reflecting the shared layer structures across the anterior and posterior brain section. **j**, Scores of *F*1_*LISI*_ (Left) and *F*1_*SC*_ (Right) showing the integration accuracy by SpaSEG, Harmony, and LIGER on the mouse brain 10x Visium datasets. **** p<0.0001.

As a tissue sample could be larger than the assayed area used for transcriptomics, we further examine the capability of SpaSEG to stitch multiple tissue sections. To accomplish this, we used two adjacent sagittal sections of a mouse brain, each of which consisted of an anterior section and a posterior section (Supplementary Table 1). Results showed that SpaSEG yielded much neater spatial domains than Harmony and LIGER (Fig. 3g), which were in great agreement with the anatomic structures (Supplementary Fig. 5b). Moreover, UMAP analysis again showed that SpaSEG gained distinct domain-specific clusters and successfully merged the four sections (Supplementary Fig. 9). By stitching the corresponding anterior section and posterior section to make the spatial coordinates consecutive, SpaSEG-identified spatial domains reflected the shared structures between the two sections (Figs. 3h, i), along with the highest integration accuracy (p<0.0001; Fig.3j). We additionally performed the integrative analysis for the 12 human DLPFC sections. As expected, SpaSEG yielded excellent consistent spatial domains across multiple sections (Supplementary Figs. 10-12) and achieved the best performance on the SRT data integration (Supplementary Figs. 13). Interestingly, compared to the use of a single tissue section, integrating multiple adjacent sections appeared to improve the accuracy of spatial domain identification (Supplementary Fig. 14), which could be attributed to the efficient capture of gene expression variations across different tissue sections by SpaSEG.

### SpaSEG enables to detect spatially variable genes

Next, we employed the representative section 151673 in the human DLPFC 10x Visium dataset to examine the detection of spatially variable genes (SVGs). SpaSEG finally detected 81 SVGs, most of which (n=73, 90.12%) were specifically expressed in domain 0 (white matter) such as *CNP* while the remaining eight SVGs (9.88%) were in the domains of 2 (layer 2), 3 (layer 3) and 4 (layer 5) such as *PCAL1*, *NEFM* and *PCP4*, respectively (Fig. 4a, Supplementary Fig. 15). Only a few detected SVGs residing in neuronal layers suggested many non-spatial patterns of genes expressed in these regions. In contrast, benchmarking methods of SpaGCN, SpatialDE and SPARK detected a total of 65, 3661 and 3187 SVGs, respectively (Supplementary Fig. 16a). Nevertheless, SVGs detected by SpatialDE and SPARK did not necessarily show spatial patterns^18^. The significantly higher values of Moran’s *I* (0.364±0.140) and adjusted Geary’s *C* (0.358±0.140) achieved by SpaSEG than SpatialDE and SPARK suggested rather clear spatial patterns of the SpaSEG-detected SVGs (p<0.0001, Fig. 4b). Although SpaSEG and SpaGCN appeared no significant difference in the SVG detection on this small SRT dataset, SpaSEG was running the fastest (Supplementary Fig. 16b).

**Fig. 4.**
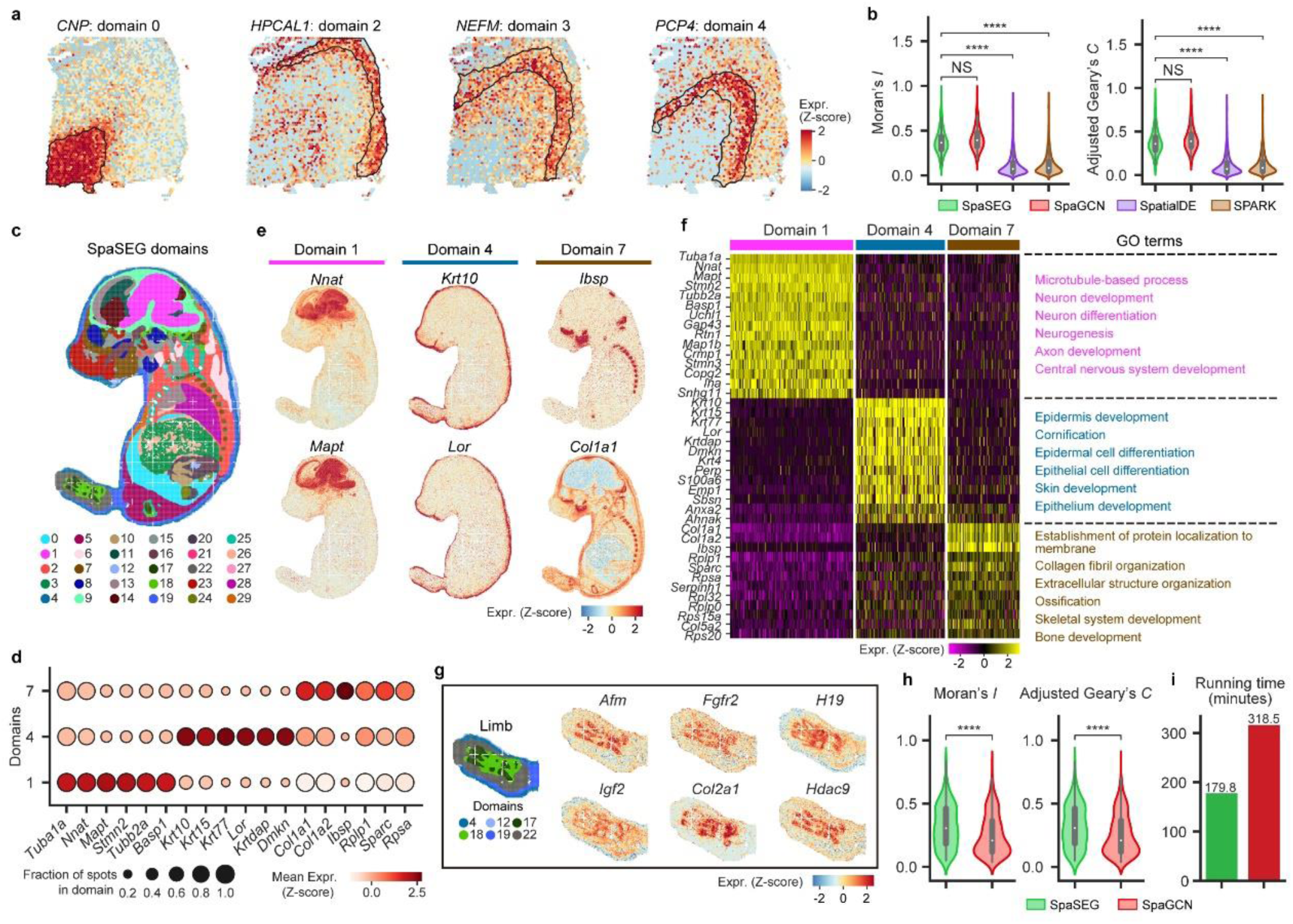
Domain-specific spatially variable genes detected by SpaSEG. **a**, Illustration of SVGs of *CNP*, *HPCAL1*, *NEFM*, and *PCP4* for domains 0, 2, 3, and 4, respectively, in 10x Visium DLPFC section 151673 data. **b**, Violin plots showing Moran’s *I* and adjusted Geary’s *C* values for SVGs detected by SpaSEG, SpaGCN, SpatialDE, and SPARK on the 10x Visium DLPFC section 151673. **c**, SpaSEG-identified spatial domain for the mouse embryo Stereo-seq data at Bin50 resolution. **d**, Dot plot showing expression of the top 6 prioritized SVGs in spots in each spatial domain. Color intensity indicates the average expression of the SVG in a domain while dot size represents the percentage of spots in that domain expressing the SVG. **e**, SVGs for domain 1 (*Nnat* and *Mapt*), domain 2 (*Krt10* and *Lor*), and domain 7 (*Ibsp* and *Col1a1*). **f**, Heatmap showing the top 15 (at most) SpaSEG-detected SVGs differentially expressed in domains 1, 4, and 7. Selected significant GO terms related to each spatial domain were highlighted. **g**, Visualization of spatial domains for mouse limb and the representative SVGs. **h**, Violin plots showing Moran’s *I* and adjusted Geary’s *C* values for SVGs in the mouse embryo Stereo-seq data detected by SpaSEG and SpaGCN. **i**, Running time for SVG detection through SpaSEG and SpaGCN. NS: not significant, **** p<0.0001.

Then, we assessed SVG detection using a large mouse embryo Stereo-seq dataset at Bin50 resolution with 72,944 spots. Based on 30 identified spatial domains (Fig.4c), SpaSEG detected a total of 252 SVGs. Of particular interest were domain 1 (brain), domain 4 (epidermis), and domain 7 (cartilage primordium/bone), which corresponded to 48, 13 and 12 SVGs, respectively (Supplementary Figs. 17-19). Most of these SVGs showed transcriptionally distinct spatial patterns and strong expressions in spots within the specific tissue domains and organs, making them useful to discern tissue regions as markers (Fig. 4d). For example, *Nnat* and *Mapt* were enriched in domain 1, *Krt10* and *Lor* in domain 4, while *Ibsp* and *Col1a1* in domain 7 (Fig. 4e). GO enrichment analysis showed that these SVGs exhibited features involved in organ-specific biological processes such as central nervous system development (domain 1), epidermis development (domain 4), and skeletal system development (domain 7) (Fig. 4f). We additionally analyzed the limb of the mouse embryo (Fig. 4g), leading to a total of 23 SpaSEG-detected SVGs (Supplementary Fig. 20). Most of them enriched in this region were implicated in limb development such as limb bud cell formation, proliferation and differentiation (*Afm, Fgfr2, H19* and *Igf2*), as well as limb skeletal development and the formation and maintenance of cartilage and bone in the limb (*Col2a1* and *Hdac9*) (Fig. 4g). For comparison, we applied SpaGCN on this Stereo-seq dataset, resulting in 458 SVGs (Supplementary Fig. 21). However, in addition to significantly higher values of Moran’s *I* and adjusted Geary’s *C* (p<0.0001, Fig. 4h), SpaSEG was nearly 1.8-fold faster than SpaGCN (Fig. 4i), suggesting its great effectiveness and efficiency in detecting SVGs with clear spatial patterns in the large SRT dataset.

### SpaSEG facilitates the analysis of ligand-receptor interactions

To investigate intercellular interactions throughout the entire tissue section, we proposed a method to analyze ligand-receptor interaction by incorporating spatial domains with the gene expression of ligands and their cognate receptors (Fig. 5a, Methods). Briefly, we first curated a list of significant ligand-receptor pairs (FDR-adjusted p value <0.05) from CellPhoneDB^38^ that exhibited putative inter- and intra-domain interactions. Spots within the corresponding domain can thus be implicitly linked to the curated ligand-receptor pairs. Cell types were deconvolved using cell2location^39^ to estimate cell abundances in each spot, where the cell type heterogeneity was measured using entropy. Finally, the LR score for a given spot was derived by leveraging the spot’s heterogeneity and the co-expression of the ligand-receptor pair among the spot and its neighbors.

**Fig. 5.**
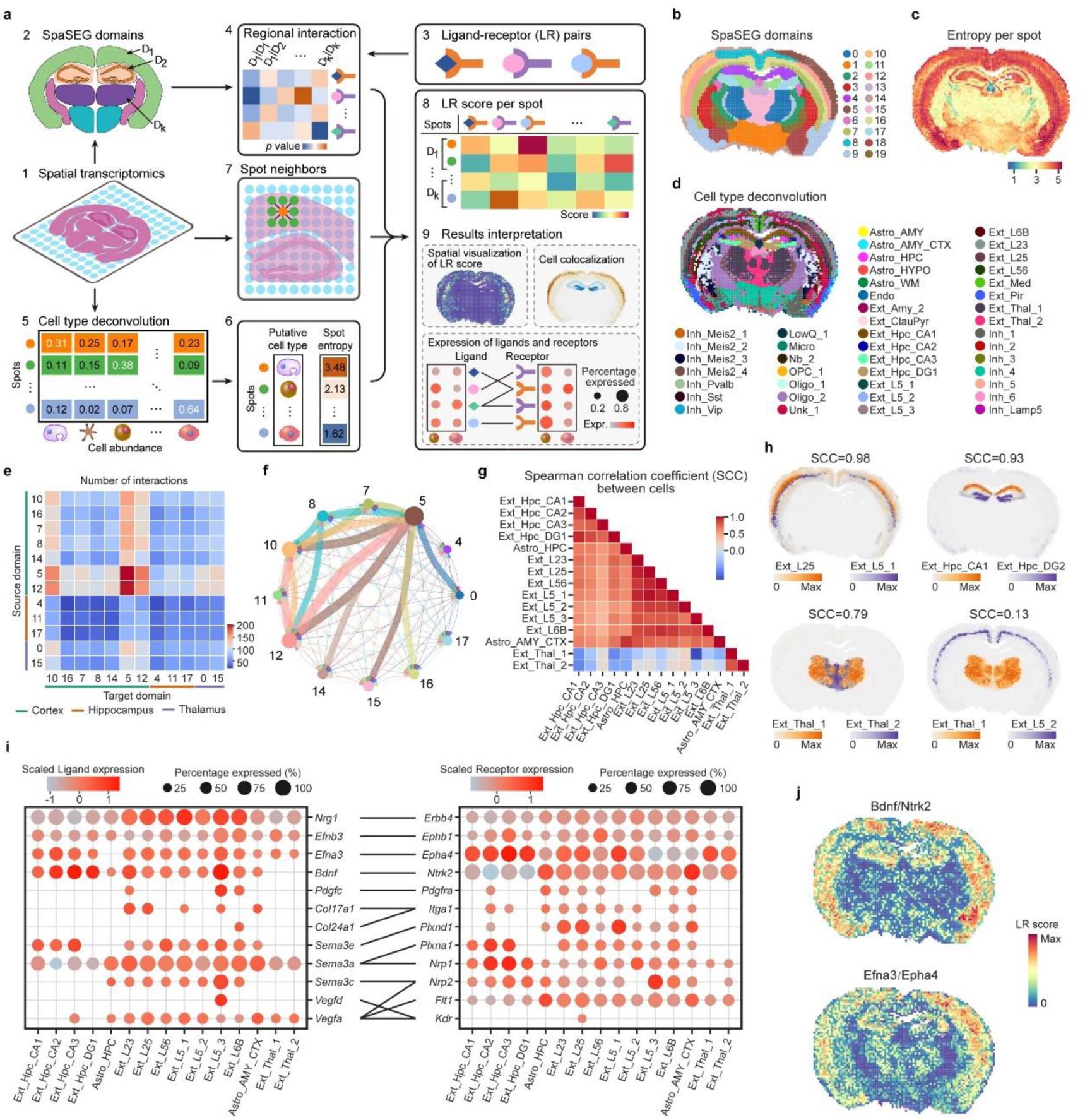
Spatial cell-cell interaction inference through SpaSEG based on gene expressions of ligand and receptor. **a**, Workflow of cell-cell interaction inference. including spatial domains identification (steps 1 and 2), ligand-receptor pairs curation (steps 3 and 4), cell type deconvolution and entropy calculation (steps 5 and 6), spot neighbor determination (step 7), LR score calculation and result interpretation (steps 8 and 9). **b**, SpaSEG-identified spatial domains for the adult mouse brain Stereo-seq data (Bin200). **c**, Entropy score showing cell type heterogeneity per spot. **d**, Cell type deconvolution by cell2location. Inh: inhibitory, LowQ: low-quality cell, Nb: neuroblasts, OPC: oligodendrocyte precursor cell, Unk: unknown cell, AMY: amygdala, HYPO: hypothalamus, WM: white matter, Endo: endothelial, Thal: thalamus, Ext: excitatory. **e**, Interaction heatmap plotting the total number of interactions between the source (y axis) and target (x axis) domains in the three mouse brain regions. **f**, Chord plot depicting interactions between spatial domains. Size of the colored domain node represents the number of interactions from and to that domain, while edge is colored the same as the from node and its thickness denotes the number of interactions from that domain (at least 90% quantile of the distribution of the interactions are shown). **g**, Spearman correlation coefficient (SCC) between cell types in the three regions. **h**, Spatial distribution of two cell-type abundances with SCC score. **i**, Dot plots showing average normalized expression of selected ligands (left) and receptors (right) in cell types in the three brain regions. Only genes from cell types with expression (>25%) are shown. Color intensity represents the average expression in the cell population while dot size represents the percentage of cells in that population expressing the gene. **j**, Visualization of spot-wise LR score within the adult mouse brain.

We applied our ligand-receptor interaction analytic method on the whole adult mouse brain Stereo-seq data at bin200 resolution with 20 SpaSEG-identified spatial domains (Fig. 5b). Analysis of cell-type deconvolution unraveled a greater cell-type heterogeneity with diverse subtypes of cortical neurons in the cerebral cortex (domains 5, 7, 8, 9, 10, 12 and 16) than other regions (Figs. 5c, d). In the hippocampus region (domains 4, 11 and 17), however, spots were predominated by excitatory neurons such as Ext_Hpc_CA1, Ext_Hpc_CA2, Ext_Hpc_CA3, and Ext_Hpc_DG1 (Supplementary Fig. 22). A total of 703 unique significant ligand-receptor pairs were detected among all spatial domains (Supplementary Fig. 23), most of which (n=576) occurred in the regions of cortex, hippocampus, and thalamus (Fig. 5e). Chord plot revealed domains of 5, 8, 10, and 12 in cortex area as central regional-interaction hubs in cortex region (Fig. 5f). Analysis of Spearman correlation coefficient (SCC) uncovered high colocalization of excitatory neurons within and between areas of cortex and hippocampus, as well as within thalamus (Figs. 5g, h, Supplementary Fig. 24). However, the low SCC of Ext_Thal_1 in the thalamus and Ext_L5_2 in the cerebral cortex may suggest their loose colocalization in these two regions. Analysis of highly expressed ligands and receptors revealed multiple signaling pathways implicated in mouse brain development (Fig. 5i), while spot-wise LR score illustrated a spatial interaction signature map (Fig. 5j, Supplementary Fig. 25). For example, the interaction of Bdnf and Ntrk2, strongly expressed in excitatory neurons and involved in synaptic plasticity and a wide range of cognitive processes^40^, was highly enriched in cortex and hippocampus. Signaling pathway of Efna3/Epha4, mainly expressed in astrocytes and neurons in the cerebral cortex, hippocampus and thalamus, have been implicated in the regulation of axon guidance, synaptic plasticity, and the formation and maintenance of neuronal circuits^41^. Together, these results implicated diverse cell crosstalk in the maintenance of adult mouse brain homeostasis and suggested the brain’s structural ligand-receptor interactions.

### Application of SpaSEG on a breast cancer sample with invasive ductal carcinoma

Finally, we applied SpaSEG on a published 10x Visium SRT dataset for an ER+/PR- /HER2+ breast cancer sample with invasive ductal carcinoma^17^ (IDC) that has been annotated by the pathologist (Fig. 6a). SpaSEG accurately dissected regions of invasive carcinoma (IC, domains 0, 2, 3, 8 and 9), in situ carcinoma (domain 6), benign hyperplasia (domain 4), and non-tumor region (domains 1, 5 and 7), which were largely in line with annotations (cluster purity=0.815, ARI=0.223, NMI=0.391, Fig. 6b, Supplementary Fig. 26). Results of cell type deconvolution showed predominant cancer cells in the invasive and in situ carcinoma regions, while other cell types were enriched in the rest where displayed great heterogeneity (Supplementary Fig. 27). SpaSEG detected a total of 159 domain-specific SVGs (Supplementary Fig. 28), with GO terms being mainly linked to tumor progression and immune response (Fig. 6c). Among these SVGs, oncogenes of *CCND1* and *MUC1*, for example, were highly expressed in domains 0, 2 and 8 while oncogene *ZNF703* in domains 3 and 9 (Fig. 6d). The IC region, including domains 0, 2, 3, 8 and 9, was enriched for genes *GATA3* and *XBP1* that were implicated in the development and progression of ER+/luminal cells ^42^. Non-invasive carcinoma region highly expressed antigen *HLA-A* that played a key role in the immune response^43^, and genes of *CD74*, *TIMP1* and *IFI27* that have been implicated in the tumor growth and proliferation and metastasis^44–46^.

**Fig. 6.**
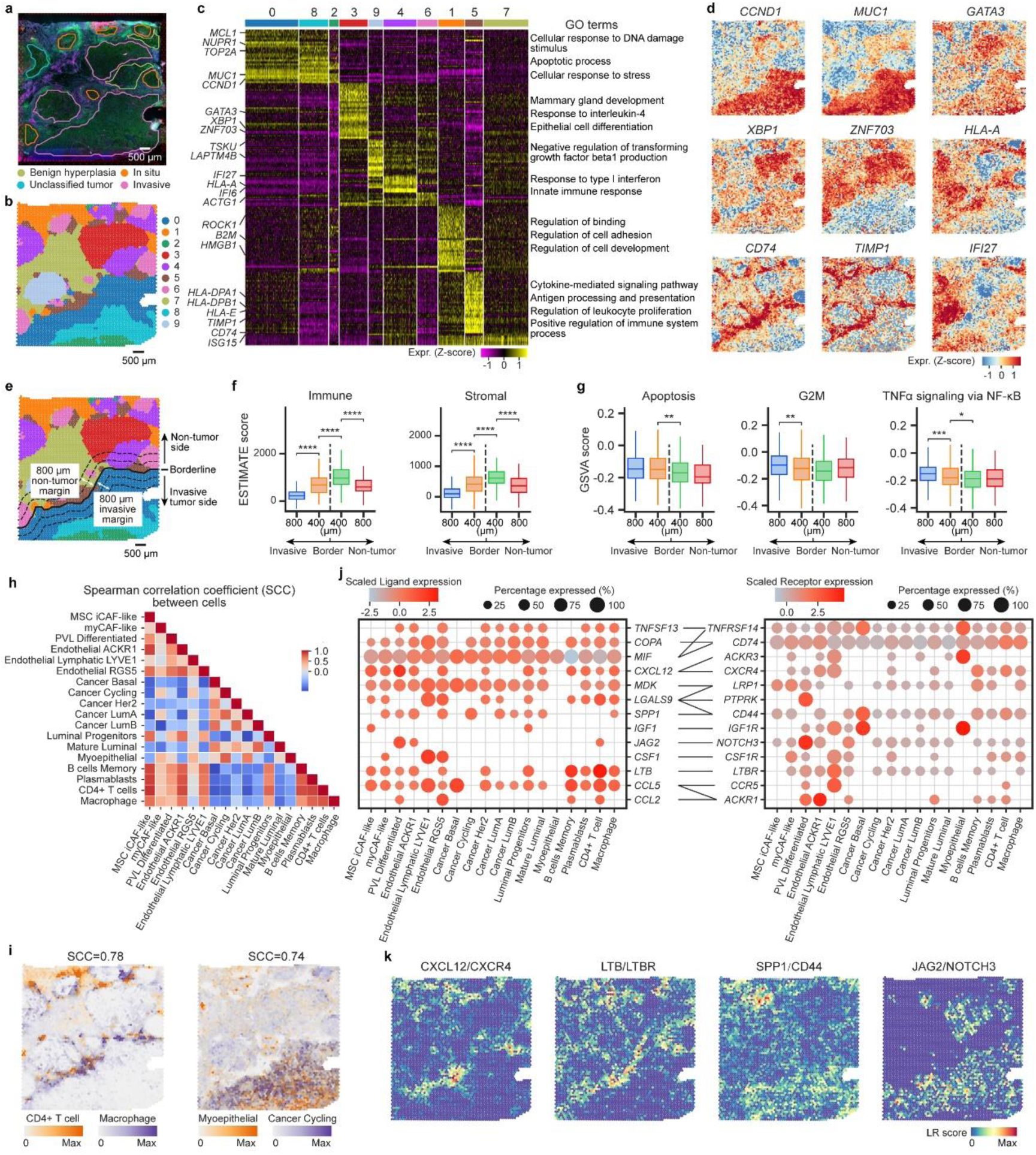
Application of SpaSEG on IDC dataset. **a**, Annotation of the raw IDC sample. **b**, SpaSEG-identified spatial domains. **c**, Heatmap showing the SpaSEG-detected SVGs differentially expressed in each spatial domain. Selected SVGs and corresponding significant GO terms related to each spatial domain are highlighted. **d**, Visualization of spatial patterns for representative SVGs. **e**, Borderline showing invasive tumor c and non-tumor regions. **f**, Boxplots showing immune and stromal ESTIMATE scores in invasive-tumor and non-tumor regions. **** p<0.0001. **g**, Boxplots showing GSVA scores of apoptosis, G2M checkpoint and TNFα signaling via NF-κB in invasive-tumor and non-tumor regions. * p<0.05, ** p<0.01, *** p<0.001. **h**, Spearman correlation coefficient (SCC) between cell types. CAFs: cancer-associated fibroblasts, MSC: mesenchymal stem cell, iCAF: inflammatory-like CAF, myCAF: myofibroblast-like CAF, PVL: perivascular-like. **i**, Spatial distribution of two cell-type abundances with SCC score. **j**, Dot plots showing average normalized expression of selected ligands (left) and receptors (right) in cell types. Only genes from cell types with expression (>25%) are shown. Color intensity represents the average expression in the cell population while dot size represents the percentage of cells in that population expressing the gene. **k**, Visualization of spot-wise LR score within the tissue.

To characterize the heterogeneity between the IC and the normal regions, we identified a border line that separated domains 0, 2 and 8 and the rest, and further partitioned four margin areas in 400 μm-width each, along the perpendicular extension direction against the borderline (Fig. 6e, Methods). Analysis of tumor-associated stromal and immune signatures using the ESTIMATE algorithm^47^ showed significantly high stromal and immune scores in the areas around the borderline (Fig. 6f), suggesting their identity as MSC iCAF-like cells, myCAF-like cells, macrophages and CD4+ T cells that were correspondingly enriched in these areas (Supplementary Fig. 29). GSVA analysis unraveled that IC region was significantly enriched for hallmarks of apoptosis and cell proliferation (G2M), as well as tumor necrosis factor (TNF)-α signaling via nuclear factor-κB (NF-κB) that can be activated by *XBP1* highly expressed in the same region to enhance tumor proliferation^48^, whereas these levels decreased significantly from IC region to paracancerous region and no significant difference was observed between areas in the normal margin (Fig. 6g). Moreover, hallmarks of glycolysis and hypoxia enriched in IC region suggested great metabolic changes in the tumor area (Supplementary Figures 30).

Analysis of the Spearman correlation coefficient (SCC) demonstrated the cell colocalization such as CD4+ T cell and macrophage (Figs. 6h, i). Analysis of ligand-receptor interaction revealed hundreds of significant ligand-receptor pairs (Supplementary Fig. 31a), and uncovered spatial domains of 0, 1, and 5 as central interaction hubs in the IDC sample (Supplementary Fig. 31b). Analysis of highly expressed ligands and their cognate receptors unraveled validated signaling pathways implicated in immunoregulation in breast cancer (Fig. 6j). For instance, cancer-associated fibroblasts (CAFs) may function in the immunoregulation via production of chemokines such as CXCL12, CCL5, and CCL2, interacting with immune cells, endothelial cells, and luminal progenitor cells^49^. The binding of cytokine LTB secreted by immune cells to LTBR expressed in lymphatic endothelial cells has been implicated in immune cell recruitment and tumor angiogenesis^50^. The interaction between differentiated perivascular-like (PVL) cells and cancer cells via JAG2/NOTCH3 signaling may be associated with tumor growth and proliferation^51^. Cancer cells interacting with immune cells through SPP1/CD44 signaling may serve roles in tumor progression, tumor angiogenesis and the formation of a pre-metastatic niche^52^. Moreover, analysis of the spot-wise LR score depicted the spatial interaction signature map within the IDC tissue (Fig. 6k, Supplementary Fig. 32). For example, interactions of CXCL12/CXCR4 and LTB/LTBR were enriched in paracancerous region and non-tumor region while JAG2/NOTCH3 and SPP1/CD44 were predominated in IC region. Collectively, these results demonstrated spatial tumor-immune dynamics in the tumor microenvironment of breast cancer with IDC.

## Discussion

The innovation of spatially resolved transcriptomics (SRT) technologies has dramatically advanced our knowledge of the functions of heterogenous cells and their interactions with the surroundings in a highly structured manner. SpaSEG is developed to cater to the growing needs of tertiary analysis tasks in this rapidly evolving field through a reliable and scalable model. It therefore, comparing favorably to existing state-of-the-art methods, provides an impressive computationally efficient approach to dissecting spatial domains in a collection of SRT datasets generated by a series of platforms at various resolutions and assayed tissue sizes. Based on spatial domains, SpaSEG is flexible to accommodate a range of downstream analysis tasks, including multiple adjacent sections integration, SVG detection, and spatial CCI inference.

Differing from the existing methods^37^, SpaSEG converts the spatial transcriptomics data into an image-like tensor, serving the spatial domain identification problem as the pixel-wise image segmentation problem in an unsupervised fashion. In each iteration, SpaSEG assigns a pseudo-label to each spot and clusters nearby spots with similar gene expression features into the same domain. To avoid overly fragmented domains, it penalizes sharp changes in the features between neighboring spots by minimizing the edge strength of the image-like tensor, which encourages a gradual transition between spatial domains. We have demonstrated that this regularization is essential to gain smooth and coherent spatial domains and thus improve spatial clustering performance (Supplementary Figs. 33 and 34a,b). The CNN-based architecture used by SpaSEG is comprised of canonical building components, making it computationally efficient, scalable and platform-agnostic, and thus suitable for analyzing various SRT datasets at different resolutions and tissue sizes (Fig. 2).

We have demonstrated the intrinsic utility of SpaSEG in integrating multiple adjacent tissue sections to facilitate uncovering important biological insights. Rather than stacking them together, we feed SpaSEG with the multiple SRT datasets and set the mini-batch size of the model parameter as the number of the SRT datasets during training, and thus learning inter- and intra-section patterns simultaneously. This implies the domain adaptation-based registration^53, 54^ for multiple SRT datasets by aligning spatial domains across distinct sections, making them consistent with and correspondent to the underlying anatomical structures (Fig. 3, Supplementary Figs. 10-12). Furthermore, registration of spatial domains would avert distortion and variation along the axis from adjacent sections, which enables SpaSEG to facilitate the reconstruction of an accurate 3D organization of tissue transcriptome atlas, and thus benefit revealing the tissue architecture comprehensively^55, 56^.

SpaSEG detects SVGs through comparing genes that are expressed significantly differentially across spatial domains, resulting in superior performance over SpatialDE, SPARK, and SpaGCN. The latter methods not only struggle with computational challenges but also suffer from suspectable spatial gene expression patterns and thereby would fail to fully reflect tissue-specific spatial functions, especially for large SRT datasets with many spots (Figs. 4h,i). Cell-cell interactions (CCIs) involve ligands and cognate receptors that contribute to transmitting signals between cells^3^. SpaSEG curates the known ligand-receptor pairs that interact among inter- and intra-domains to confer their spatial proximity^29^. SpaSEG explores putative CCIs within a tissue by aggregating gene expressions of ligand-receptor pairs in neighboring spots to derive spot-wise LR scores, providing a comprehensive view of the CCI map. Together with the consideration of cell type heterogeneity, multi-cell spots with high heterogeneity degrees and high co-expression of ligand-receptor in surroundings would have high LR scores, suggesting a great likelihood of related biological signaling events taking place between cells occupying the corresponding spots. Additionally, we quantify cell colocalization in a tissue using Spearman correlation coefficients (SCC). A positive high SCC between two cell types would indicate the colocalization with a high degree of overlap or correlation in their abundance profiles, suggesting that they would be likely to interact with each other, either directly or indirectly, within the tissue^6, 57, 58^.

Despite the impressive performance, we acknowledge that SpaSEG has serval limitations. First, SpaSEG now focuses on gene expressions and spatial coordinates due to the unavailability of other modalities such as H&E images from all existing SRT platforms currently, which is incompatible with the goal of SpaSEG as a platform-agnostic tool for SRT data analysis. However, SpaSEG in the future would integrate spatial multi-omics and multi-modal data that has provided invaluable complementary information for examining cellular biology and morphology^59^. Second, SRT often accompanies noise and plenty of zero gene expressions partly due to the diffusion of mRNA molecules^60^ and a relatively high dropout rate during the experiment. Further studies would be undertaken to explore the denoising and imputation of SRT data and thus enhance performance. Overall, we anticipate that SpaSEG could serve as a valuable SRT data analysis tool to deliver insights into the tissue-specific function, development, and pathology through elucidating tissue architectures with myriad cell types.

## Methods

### Data description

We apply SpaSEG on a collection of SRT datasets that are generated by a range of platforms at different resolutions, including seqFISH, MERFISH, Slide-seqV2, 10x Visium, and Stereo-Seq. The seqFISH and MERFISH provide single-cell spatial resolution but are limited to only capturing tens or thousands of genes, while the Slide-seqV2, 10x Visium, and Stereo-seq allow the transcriptome-wide detection of mRNAs. The Slide-seqV2 provides a near-cellular spatial resolution of 10 μm-diameter, whereas the 10x Visium offers a 6.5mm × 6.5mm capture area with a multicellular spatial resolution of 55μm in diameter. The Stereo-seq uses DNA nanoball patterned array chips to detect mRNA molecules typically with 1cm × 1cm capture area, where the data can be binned into various spatial resolutions, from subcellular to multicellular, depending on the bin size.

Specifically, the benchmark dataset of human DLPFC is generated by 10x Visium. For evaluating the reliability of SpaSEG on different SRT platforms, we exploit four datasets, including the mouse hemibrain Stereo-seq data that is an image-based cell segmentation dataset, the seqFISH data that is collected from the embryo 1 of sagittal sections of 8-12 somite-stage embryos (E8.5-E8.75), the mouse hypothalamic preoptic region MERFISH data (Bregma: -0.24mm), and the mouse hippocampus Slide-seqV2 data. To facilitate our analysis for different resolution levels using the raw whole adult mouse brain Stereo-seq data, we aggregate transcripts of the same gene into non-overlapping bin areas that cover corresponding DNA nanoball (DNB) sites. These bins are of sizes in 10μm diameter (Bin20; 20×20 DNB sites; equivalent to ∼1 medium mammal cell size), 25μm diameter (Bin50; 50×50 DNB sites), 50μm diameter (Bin100; 100×100 DNB sites), and 100μm diameter (Bin200; 200×200 DNB sites). To assess the ability of SpaSEG for the integrative analysis, we collect the mouse olfactory bulb Stereo-seq dataset at Bin14 resolution that contains two adjacent sections, and the mouse brain 10x Visium data that consists of adjacent sagittal anterior sections 1 and 2, and their corresponding sagittal posterior sections 1 and 2. In addition, we use the E16.5 mouse embryo Stereo-seq data at Bin50 resolution for the evaluation of SVG detection. We also collect the 10x Visium data for a breast cancer sample that is an estrogen receptor-positive (ER+), progesterone receptor-negative (PR-), human epidermal growth factor receptor (HER)2-amplified (HER2+) invasive ductal carcinoma (IDC). The number of spots and captured genes, as well as the information about the annotations for each of these SRT datasets are detailed in Supplementary Table 1.

### Data preprocessing

For each raw SRT dataset, we exclude genes expressed in less than five spots and spots expressing less than 5% quantile of gene count distribution. The retained gene counts per spot are further normalized using the library size, followed by the log-transformation. Principal component analysis (PCA) is then performed on the gene expression data in a *N* × *M* matrix with *N* spots and *M* genes, and prioritized top *d* PCs per spot are subsequently extracted. The optimal value of *d* varies from 15 to 100, depending on the SRT resolution and the analysis tasks (Supplementary Tables 4, 5). These PCs can explain the sufficient variability in the data and mitigate the computational intension, as well as yield the best spatial clustering performance (Supplementary Fig. 34c). Of note, the value of *d* is necessarily equal to or greater than the number of expected spatial domains. We recommend picking a greater value of *d* for the SRT data at a higher resolution. We further perform *z*-score normalization such that each PC has zero mean and unit variance. For the SRT data where spots are not necessarily arranged in a regular grid or lattice form such as that generated by seqFISH, MERFISH, Slide-seqV2, or Stereo-seq at the CellBin resolution, we scale down spatial coordinates to reduce the spot sparsity and thus narrow down the exceedingly large number of unwanted blank pixels when constructing the image-like tensor thereafter (Supplementary Table 4). For integrative analysis of multiple SRT datasets from adjacent tissue sections, SpaSEG undergoes the same preprocessing procedures except that we conduct PCA for dimensionality reduction over the multiple datasets based on the shared genes.

### Conversion of SRT data into image-like tensor

To enable SRT data analysis through SpaSEG, we convert the gene expression data with spatial information into an image-like tensor. Specifically, a spot *n* in the SRT array at row *i* and column *j* is represented by a PCA feature vector of 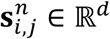, where *d* is the number of extracted PCs. Consequently, an *d*-channel image-like tensor 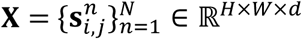 is created, where *N* is the number of spots, *H* and *W* are the height and width of the SRT array, *i* ∈ {1, 2, …, *H*}, *j* ∈ {1,2,, …, *W*}. For simplification, we denote the image-like tensor by 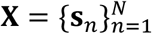 unless otherwise specification.

### SpaSEG model development

The architecture of SpaSEG starts with a batch normalization layer, followed by two convolutional blocks and a refinement module consecutively (Fig. 1b). Each convolutional block is composed of a 3 × 3 (filter size) convolutional layer with stride of 1 and padding of 1, a batch normalization layer, and a leaky ReLU activation layer with a fixed value of 0.2 for the intermediate parameter. The number of filters in the two convolutional blocks are *n*_1_ and *n*_2_, respectively. We set *n*_1_ and *n*_2_ equal to *d* in all experiments for simplification. The refinement module consists of a 1 × 1 convolutional layer with *d* filters, followed by a batch normalization layer. It is worth noting that the output shape of each convolutional layer remains the same as input.

Formally, given a SRT dataset that was represented by an image-like tensor 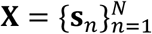, the latent feature representation for spot *n* can be learned by

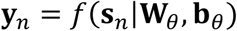

where *f*(·) is the SpaSEG network model with the trainable parameters ***W***_*θ*_ and *b*_*θ*_ that can be learned during the iterative training process, 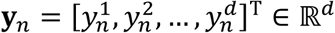. Then, the class pseudo-label for spot *n* can be obtained by

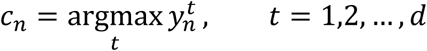

### Loss function

We regard the spatial domain identification as a spot-wise segmentation problem in an unsupervised fashion, where the class label for each spot can be viewed as a segment. To this end, we first consider the commonly used cross entropy loss with L2-norm regularization over the pseudo-labels as follows.

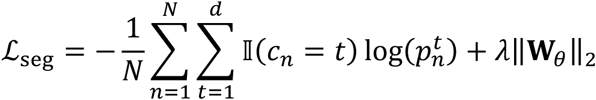

where 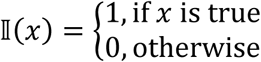, 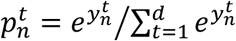, and *λ* is regularization parameter that controls the L2-norm regularization penalty and we empirically set it to 0.00001 in our experiments.

To encourage smoothness and coherence in the SRT segmentation results, we further consider a penalty term that measures edge strength of the image-like tensor. Inspired by the previous study^61^, we define the edge strength as the L1-norm of the gradient of image-like tensor to measure the horizontal and vertical changes in spot latent feature representations. The gradient of the SRT image-like tensor is calculated as:

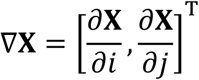

Then, the edge strength loss function can be given by:

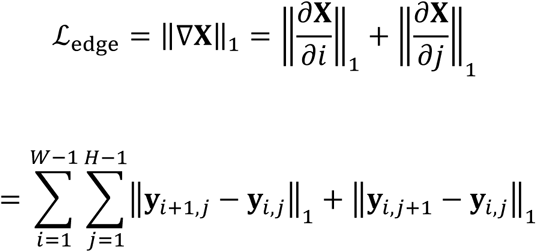

Consequently, the overall loss is then given by:

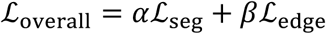

where *α* and *β* are weighting factors for segmentation and edge strength, respectively. The overall loss function is optimized iteratively during the segmentation process.

### SpaSEG training

SpaSEG is an end-to-end CNN-based model and trained de novo. Rather than randomly initializing the parameters, we first pre-train SpaSEG in the first 400 training epochs (Supplementary Fig. 34d) using the mean squared error loss defined as 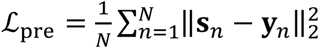 to initialize the model parameters. In the subsequent epochs, the latent feature representation ***y***_*n*_ and the corresponding pseudo-label *c*_*n*_ for each spot is obtained and the model parameters are updated by interactively calculating and backpropagating the overall loss ℒ_overall_. This process is repeated until the number of expected spatial domains is reached up to 2000 epochs during training. Otherwise, an additional post-process is performed in the following 100 epochs. More specifically, in each epoch we calculate pairwise Euclidian distances between centroids of the resulting spatial domains to measure their similarities. The spots within the two most similar domains are re-labeled as the same class. After updating the spot labels, the training process goes on. This post-process seeks to further reduce outliers in spatial domains and make them neater. The pseudo-code for training SpaSEG is listed in Supplementary Fig. 35.

We use Adam optimizer^62^ with the default parameters *β*_1_ = 0.9 and *β*_2_ = 0.999, and a mini batch size of *m*, where *m* is the number of SRT sections. We have searched various values of learning rate and set it to be 0.002 as an optimal one for all experiments (Supplementary Fig. 34e). The optimal values of weight factors *α* and *β* depend on analysis task. We suggest choosing *α* and *β* to be 0.4 and 0.7 for single-section SRT dataset (Supplementary Figs. 34a,b, Supplementary Table 4) while 0.2 and 0.4 for multi-section SRT datasets analysis, respectively (Supplementary Table 5), but there has variations for Stereo-seq data at different resolutions (Supplementary Table 4). In general, these values for SpaSEG hyper-parameters are determined in the way of combining grid search and manual tuning such that the optimal performance can be achieved.

### Spatial variable genes detection

After identifying spatial domains, SpaSEG first selects significant genes by performing differentially expressed genes analysis between the target domain and the rest of domains (termed as out-domains) using the Wilcoxon rank-sum test with the FDR-adjusted p value <0.05 by calling ‘scanpy.tl.rank_genes_groups()’ in the Scanpy package^63^ (v1.9.1). For the convenience of description, we additionally designate the domain with the highest ratio of spots expressing a given gene in the out-domains as max-out-domain.

Next, to facilitate the discovery of spatially variable genes (SVGs) with significant spatial patterns, we calculate following indices: (1) log2FC, defined as the log2 fold change of the gene expression between the target domain and the out-domains; (2) log2FC-in-max; referred to as the log2 fold change of the gene expression between the target domain and max-out-domain; (3) in-ratio, out-ratio, and max-out-ratio that are defined as the ratio of spots expressing the gene in the target domain, out-domains and max-out-domain, respectively; (4) in-CV and max-out-CV that are referred to as the coefficient of variation (CV) of the gene in the target domain and max-out-domain, respectively. The CV is used to quantify the degree of expression variation of the gene in the specific spatial domain, which can be calculated by 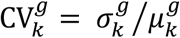, where 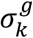 and 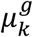 represent the standard deviation and the mean of expression of gene *g* in domain *k*. A gene with a low CV indicates its similar expression level in all spots within the specific domain.

Based on the selected significant genes, SpaSEG obtains candidate SVGs according to the following initial screening criteria: (1) both log2FC and log2FC-in-max >1.5; (2) in-ratio > 0.75; (3) in-ratio/out-ratio >1; and (4) in-ration/max-out-ratio >0.95. After that, the final list of SVGs are determined only if the candidates meet one of the following three conditions: (1) sufficiently expressed in the target domain, that is, in-ratio/out-ratio >1.2 or in-ratio/max-out-ratio >1.2; (2) highly differentially expressed in the target domain, that is, log2FC >8.0; or (3) both log2FC and log2FC-in-max >3.0, as well as in-CV <0.6 and in-CV/max-out-CV <1.0. The GO enrichment analysis is further performed on the final list of SVGs for each spatial domain using GSEApy^64^ (v1.0.4; FDR-adjusted p value <0.05).

### Cell type deconvolution

To spatially map the cell types annotated in single-cell or single-nuclei RNA sequencing (scRNA-seq or snRNA-seq) data to SRT data at multi-cell resolution, we employ cell2location^39^ (v0.8a0) to deconvolve multi-cell spots into cell-type abundances. For the whole the adult mouse brain Stereo-seq data at bin200 resolution, we use a mouse brain snRNA-seq data as reference that was published accompanying the cell2location paper. This raw mouse brain snRNA-seq data contained 40,532 annotated cells of totally 59 distinct cell types with 31,053 genes. For the IDC breast cancer 10x Visium data, a publicly available breast cancer scRNA-seq dataset^65^ is utilized as reference. Given that the IDC sample was characterized by status of estrogen receptor-positive (ER+), progesterone receptor-negative (PR−), and human epidermal growth factor receptor (HER) 2-amplified (HER+), we only select single cells from the all eleven ER+ and five Her2+ patients. Consequently, the raw breast cancer scRNA-seq data include 57,552 cells with totally 29 annotated cell types and 29,733 genes. After excluding genes expressed less than five cells or 3% of total cells from the reference single-cell datasets (resulting in 12,949 and 13,644 genes for the snRNA-seq and scRNA-seq data, respectively), we run cell2location with default parameter values, except ‘max_epochs’= 20,000, ‘batch_size’= 4096, ‘N_cells_per_location’=30, and ‘detection_alpha’=200.

### Ligand-receptor interaction analysis

To infer cell-cell interactions with the tissue section (Fig. 5a), we first identify spatial domains using SpaSEG. Next, we utilize Squidpy^66^ (v1.2.3) to curate a list of significant known ligand-receptor pairs between source and target domains (permutation test with 1000 permutations, FDR-adjusted p value <0.05). The Squidpy re-implemented a fast CellphoneDB^38^ analysis, supplemented additionally with the Omnipath^67^ database for ligand-receptor annotations. As a result, spots belonging to a specific spatial domain were implicitly linked to the corresponding ligand-receptor pairs. In parallel, we deconvolve the multi-cell spots into cell-type abundances using the cell2location algorithm. The cell-type abundances represented the proportion of the cell types defined in single-cell data residing in a multi-cell spot. Thus, the putative target cell type for a spot can be simply determined by the one with the maximal abundance. Besides, the cell-type abundance profiles also suggest the heterogeneity of cell types in a multi-cell spot, which can be quantified by the entropy defined as follows:

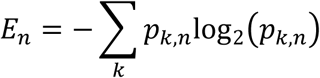

where *p*_*k*,*n*_ represents the proportion of cell type *k* in spot *n*, which is obtained by normalizing the cell-type abundances per spot into the range [0, 1]. A high entropy score in a multi-cell spot reflects a great degree of cell type heterogeneity.

To derive the LR score, we first define the co-expression signature of a ligand *L* in spot *i* and a receptor *R* expressed in spot *j* using the geometric mean of the expression *L* and *R* as follows:

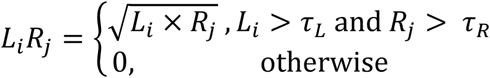

where *τ*_*L*_ and *τ*_*R*_ represent the minimum threshold of gene expression levels of *L* and *R*, which are set to 0 in our experiments. Then, we consider two rings of spots around a center spot *n* as its spatial neighbors by calling ‘squidpy.gr.spatial_neighbors()’ function in the Squidpy package. Thus, the LR score for the spot *n* is given by:

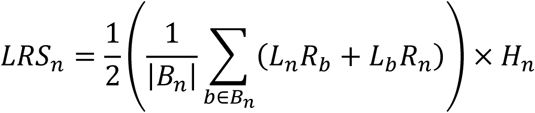

where *B*_*n*_ is the set of spatial neighbors of spot *n* while |*B*_*n*_| represents the number of the neighbors. *H*_*n*_ denotes the heterogeneity of the spot *n*, which is 1 for the single-cell spot or the value of entropy *E*_*n*_ for the multi-cell spot.

### Cell type colocalization

We adopt cell-type abundances to calculate spot-wise Spearman correlation coefficient (SCC) between cell types, which was used to assess their colocalization within the tissue. Suppose *A*_*k*_ is the abundance vector of cell type *k* in all spots within a tissue, the SCC between two cell types *k* and *m* is then defined as follows:

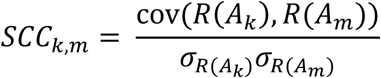

where *R*(*A*_*k*_) and *R*(*A*_*m*_) represents the rank abundances of cell types *k* and *m*, cov(*R*(*A*_*k*_), *R*(*A*_*m*_)) is their covariance while *σ*_*R*(*Ak*)_ and *σ*_*R*(*Am*)_ are their standard deviation. The SCC value ranges from -1 to 1 and a high positive value suggests high colocalization between two cell types.

### Tumor-normal borderline identification

With the SpaSEG-identified spatial domains in the IDC section, of particular interest are the domains 0, 2, and 8 that have been annotated as the invasive tumor. To facilitate the identification of a border between this invasive tumor region and the remaining regions, we convert the IDC section with spatial domains into a binary image, where a pixel corresponds to a spot and the spots not in the domains 0, 2 and 8 are masked.

We use Python OpenCV (opencv-python; v4.7.0) to first detect Canny edge based on the manually tailored threshold and the Gaussian blurring operation such that many unexpected small areas can be avoided. We further perform the dilation operation on the edged image and finally find the contours in the dilated image. We select the contour with the largest area and further smooth and approximate its shape by calling functions of ‘arcLength()’ and ‘approxPolyDP()’, leading to the final border line. Next, we further create parallel curve lines that perpendicularly extend 400μm and800 μm to both invasive tumor side and normal tissue side using ‘LineString.buffer()’ function in the shapely Python package (v2.0.1; https://shapely.readthedocs.io/en/stable/), leading to four margins that cover regions from the 800 μm-line to the 400 μm-line and from the 400 μm-line to the border line on both sides. We exclude spots on the normal tissue side that have been originally identified in the non-normal spatial domains, ensuring that the margins on that normal tissue side do not contain any tumor spots.

### Analysis of cell type composition, and immune, stromal and tumor hallmark score in the tumor and normal tissue margin

We compare the enrichment of cell types derived by cell2location among the four margins that are determined by the border line between invasive tumor side and normal tissue side. Based on the normalized gene expression profiles, we use ESTIMATE algorithm^47^ (v1.0.13) to calculate immune and stromal scores for each of the four margins. Besides, tumor hallmark pathways exported using the Molecular Signature Database (MSigDB) are also analyzed for each margin using the Gene Set Variation Analysis^68^ (GSVA; v1.46.0).

### Evaluation metrics

Apart from qualitative visualization, the performance of SpaSEG on SRT data analysis is also evaluated using the following quantitative metrics:

(1) Adjusted rand index (ARI)

ARI score is widely used to measure the similarity between predicted clusters and the reference labels, which is defined as:

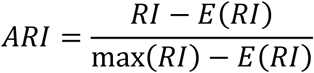

where *E*(*RI*) and max(*RI*) are the expectation and maximum of rand index (RI) distribution, 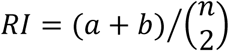, *a* is the number of pairs with the same true labels being correctly assigned to the same clusters, *b* is the number of pairs with different true labels being correctly assigned to different clusters, 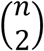 is the total number of unordered pairs in the data. By adjusting RI to ensure a value close to 0 for random labeling, the ARI ranges from -1 to 1 with a high positive score suggesting the high consistency between predicted clustering results and true label distribution. We calculate ARI score by calling ‘sklearn.metrics.adjusted_rand_score()’ in scikit-learn Python package (v1.2.2).

(2) Normalized mutual information (NMI)

NMI score is another well-known measure of the similarity between predicted clusters and the reference clusters. The NMI rescales mutual information (MI) to have a limit from 0 to 1, which is defined as:

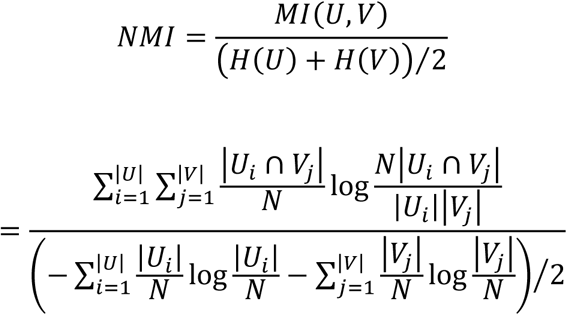

where *U* and *V* are reference cluster set and predicted cluster set, *H*(*U*) and *H*(*V*) are entropy values of clusters in the sets of *U* and *V*, |*U*| and |*V*| are the number of clusters in the sets of *U* and *V*, *N* is the total number of spots in the data, |*U*_*i*_| and |*V*_*j*_| are the number of spots in the true cluster *U*_*i*_ and predicted cluster *V*_*j*_. The MI measures the amount of information about true labels in the predicted clusters while NMI trade off quality of clustering against number of clusters. A high NMI score indicates high agreement of the predicted labels to its true clusters. The function of ‘sklearn.metrics.normalized_mutual_info_score()’ in scikit-learn is used to calculate NMI.

(3) Local inverse Simpson’s index (LISI)

To assess clustering performance without ground-truth annotation, we employ LISI^34^ score to measure the spots grouped into their own domains (i.e., domain LISI, denoted as dLISI). The dLISI is computed using the selected nearest neighbors based on local distance distribution with a fix perplexity with ranges greater than 1. A dLISI value close to 1 indicates the high purity of predicted spot labels in clusters. We use Python version (https://github.com/slowkow/harmonypy/blob/master/harmonypy/lisi.py) to calculate the dLISI for each spot in the data.

(4) *F*1_*LISI*_ and *F*1_*SC*_

In the analysis of multiple adjacent tissue sections, we have two facets to assess the integration performance. One is to measure how well the different sections are registered and aligned such that spots predicted in the same cluster from different sections should be mixed together. Another is to measure how well the spots are grouped into their own cluster. To this end, we calculate both section LISI (denoted as sLISI) and dLISI. A sLISI score close to the number of sections suggests the good registration and alignment of multiple sections. To combine the assessments of section mixing and spot purity, we normalize the sLISI and dLISI to [0, 1] and then compute a harmonic mean (F1 score) as follows:

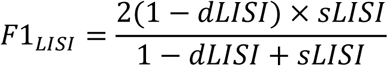

In addition, the Silhouette coefficient (SC) can also be used to measure how well a spot lies in its own cluster in comparison to other clusters, which is defined as:

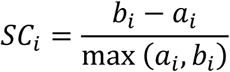

where *a*_*i*_ is the average distance between spot *i* and other spots in its cluster, *b*_*i*_ is the average distance between spot *i* and the spots in its nearest cluster. The SC value is between -1 and 1. A positive high SC of a spot suggests a more closeness of the spot to its own cluster but discrepancy to others. We call ‘sklearn.metrics.silhouette_samples()’ function in scikit-learn to compute section SC (sSC) and domain SC (dSC) for each spot, and further scale sSC and dSC into the range of [0,1]. Finally, we incorporate these two scores by calculating their harmonic mean:

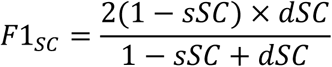

High *F*1_*LISI*_ and *F*1_*SC*_ suggest good multi-section integration.

(5) Moran’s *I* and adjusted Geary’s *C*

We employ two statistics of Moran’s I and Geary’s C to assess spatial expression patterns of the detected SVGs. Both Moran’s *I* and Geary’s *C* are correlation coefficients to measure the overall spatial autocorrelation of the data. They can be used to quantify the degree of gene expression similarity between one spot and its neighbors.

The Moran’s *I* score is calculated by:

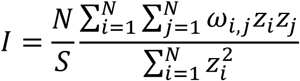

where *N* is the total number of spots in the data, *ω*_*i*,*j*_ is the spatial weight between spot *i* and *j*, *z*_*i*_ is the deviation for the gene expression in spot *i* from its mean (*x*_*i*_ − *x̄*), *S* is the aggregate of all spatial weights 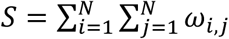.

The Geary’s *C* is given by:

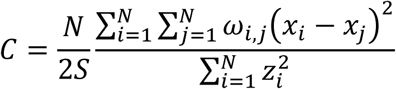

For Moran’s *I*, the cross-product is based on the deviations from the mean for the gene expression in two spots while for Geary’s *C*, the cross-product used the actual gene expression at each spot. We use ‘squidpy.gr.spatial_autocorr()’ function implemented in Squidpy package with parameters of ‘mode’ being set as ‘moran’ and ‘geary’ for Moran’s *I* and Geary’s *C*, respectively.

For the convenient explanation, we adjusted the Geary’s *C* score using *C*_*adj*_ = 1 − *C*. As a result, the ranges of the Moran’s *I* and the adjusted Geary’s *C* are both from -1 to 1. A higher positive score suggests a higher spatial autocorrelation, and thus, a clearer spatial expression pattern.

### Visualization and Statistics

Each box in the Boxplots or Violin plots corresponds to interval between 25th and 75^th^ percentile (interquartile range, IQR) with center line indicating median and whiskers = 1.5 × IQR. The two-side Mann-Whitney U test with Benjamini-Hochberg correction is used for statistical test. P values of <0.05 are considered statistically significant for all analyses. Error bars in plots represents mean ± standard deviation. All values are presented as median ± standard deviation unless specified otherwise.

### Experiment setting

SpaSEG is implemented using Pytorch (v1.12.0) and Python (v3.9). All experiments are carried out on a workstation with one NVIDIA Tesla P100 GPU addressing 16 GB RAM, and two 24-core Intel Xeon Gold 5118 CPUs addressing 187 GB RAM.

### Competing methods

We compared SpaSEG with Leiden, stLearn, Giotto, SpaGCN, BayesSpace, and SEDR for spatial domain identification, Harmony and LIGER for multiple adjacent sections integration, and SpatialDE, SPARK and SpaGCN for SVG detection. Description of all these methods and their parameter settings are detailed in Supplementary Information.

### Data availability

All datasets analyzed in this paper are publicly available. The web links to all data sources are provided in Supplementary Table 1. For reference datasets to deconvolve cell types, mouse brain snRNA-seq data is available at https://cell2location.cellgeni.sanger.ac.uk/mouse-brain, whereas the breast cancer scRNA-seq dataset is collected from https://singlecell.broadinstitute.org/single_cell/study/SCP1039/a-single-cell-and-spatially-resolved-atlas-of-human-breast-cancers.

### Code availability

The code for development and analysis of SpaSEG is available at:https://github.com/y-bai/SpaSEG.

## Acknowledgements

We would like to thank Dr. Lei Han, Dr. Liang Wu, and Dr. Yiwei Lai at BGI-Shenzhen for constructive discussions.

## Author contributions

Conceptualization and study design: Yong Bai

Supervision: Yong Bai, Xin Jin, Xun Xu, Ao Chen, Liang Wu, Yong Zhang

Data collection: Xiangyu Guo, Keyin Liu

Model development: Yong Bai, Xiangyu Guo, Keyin Liu

Model comparison: Xiangyu Guo, Yingyue Wang, Qiuhong Luo

Result analysis and interpretation: Yong Bai, Keyin Liu, Bingjie Zheng,

Visualization: Yong Bai, Xiangyu Guo, Keyin Liu

Manuscript draft: Yong Bai, Xiangyu Guo, Keyin Liu

All authors reviewed and edited the manuscript.

## Supplementary Information

### Competing methods and parameter settings

We benchmarked SpaSEG with a couple of state-of-the-art methods for various SRT analysis tasks, including:

- Leiden is a well-known community detection algorithm that has been broadly used for the clustering task in scRNA-seq data analysis. We used the same data preprocessing procedure as SpaSEG and ran the ‘scanpy.tl.leiden()’ function in the Scanpy package with the optimal resolution parameter that was manually tailored to yield the annotated cluster number or the expected cluster number in case of no available annotation for the data (and hereafter).stLearn is designed for SRT data analysis by incorporating spatial transcriptomics profiles and associated H&E pathology image morphology information. Following the official tutorial (https://stlearn.readthedocs.io/en/latest/tutorials.html), we performed the clustering task using stLearn (v0.3.2) with default parameters except for setting the number of principal components to 15.
- Giotto is developed for SRT data analysis based on the hidden Markov random field (HMRF) model. According to the online tutorial (https://giottosuite.readthedocs.io/en/master/), we ran Giotto (v1.0.4) for spatial clustering using default parameters except filtering highly variable genes (HVGs) with ‘perc_cells>3’ and ‘mean_expr_det>0.4’, as well as using KNN method with ‘k=5’ to create a spatial network. The number of HMRF domains was set to be equal to the number of ground truth clusters or the expected number of clusters.
- SEDR is developed to learn low-dimensional latent embeddings of gene expression with spatial location information through a deep autoencoder and a variational graph autoencoder. We ran SEDR that was downloaded from the Github repository (https://github.com/JinmiaoChenLab/SEDR) with default parameter settings and performed Leiden clustering with the optimal resolution parameter.
- SpaGCN allows spatial domain identification and SVG detection for SRT data by developing a graph convolutional network that jointly considered gene expression, spatial location and histology information. We followed the tutorial (v1.2.0; https://github.com/jianhuupenn/SpaGCN) to perform spatial domain identification and SVG detection with default parameters except setting ‘min_cell=3’ for gene filtering, ‘alpha=1’ and ‘beta=49’ for adjacent matrix calculation, as well as setting histology to ‘True’ in the case histology image was available. We searched the hyperparameter ‘*l*’ from 100 to 500 with step equal to 1 based on default ‘p=0.5’, and finally conduct Leiden clustering. We trained SpaGCN using default values of learning rate equal to 0.05 and max training epoch equal to 200.
- BayesSpace performs spatial clustering analysis based on a Bayesian statistical method that uses a t-distributed error model. We ran BayesSpace (v1.2.1) following the Github tutorial (https://github.com/edward130603/BayesSpace) with default parameters. By default, the top 2000 highly variable genes were selected to perform PCA and we set ‘q’ to the number of PCA, ‘d’ to the desirable cluster number, ‘nrep’ to 50000, and ‘gamma’ to 3.
- Harmony is a popular algorithm for integrating multiple scRNA-seq datasets. We used the same principal components as SpaSEG for the Harmony algorithm. We employed harmonypy (v0.06; https://github.com/slowkow/harmonypy) which was also warped by the ‘scanpy.pp.harmony_integrate()’ function in the external Scanpy package. Default parameters were used and Leiden was adopted to conduct clustering with the optimal resolution parameter.
- LIGER integrates multiple scRNA-seq datasets using nonnegative matrix factorization to reduce the dimension of SRT data. The pyliger (https://github.com/welch-lab/pyliger) was mainly used in our experiments. We conducted the same procedure of filtering genes and spots as SpaSEG and performed integrative analysis following the online tutorial with default parameter and used Leiden with the optimal resolution parameter to perform clustering.
- SpatialDE and SPARK are methods to identify genes with spatial expression patterns (SVGs) using Gaussian process regression and generalized spatial linear model, respectively. Both approaches were applied in our experiment with default parameters following the online tutorials (https://github.com/Teichlab/SpatialDE and https://github.com/xzhoulab/SPARK).

---

**Supplementary Fig. 1.**
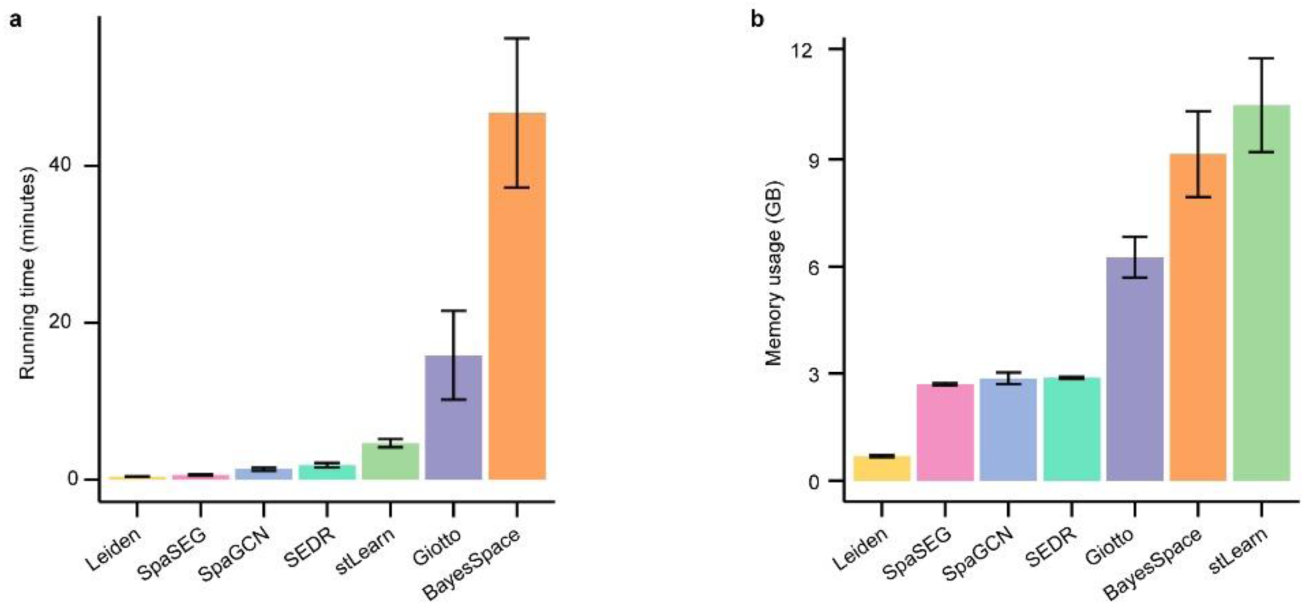
Computational cost for identifying spatial domains in all 12 tissue sections in the 10x Visium DLPFC dataset using SpaSEG and the competing methods. **a**, Running time analysis (in minutes)**. b,** Memory usage analysis (in GB). Error bars: mean ± standard deviation.

**Supplementary Fig. 2.**
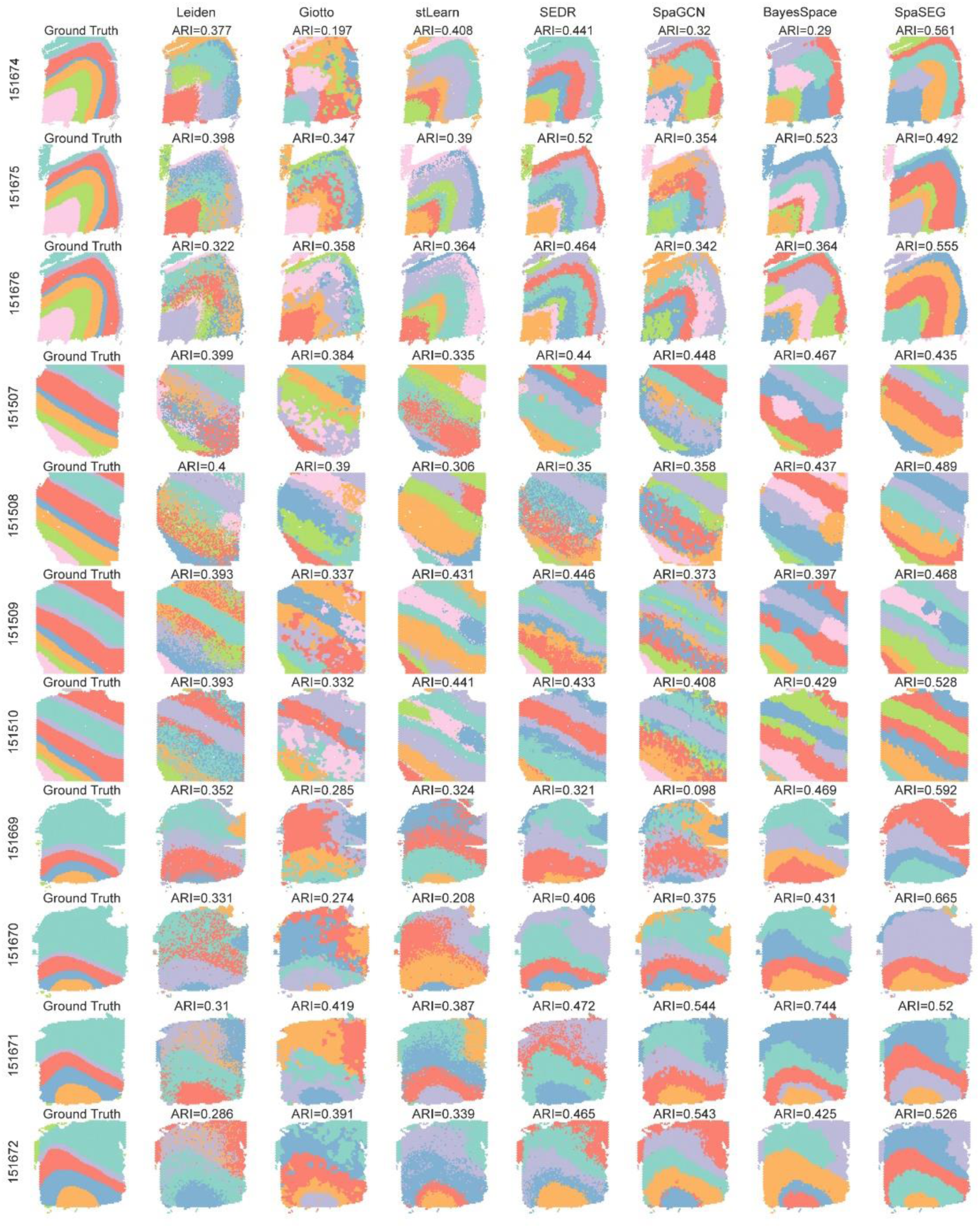
Visualization of spatial domains in the 10x Visium DLPFC dataset. Row: Eleven tissue sections in the 10x Visium DLPFC dataset (except the **representative section 151673).** First column: Ground truth annotations. The remaining columns: Spatial domains identified by SpaSEG and the competing methods. ARI value obtained by each method is shown.

**Supplementary Fig. 3.**
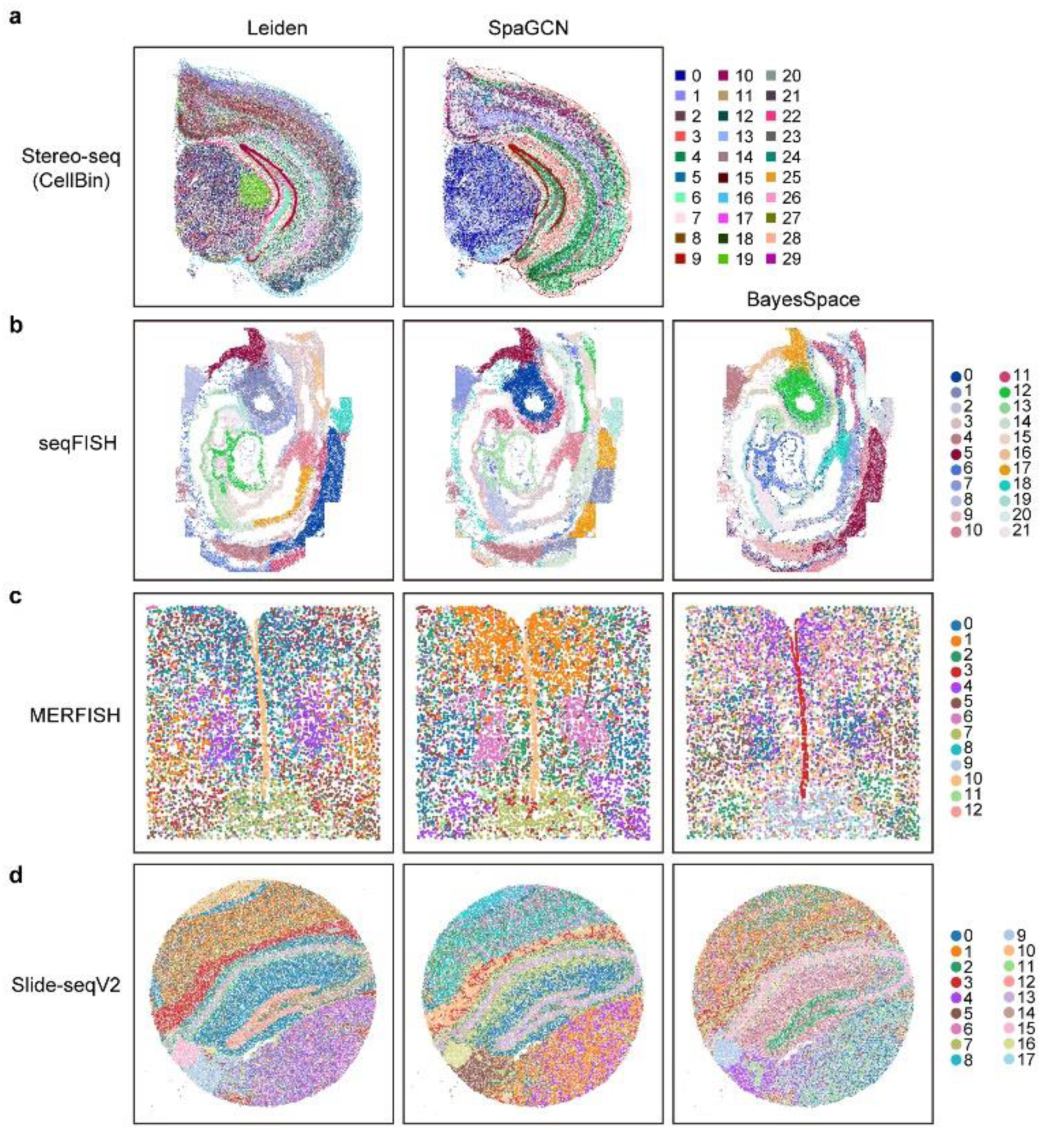
Spatial domain identified by the competing methods for SRT datasets generated by different SRT platforms. **a-d**, Visualization of Spatial domains identified by Leiden, SpaGCN, and BayesSpace for the mouse hemibrain Stereo-seq data at CellBin resolution (**a**), mouse organogenesis seqFISH data (**b**), mouse hypothalamic preoptic region MERFISH data (**c**), and mouse hippocampus Slide-seqV2 data (**d**). BayesSpace is unable to perform spatial clustering on the mouse hemibrain Stereo-seq data at CellBin resolution.

**Supplementary Fig. 4.**
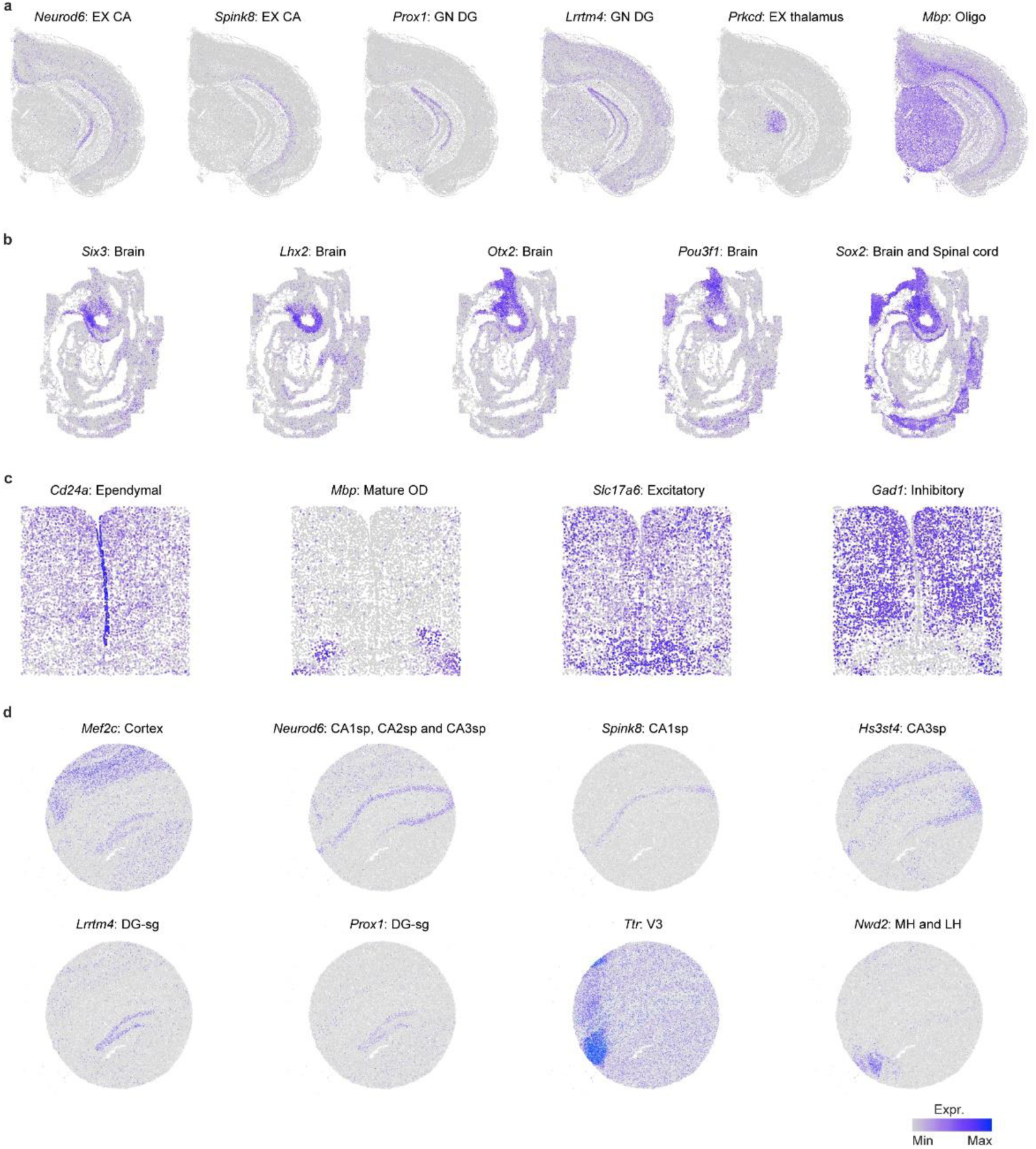
Spatial expression of marker genes for specific spatial domains or cell types. **a**, Spatial expression of marker genes *Neurod6* and *Spink8* for EX CA, *Prox1* and *Lrrtm4* for GN DG, *Prkcd* for EX thalamus, and *Mbp* for Oligo in the mouse hemibrain Stereo-seq data at CellBin resolution. **b**, Spatial expression of marker genes *Six3*, *Lhx2*, *Otx2*, *Pou3f1*, and *Sox2* for spatial domains of brain and spinal cord in mouse organogenesis seqFISH data. **c**, Spatial expression of marker gene *Cd24a* for ependymal cells, *Mbp* for mature OD, *Slc17a6* for excitatory cells, and *Gad1* for inhibitory cells in mouse hypothalamic preoptic region MERFISH data. **d**, Spatial expression of marker genes *Mef2c* for spatial domains of cortex, *Neurod6*, *Spink8* and *Hs3st4* for CA3sp, *Lrrtm4* and *Prox1* for DG-sg, *Ttr* for V3, and *Nwd2* for MH and LH in mouse hippocampus Slide-seqV2 data.

**Supplementary Fig. 5.**
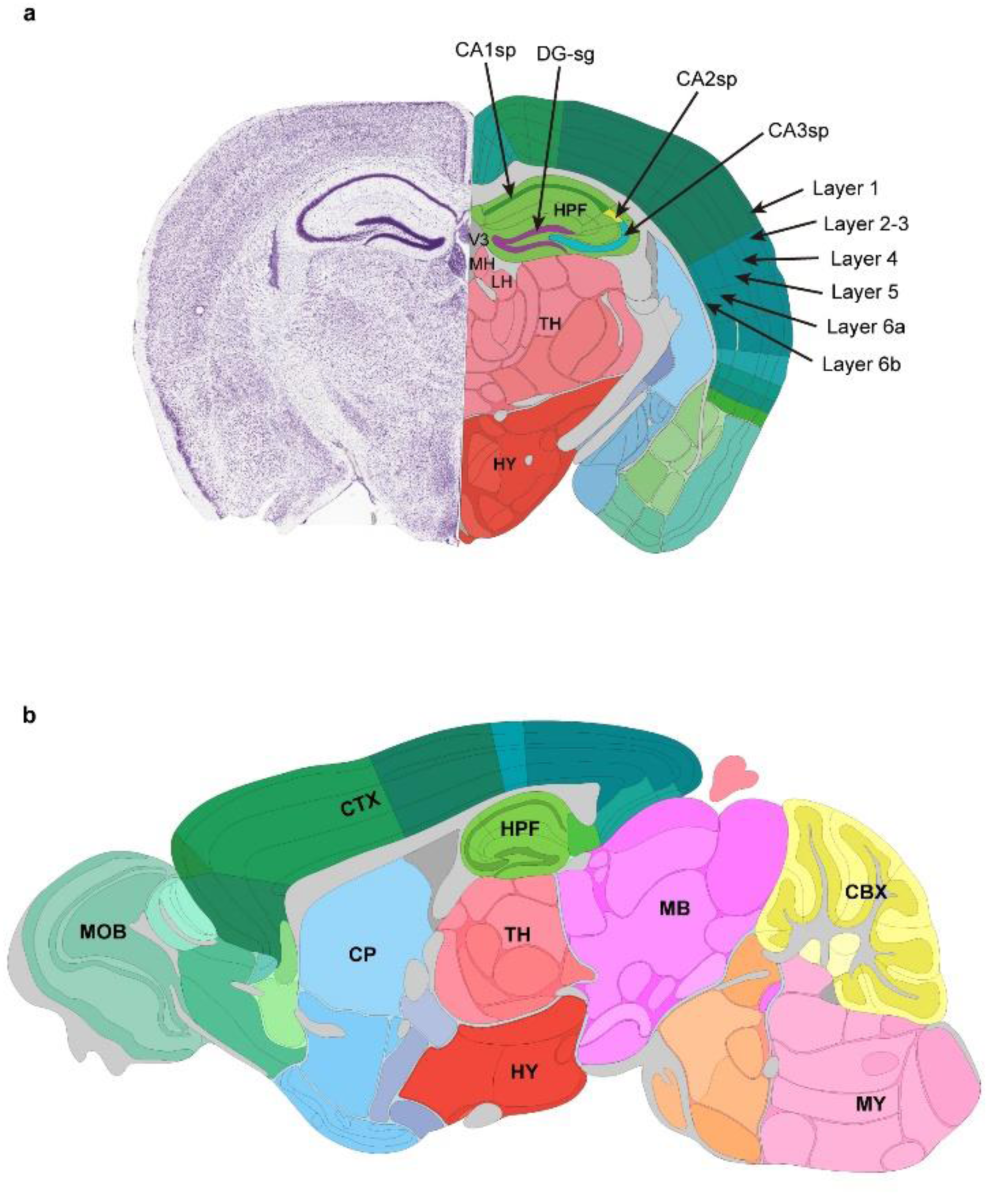
Annotations of mouse brain from the Allen Mouse Brain Atlas. **a**, Annotations of the coronal section. **b**, Annotations for the sagittal section.

**Supplementary Fig. 6.**
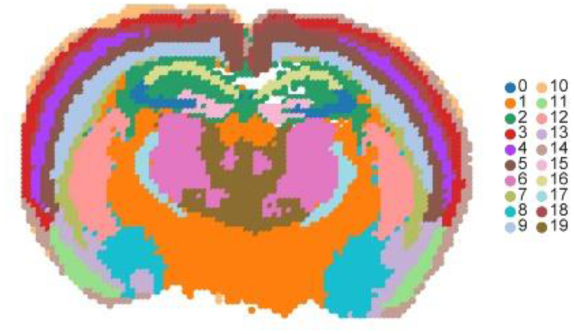
Spatial domains identified by BayesSpace for the whole adult mouse brain Stereo-seq data at Bin200 resolution.

**Supplementary Fig. 7.**
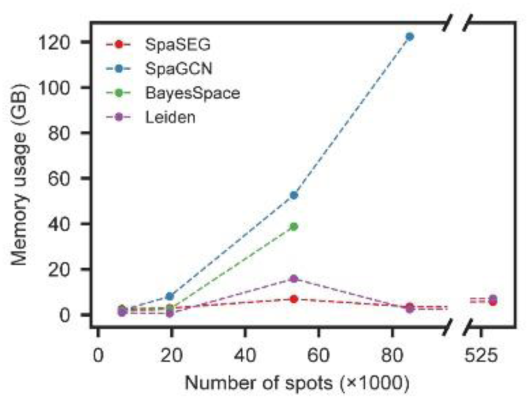
Memory usage (in GB) against varying number of spots in the identification of spatial domains by Leiden, SpaGCN, BayesSpace, and SpaSEG. Given the assayed tissue size, SRT data at higher resolution corresponds to a larger number of spots.

**Supplementary Fig. 8.**
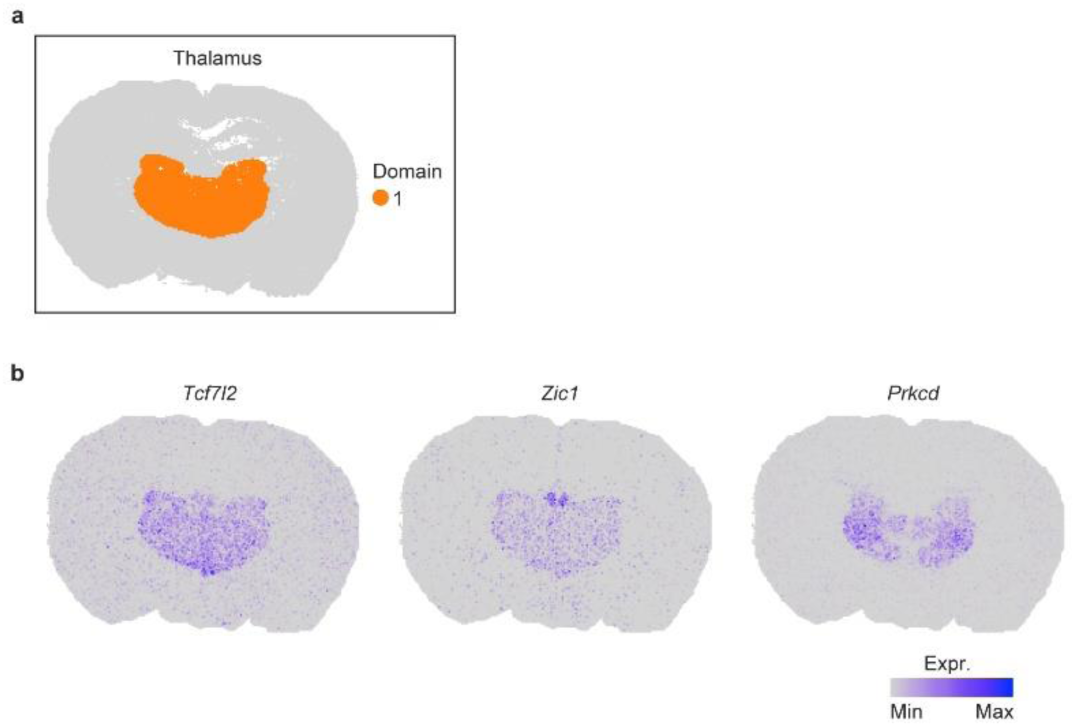
Spatial expression of marker genes for the thalamus in adult mouse brain.

**Supplementary Fig. 9.**
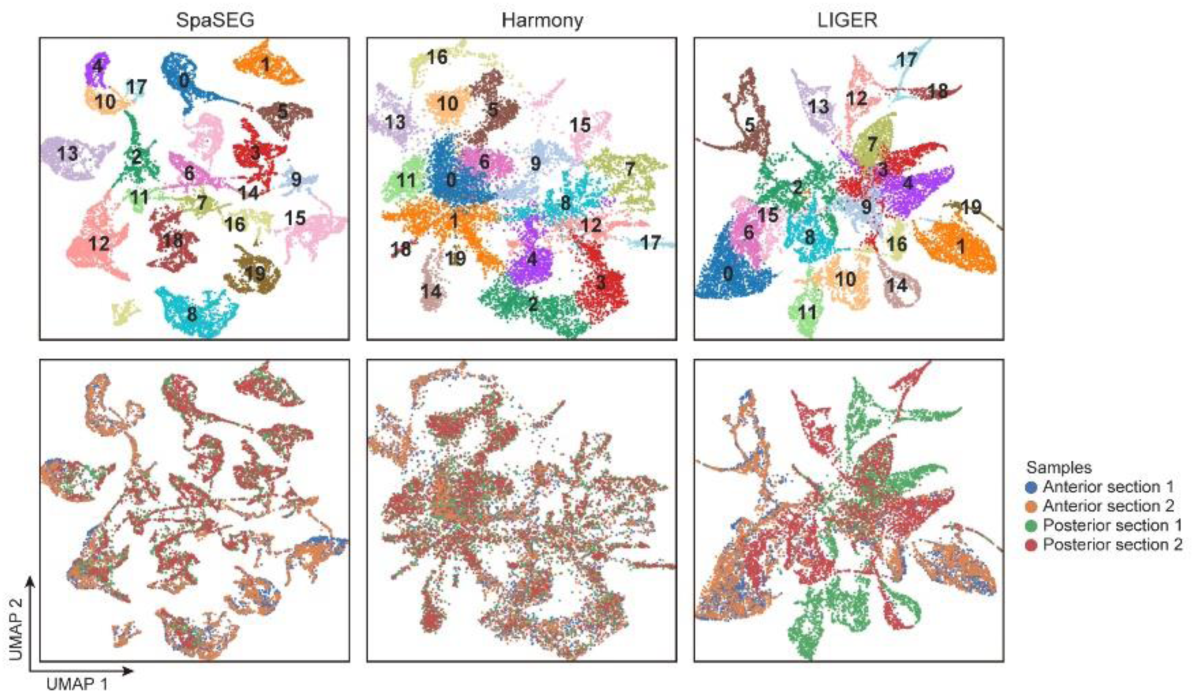
UMAP plots illustrating qualitative evaluation of the SRT data integration over the four sections of mouse brain sections 10x Genomics Visium datasets. Columns illustrate the UMAP results on the basis of the latent representations derived by SpaSEG (Left column), Harmony (Middle column), and LIGER (Right column). Rows represent spatial domains (Top row) and tissue sections (Bottom row).

**Supplementary Fig. 10.**
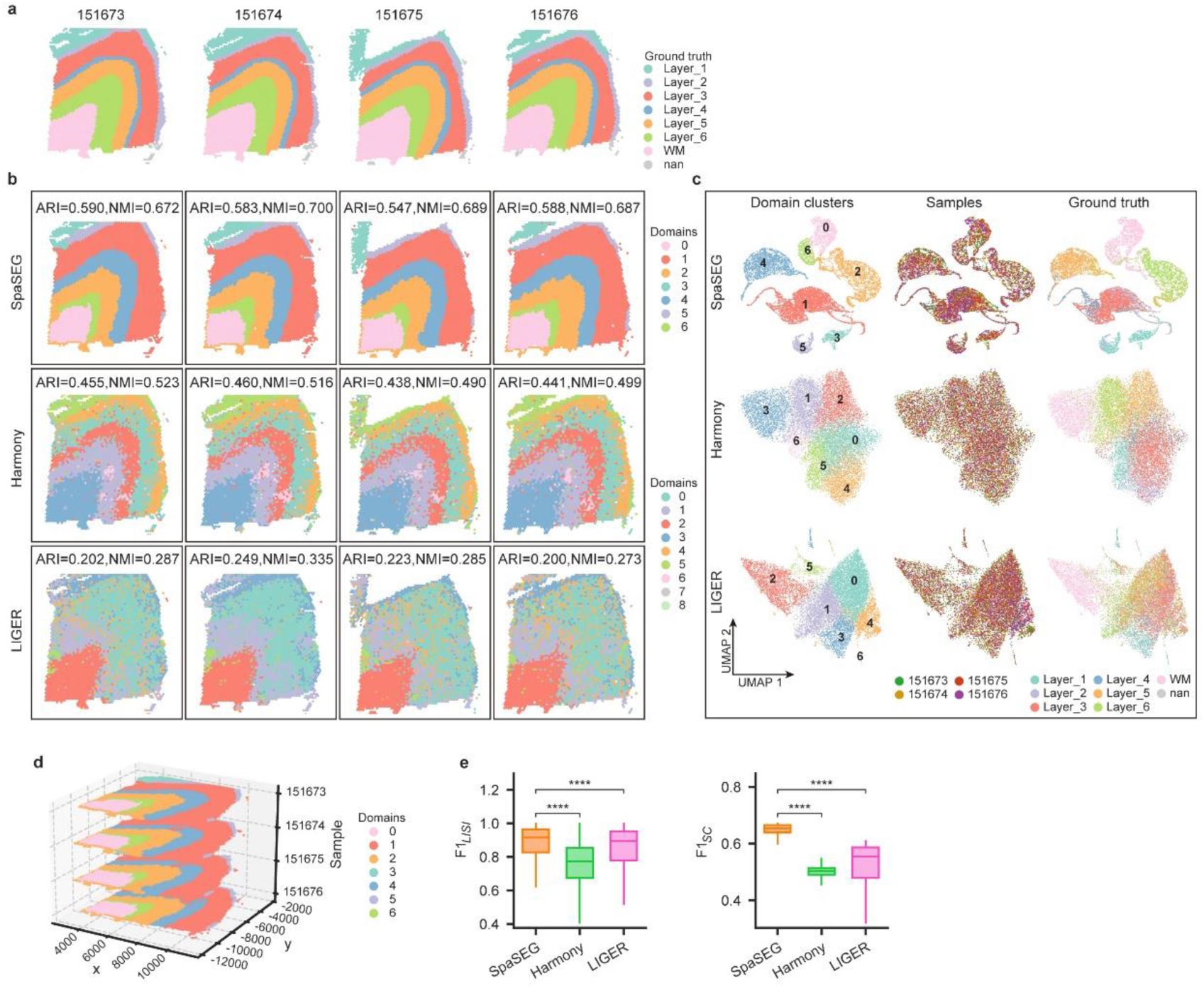
Integrative analysis of adjacent tissue sections of 151673, 151674, 151675, and 151676 from one subject in 10x Visium DLPFC dataset. **a,** Ground truth annotation of the four sections. **b**, Visualization of spatial domains identified and aligned by SpaSEG, Harmony, and LIGER over the four sections. ARI and UMI values were shown for each section. **c**, UMAP plots showing the qualitative evaluation of the SRT data integration by SpaSEG (Top row), Harmony (Middle row), and LIGER (Bottom row). Spatial domains (Left column), tissue sections (Middle columns), and Ground truth annotations (Right column) are colored separately. **d**, Visualization of 3D tissue structure map using the four adjacent sections with SpaSEG-identified spatial domains. **e**, Scores of *F*1_*LISI*_ and *F*1_*SC*_ showing the integration accuracy by SpaSEG, Harmony, and LIGER on the four adjacent sections.

**Supplementary Fig. 11.**
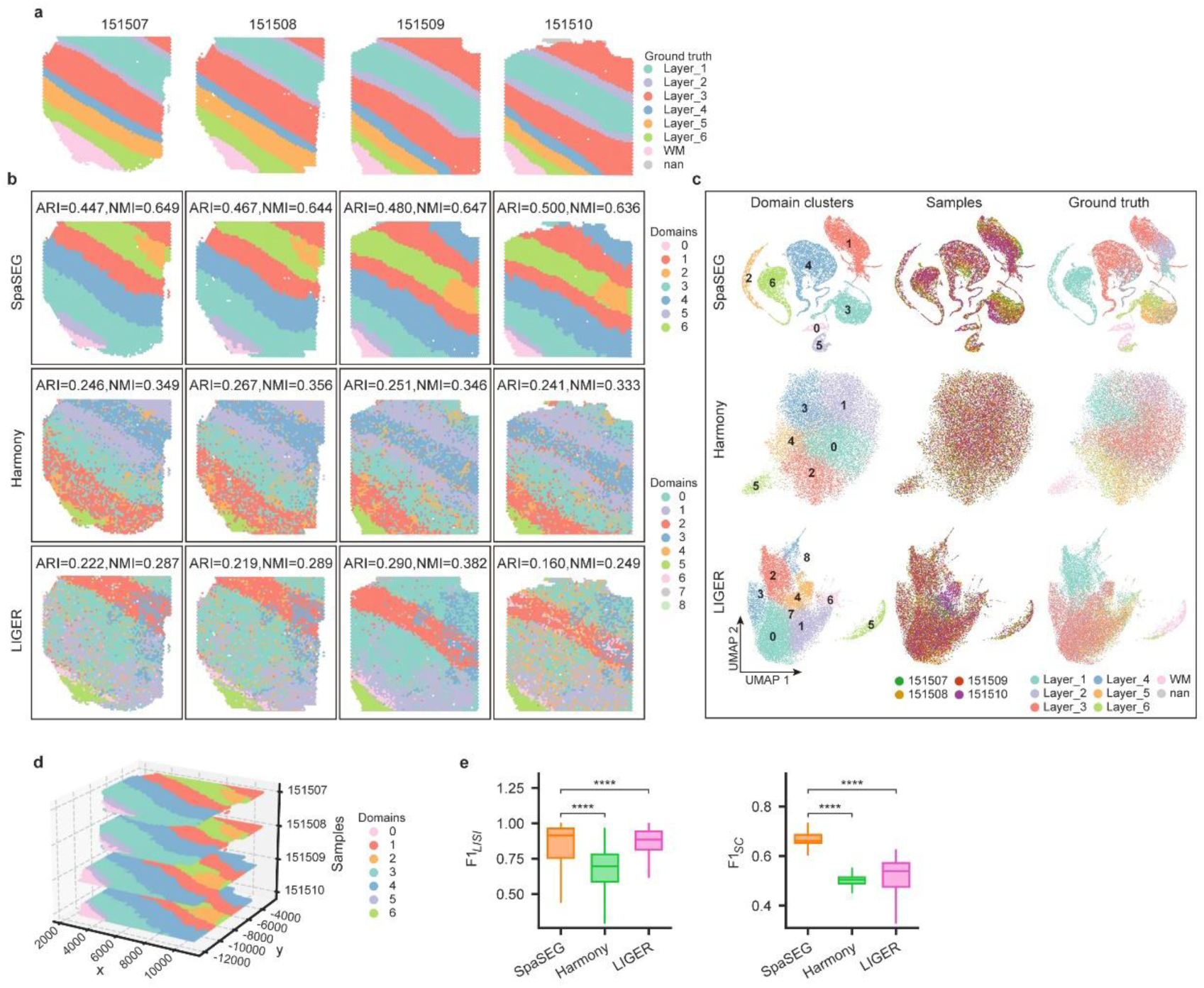
Integrative analysis of adjacent tissue sections of 151507, 151508, 151509, and 151510 from one subject in 10x Visium DLPFC dataset. **a,** Ground truth annotation of the four sections. **b**, Visualization of spatial domains identified and aligned by SpaSEG, Harmony, and LIGER over the four sections. ARI and UMI values were shown for each section. **c**, UMAP plots showing the qualitative evaluation of the SRT data integration by SpaSEG (Top row), Harmony (Middle row), and LIGER (Bottom row). Spatial domains (Left column), tissue sections (Middle columns), and Ground truth annotation (Right column) are colored separately. **d**, Visualization of 3D tissue structure map using the four adjacent sections with SpaSEG-identified spatial domains. **e**, Scores of *F*1_*LISI*_ and *F*1_*SC*_ showing the integration accuracy by SpaSEG, Harmony, and LIGER on the four adjacent sections.

**Supplementary Fig. 12.**
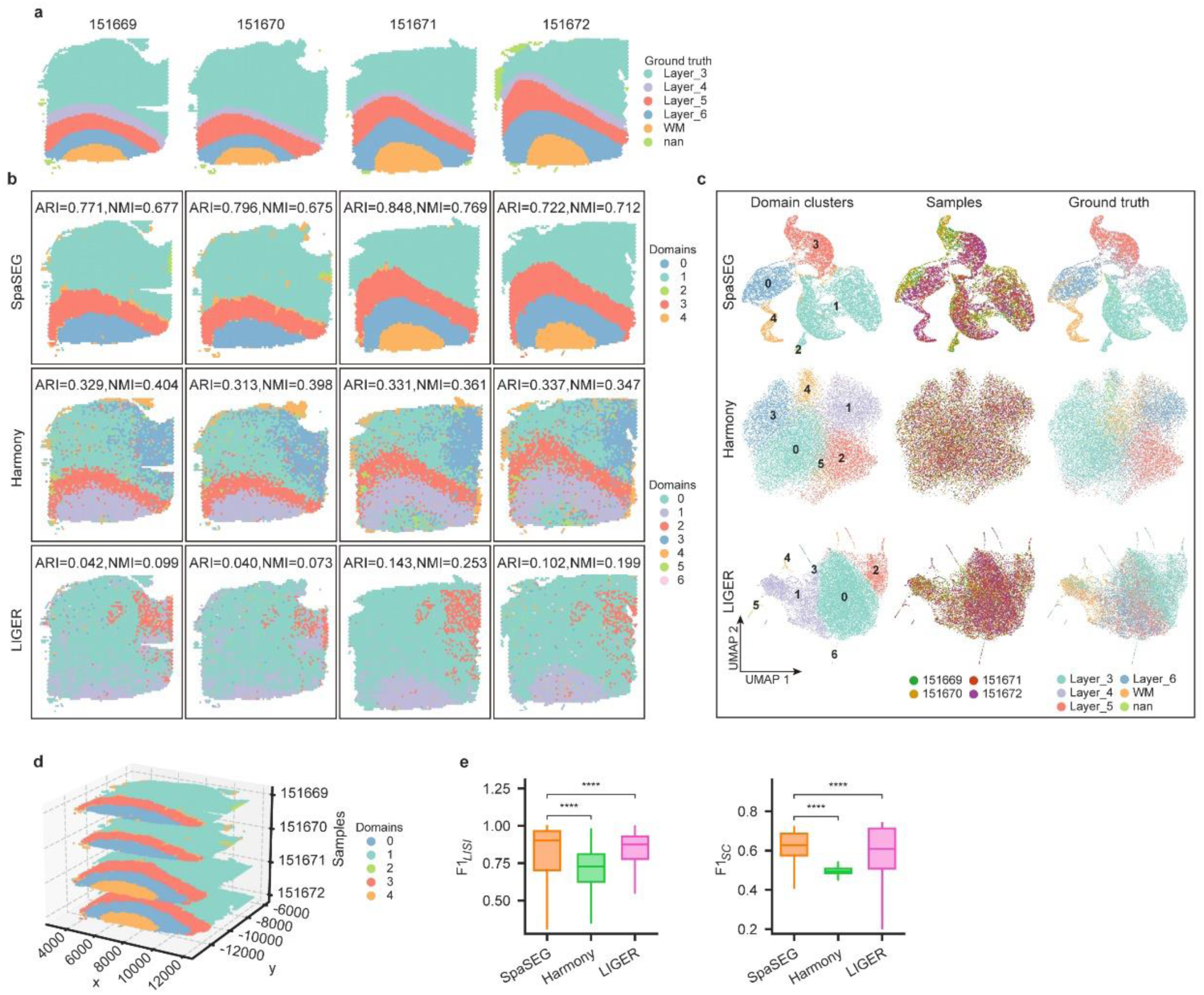
Integrative analysis of adjacent tissue sections of 151669, 151670, 151671, and 151672 from one subject in 10x Visium DLPFC dataset. **a,** Ground truth annotation of the four sections. **b**, Visualization of spatial domains identified and aligned by SpaSEG, Harmony, and LIGER over the four sections. ARI and UMI values were shown for each section. **c**, UMAP plots showing the qualitative evaluation of the SRT data integration by SpaSEG (Top row), Harmony (Middle row), and LIGER (Bottom row). Spatial domains (Left column), tissue sections (Middle columns), and Ground truth annotation (Right column) are colored separately. **d**, Visualization of 3D tissue structure map using the four adjacent sections with SpaSEG-identified spatial domains. **e**, Scores of *F*1_*LISI*_ and *F*1_*SC*_ showing the integration accuracy by SpaSEG, Harmony, and LIGER on the four adjacent sections.

**Supplementary Fig. 13.**
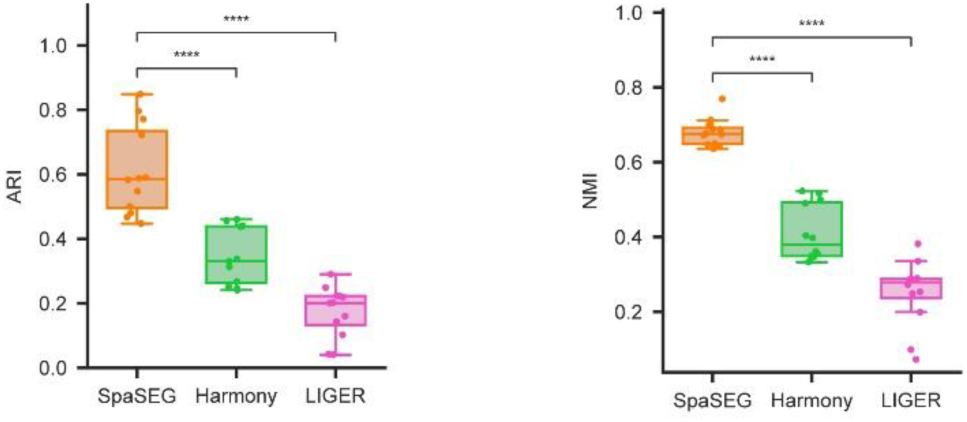
ARI and NMI evaluation metrics showing the integration accuracy by SpaSEG, Harmony, and LIGER for all 12 sections of the 10x Visium DLPFC dataset against competing methods. **** p<0.0001.

**Supplementary Fig. 14.**
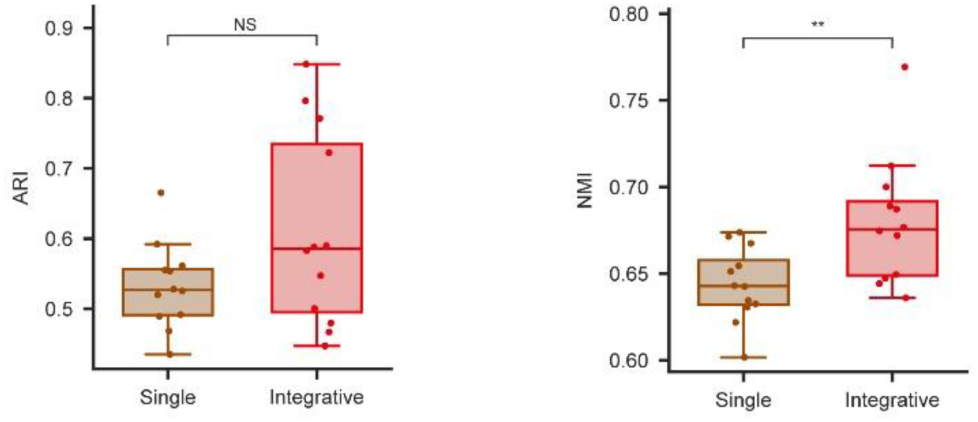
Comparison of ARI and NMI evaluation metrics for spatial domains identified by SpaSEG using each single section against integration of multiple adjacent sections for the 12 sections in the 10x Visium DLPFC dataset. NS: not significant, ** p<0.01.

**Supplementary Fig. 15.**
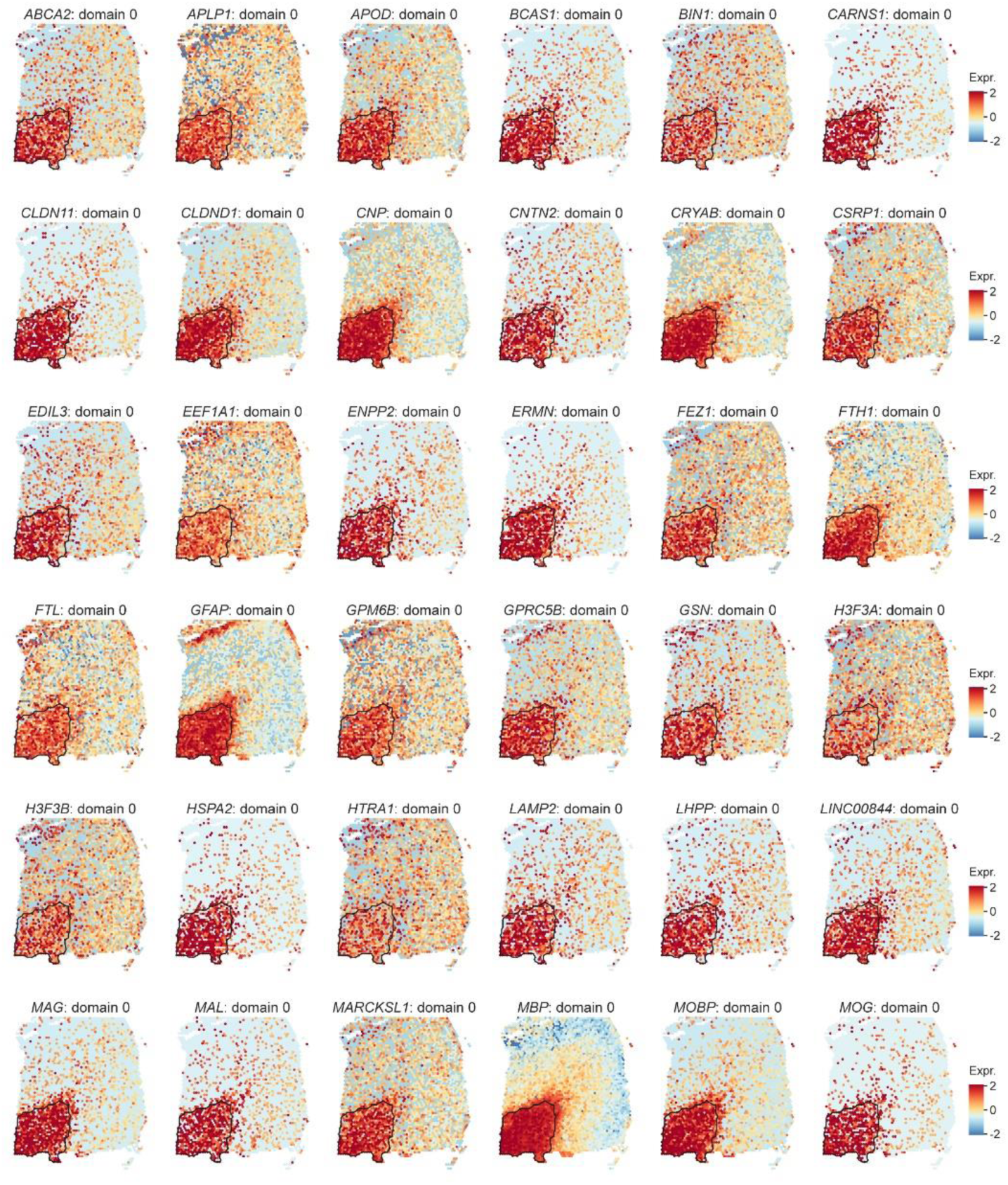
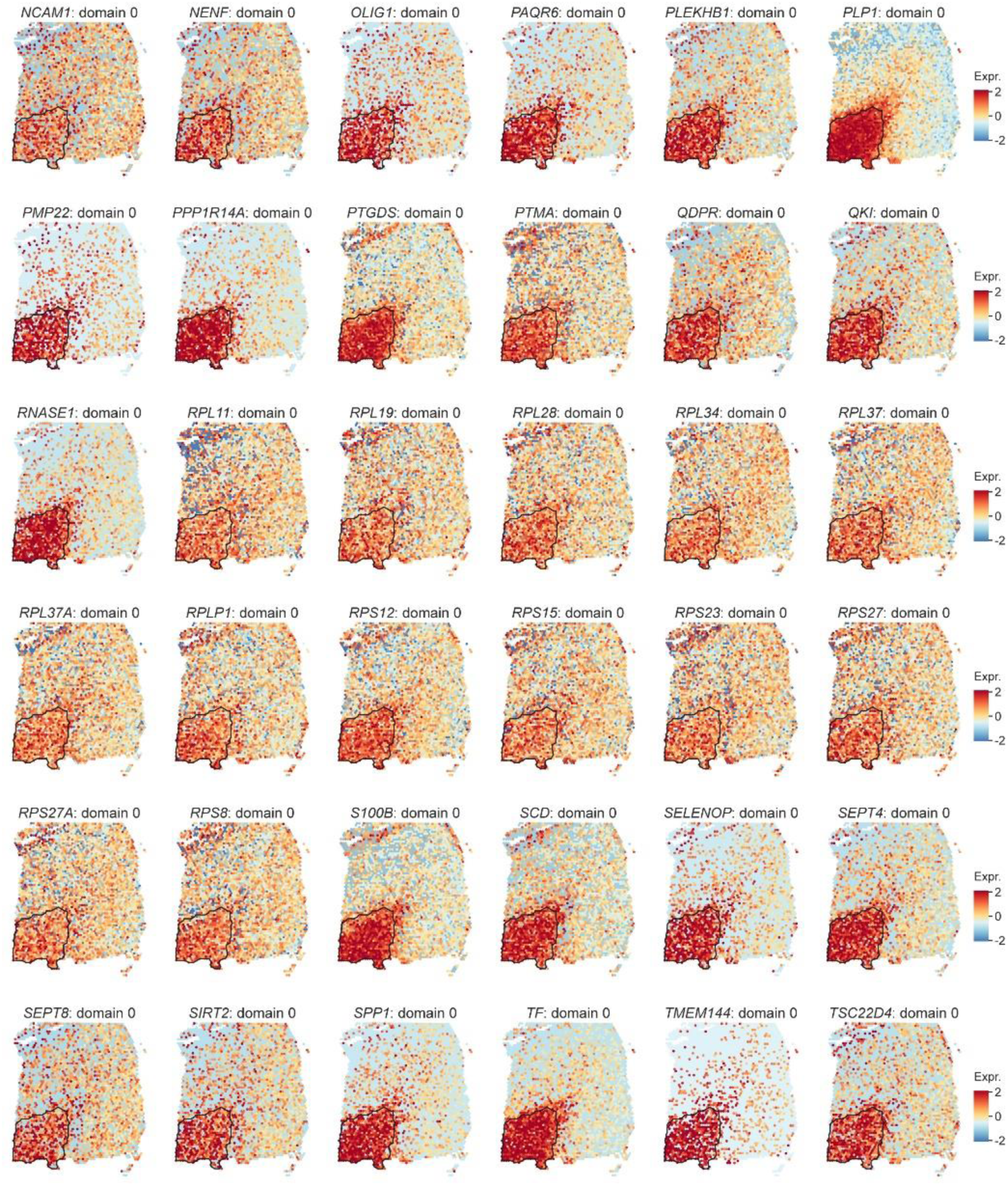
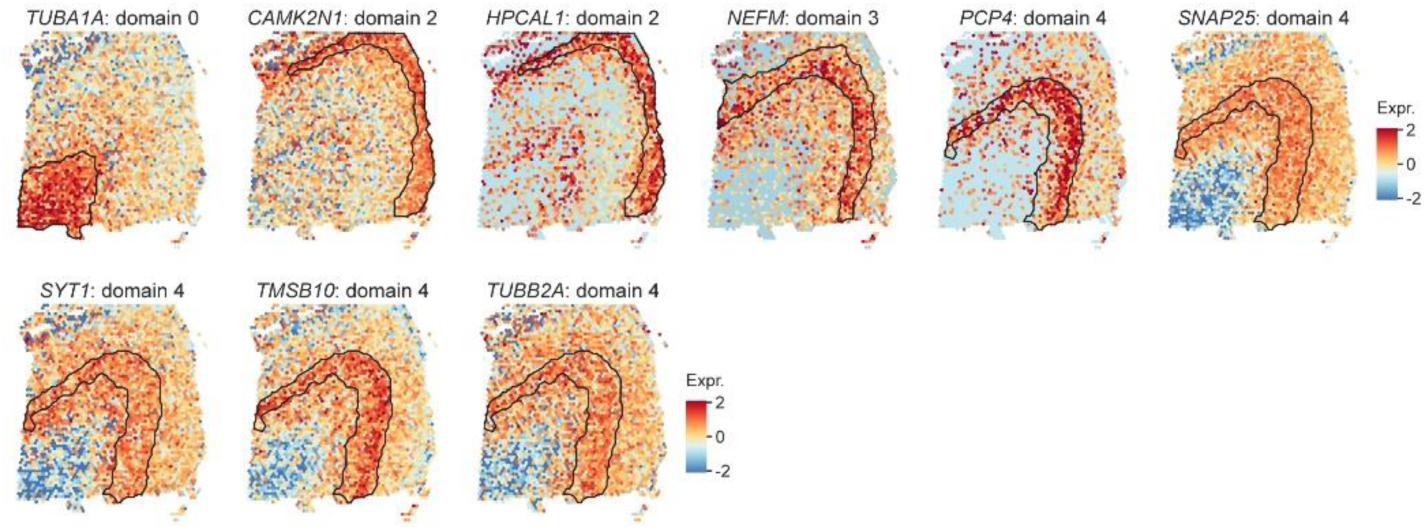
Visualization of all SpaSEG-detected SVGs (n=81) for the section 151673 in the DLPFC 10x Visium data.

**Supplementary Fig. 16.**
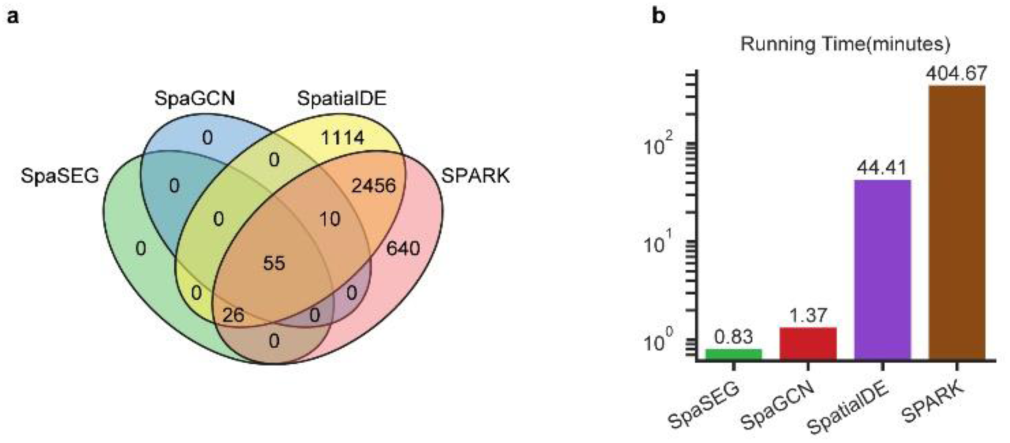
The SVGs that detected by SpaSEG, SpaGCN, SpatialDE, and SPARK for section 151673 in the DLPFC 10x Visium data, and their time consumption. **a**, Venn diagram of the detected SVGs. **b**, Running time taken by each method for SVG detection.

**Supplementary Fig. 17.**
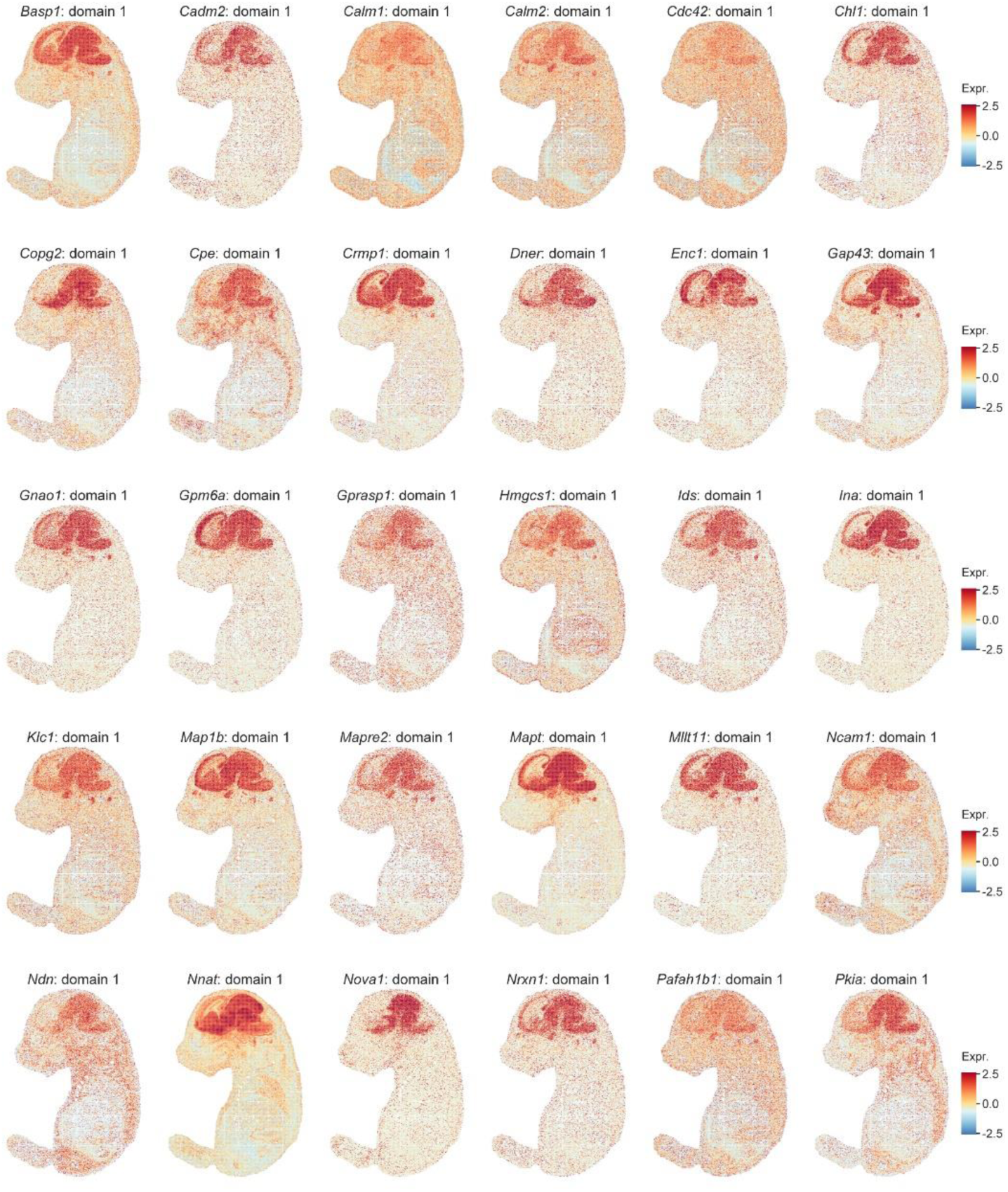
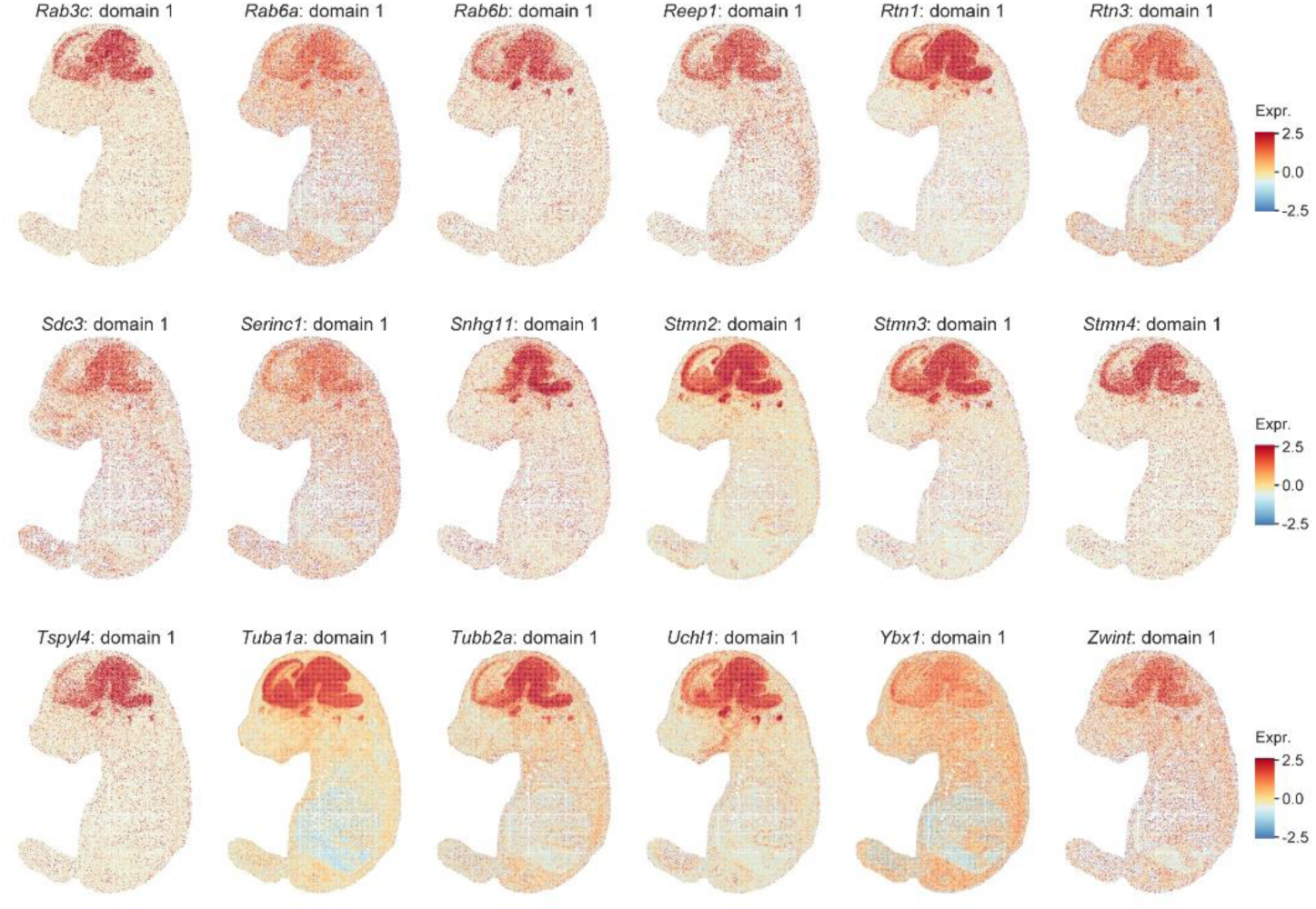
Visualization of SpaSEG-detected SVGs (n=48) for domain 1 (brain) in the E16.5 mouse embryo Stereo-seq dataset at Bin50 resolution.

**Supplementary Fig. 18.**
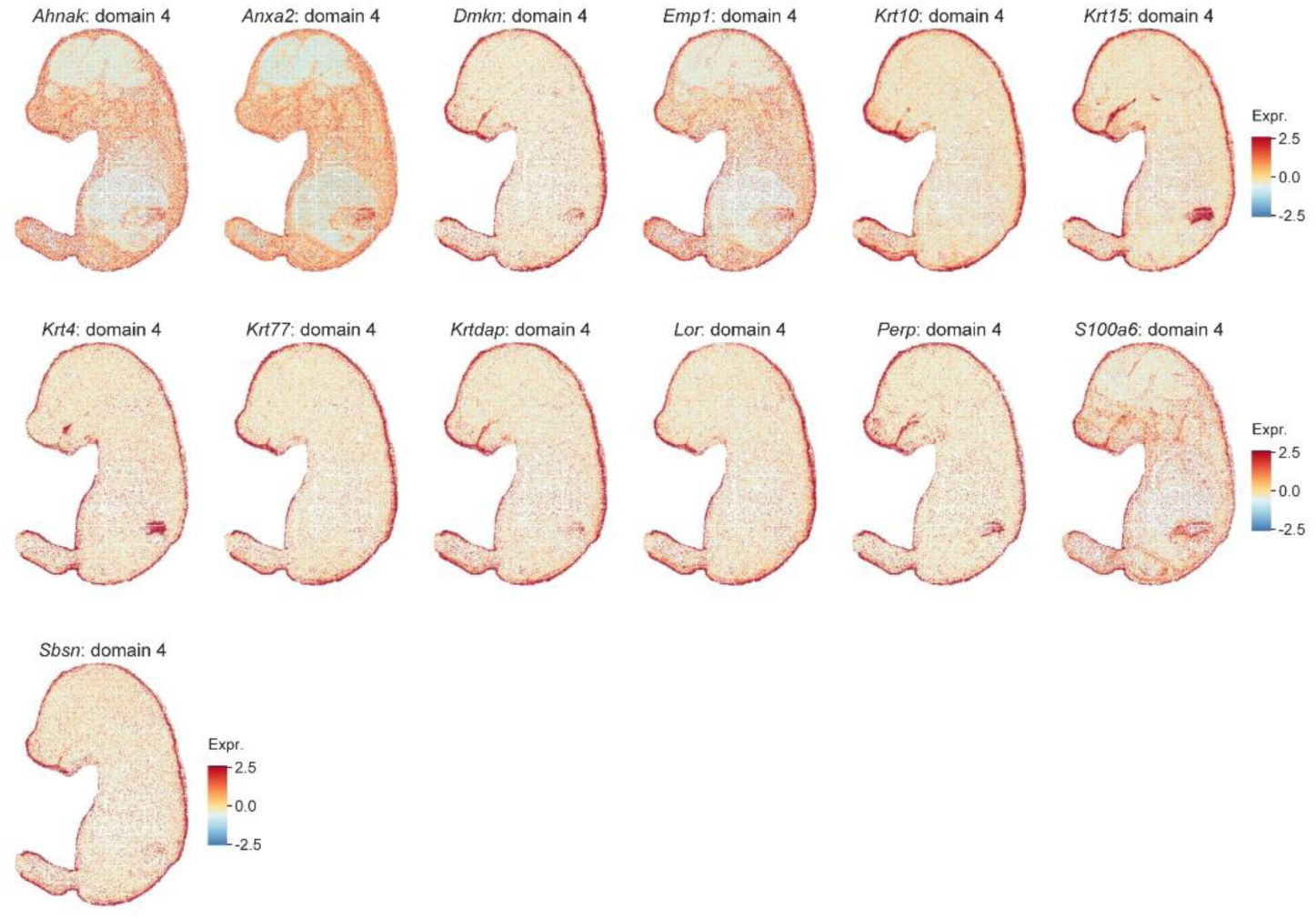
Visualization of SpaSEG-detected SVGs (n=13) for domain 4 (epidermis) in the E16.5 mouse embryo Stereo-seq dataset at Bin50 resolution.

**Supplementary Fig. 19.**
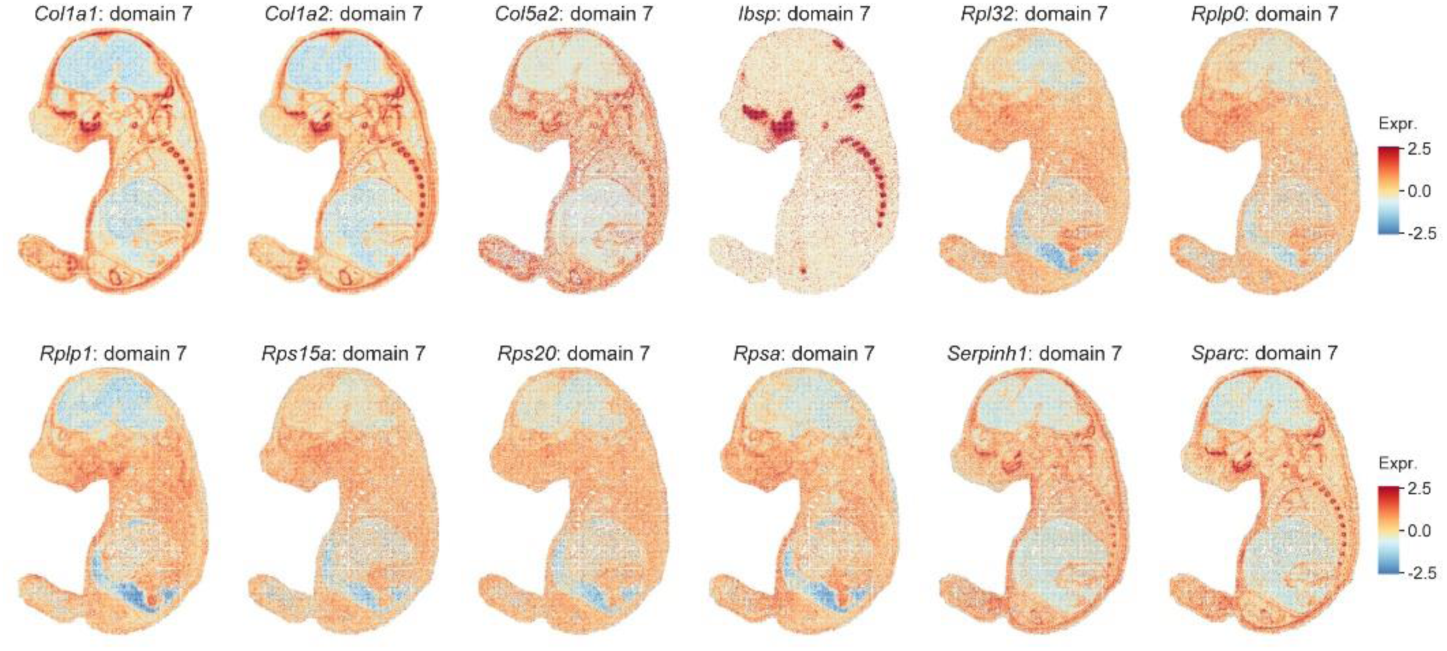
Visualization of SpaSEG-detected SVGs (n=12) for domain 7 (cartilage primordium/bone) in the E16.5 mouse embryo Stereo-seq dataset at Bin50 resolution.

**Supplementary Fig. 20.**
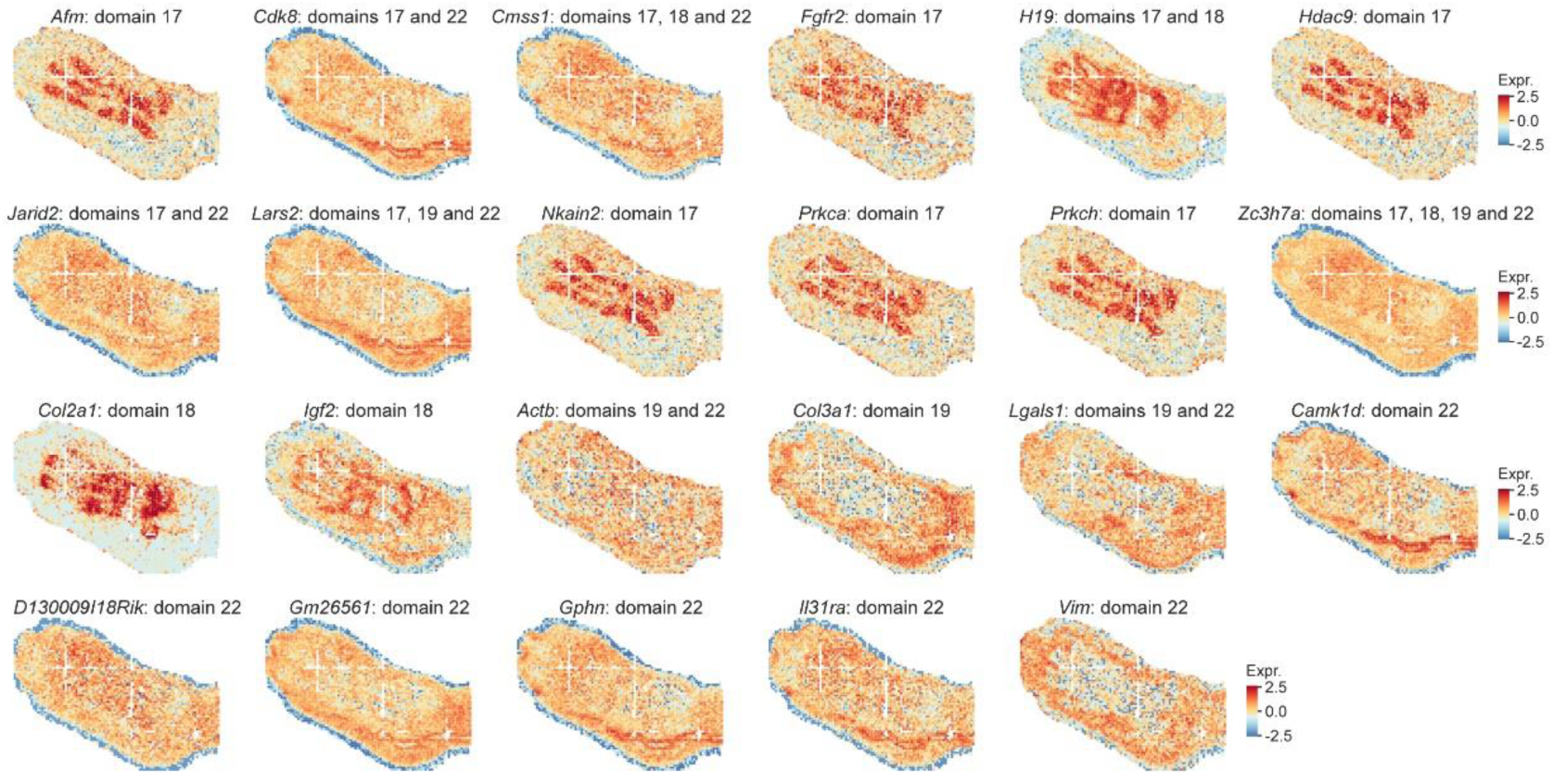
Visualization of SpaSEG-detected SVGs for limb (n=23) in the E16.5 mouse embryo Stereo-seq dataset at Bin50 resolution. We excluded SVGs linked to domain 4.

**Supplementary Fig. 21.**
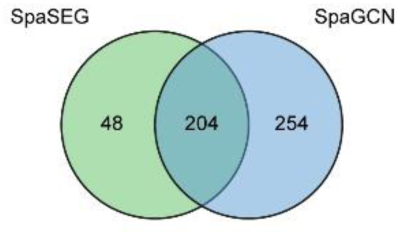
Venn diagram for the SVGs detected by SpaSEG and SpaGCN for the E16.5 mouse embryo Stereo-seq dataset at Bin50 resolution.

**Supplementary Fig. 22.**
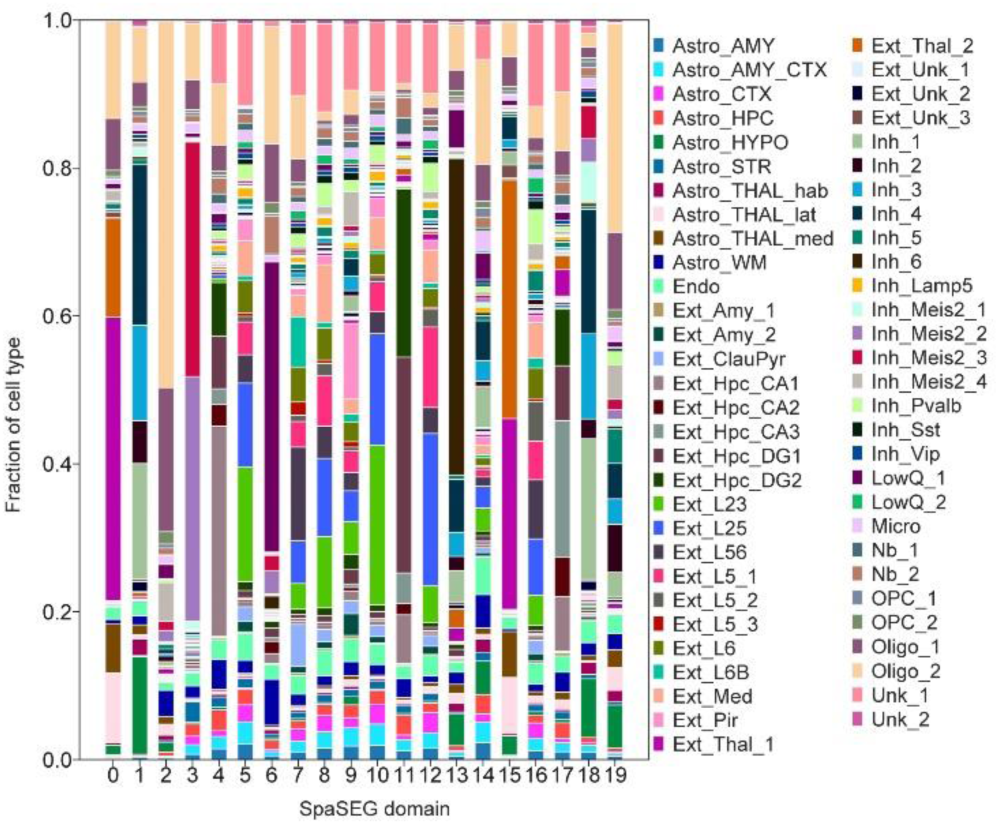
Fraction of Cell type in each SpaSEG-identified spatial domain in the adult mouse brain Stereo-seq data at Bin200 resolution. Cell types were deconvolved by the cell2location algorithm. Fraction is calculated using the estimated cell type abundances per domain.

**Supplementary Fig. 23.**
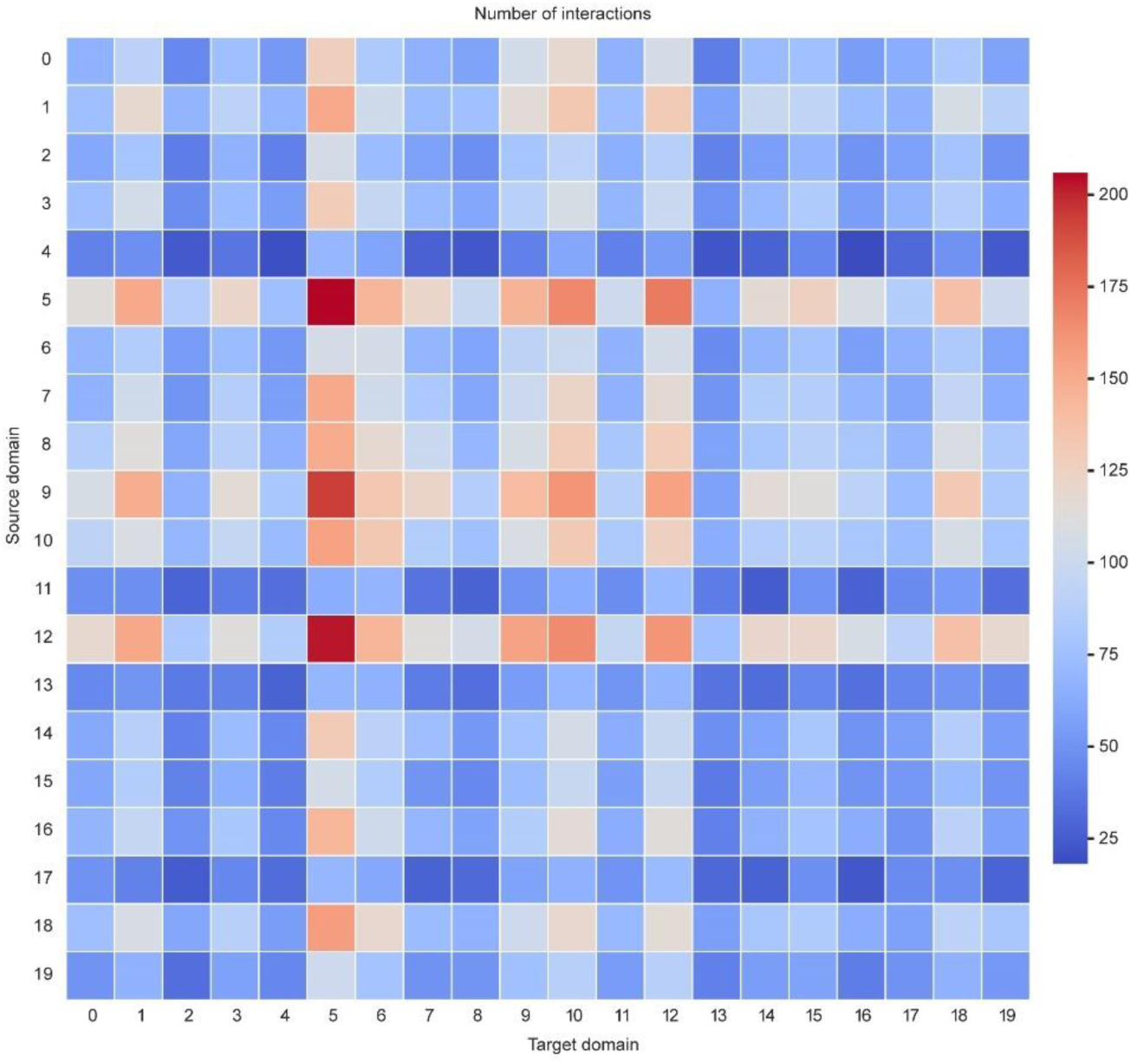
Interaction heatmap plotting the total number of interactions between all source domains (y axis) and target domains (x axis) in the whole adult mouse brain at Bin200 resolution.

**Supplementary Fig. 24.**
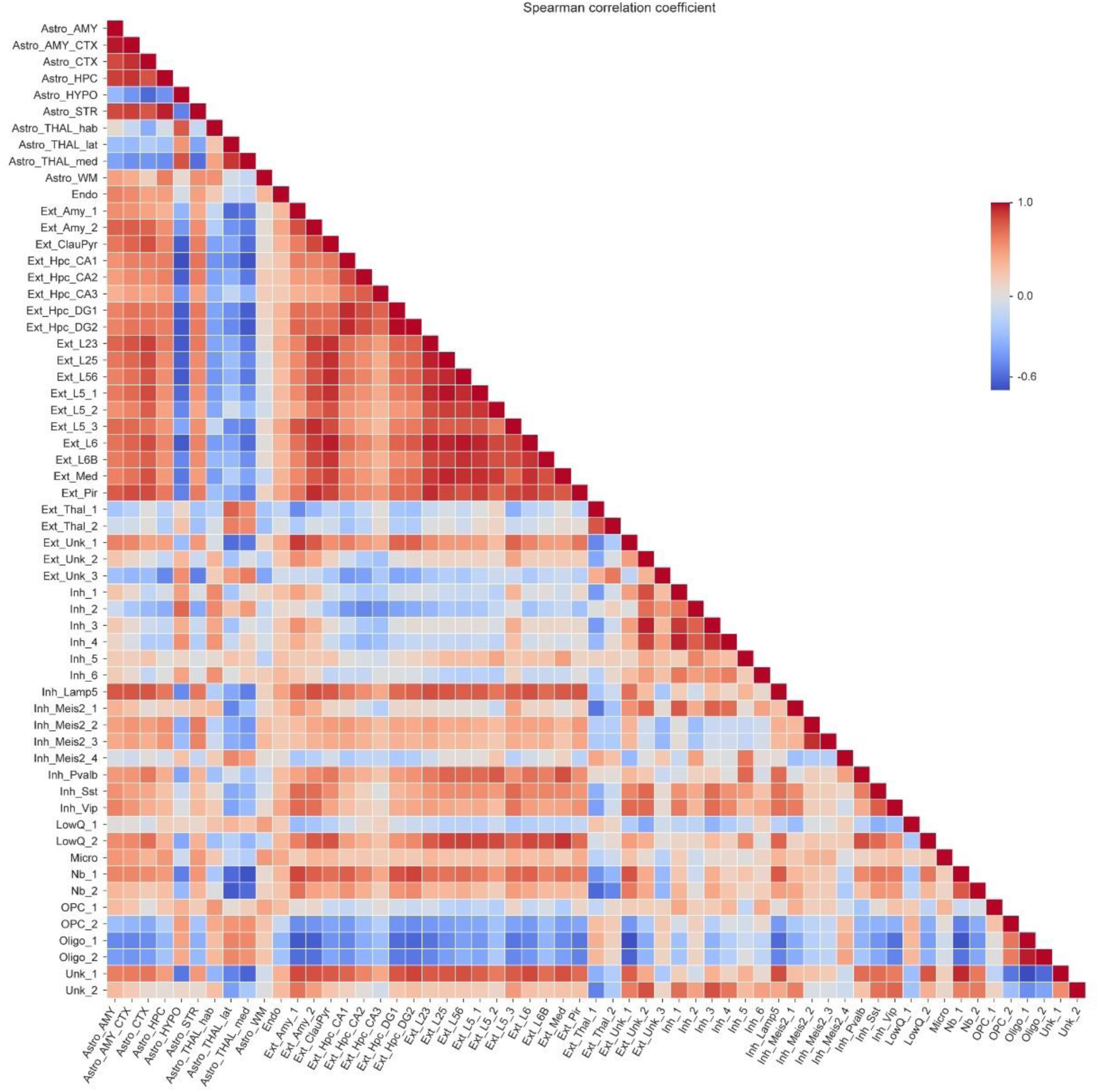
Spearman correlation coefficient (SCC) between cell2location-derived cell types in the whole adult mouse brain at Bin200 resolution.

**Supplementary Fig. 25.**
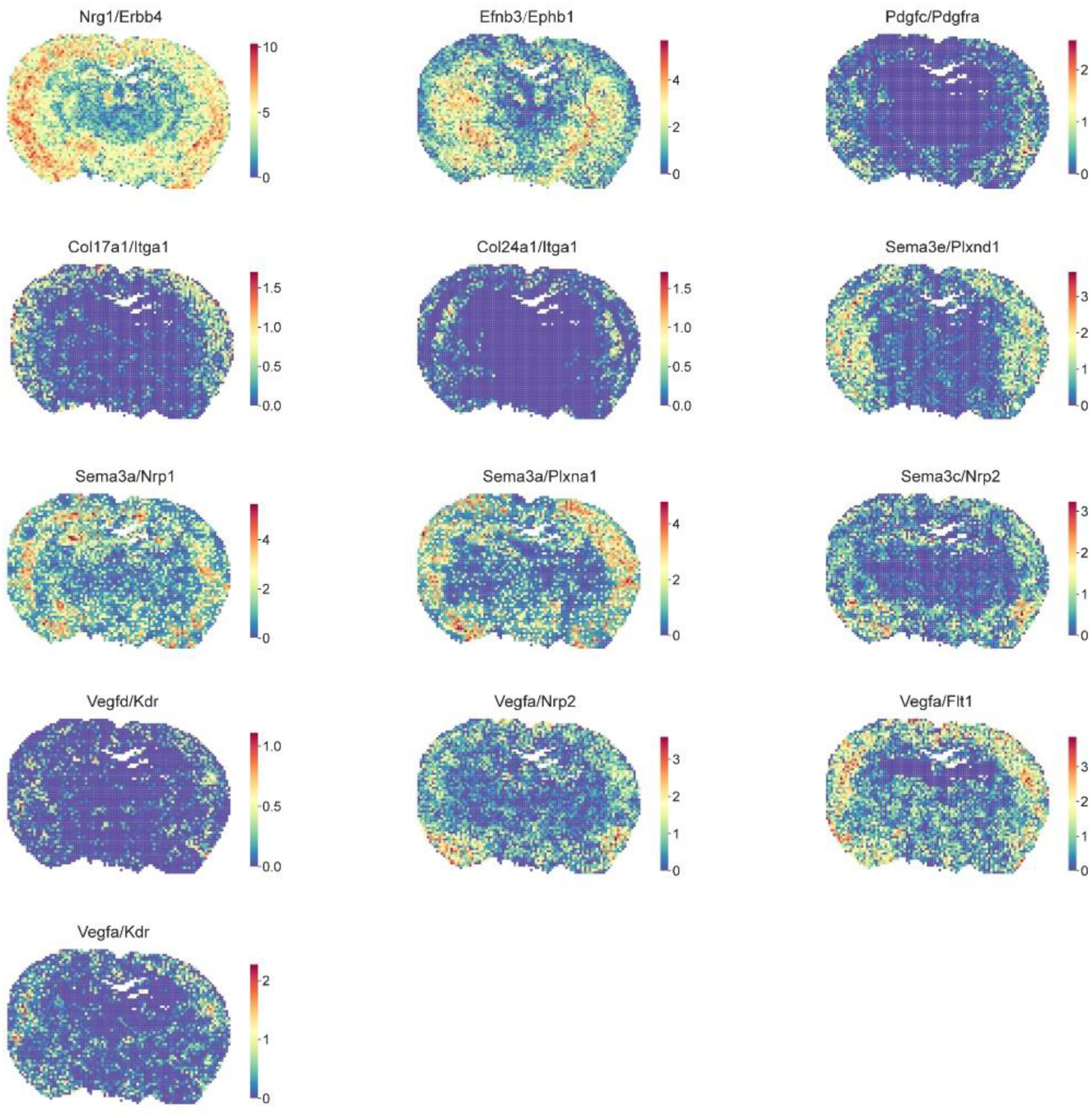
Visualization of additional spot-wise scores of ligand-receptor pairs corresponding to those shown in Fig. 5i.

**Supplementary Fig. 26.**
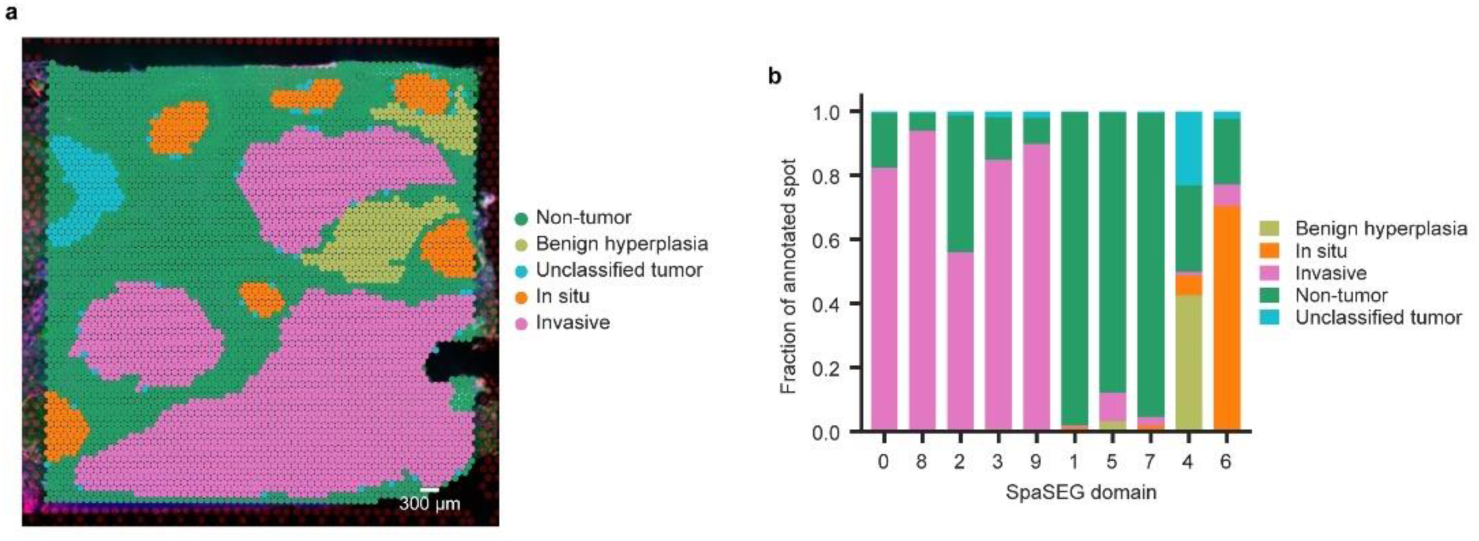
Annotation for the 10x Visium IDC dataset. **a,** Visualization of spot-wise annotations. **b,** Fraction of annotated spots in each SpaSEG-identified spatial domain.

**Supplementary Fig. 27.**
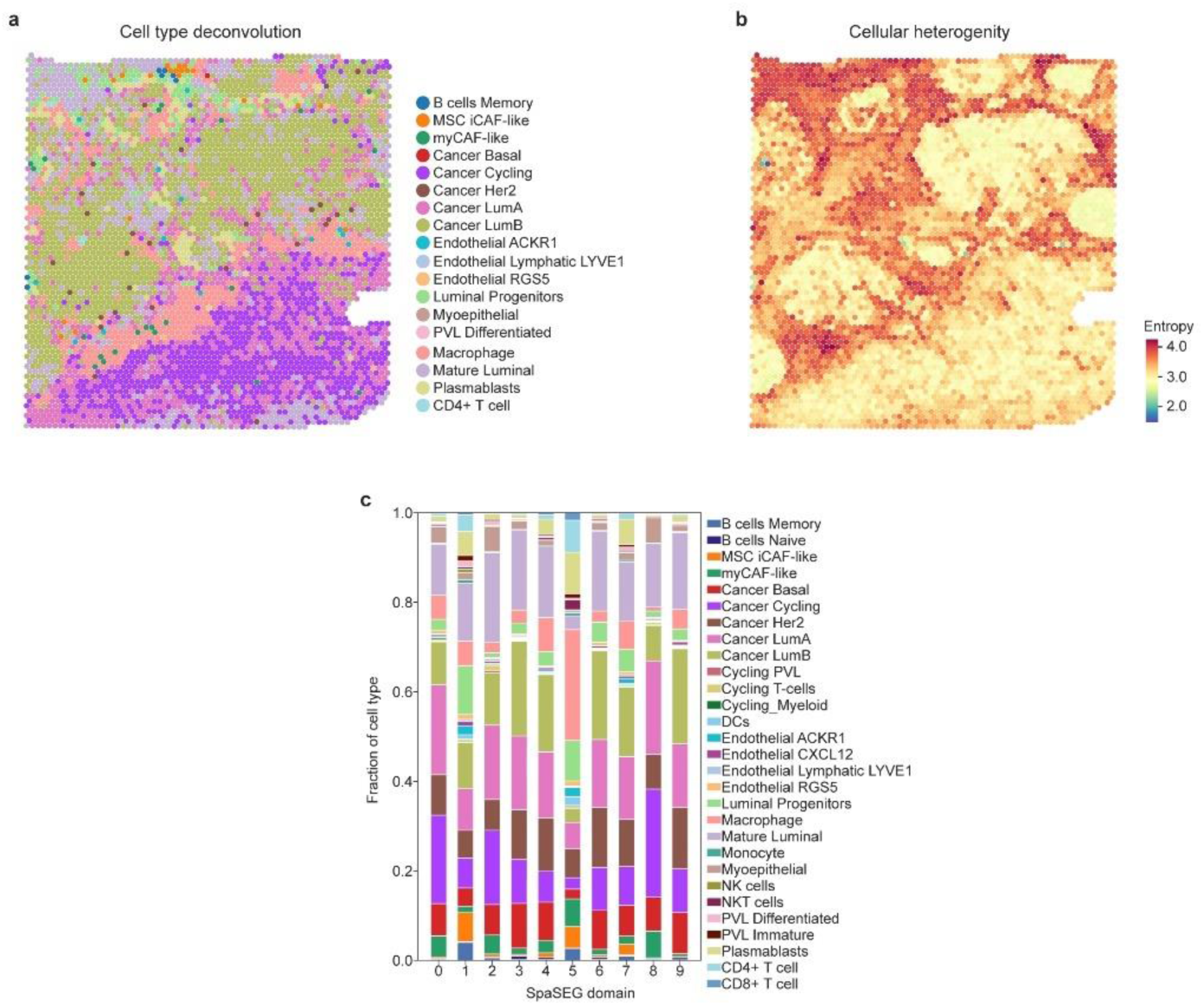
Result of cell type deconvolution using the cell2location algorithm for the 10x Visium IDC dataset. **a**, Visualization of spatial distribution of cell types. Each spot is represented by the cell type with the highest abundance. **b**, Entropy score showing cell type heterogeneity per spot. **c**, Fraction of cell types in each SpaSEG-identified spatial domain in 10x Visium IDC dataset. Fraction is calculated using the estimated cell type abundance per domain.

**Supplementary Fig. 28.**
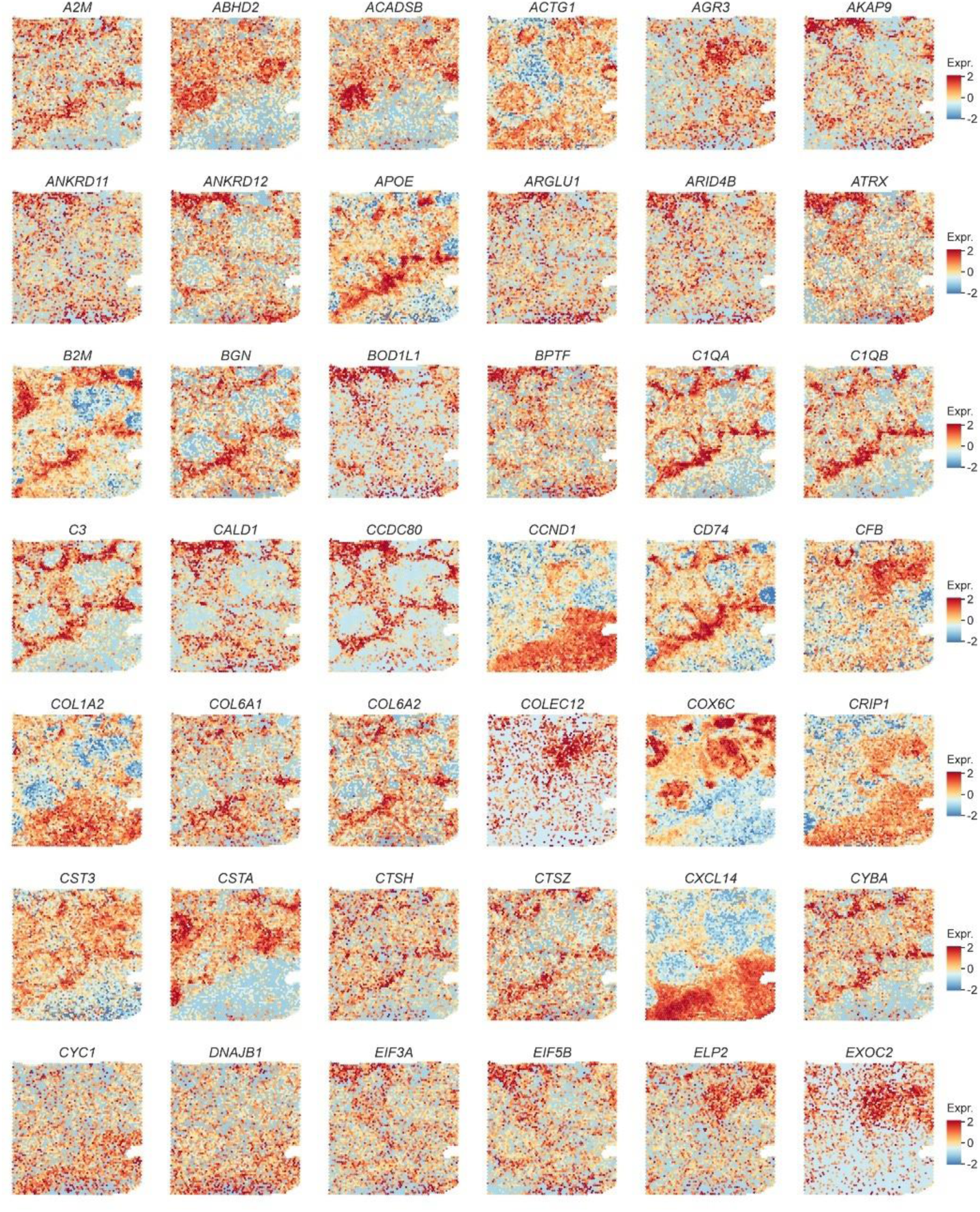
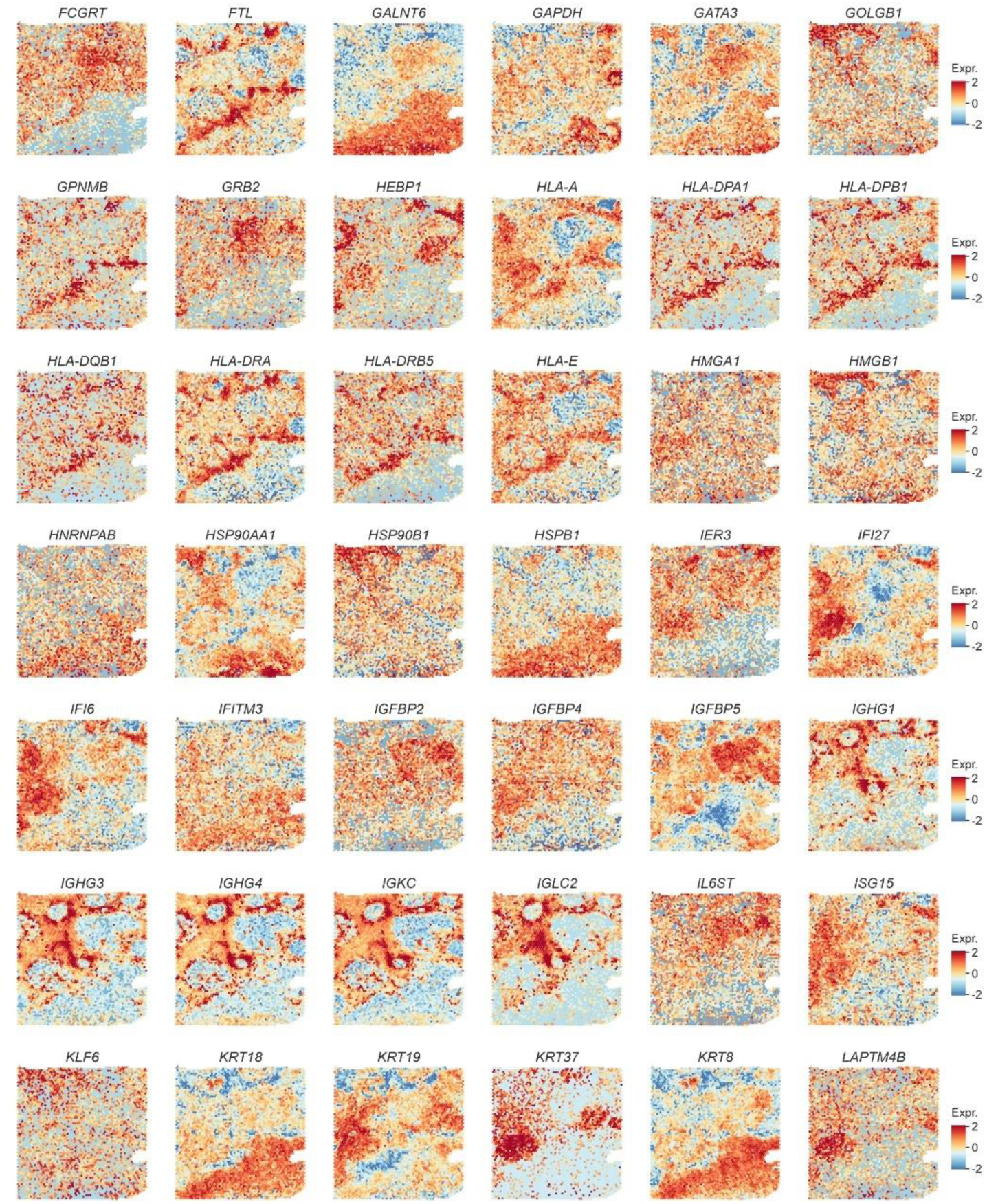
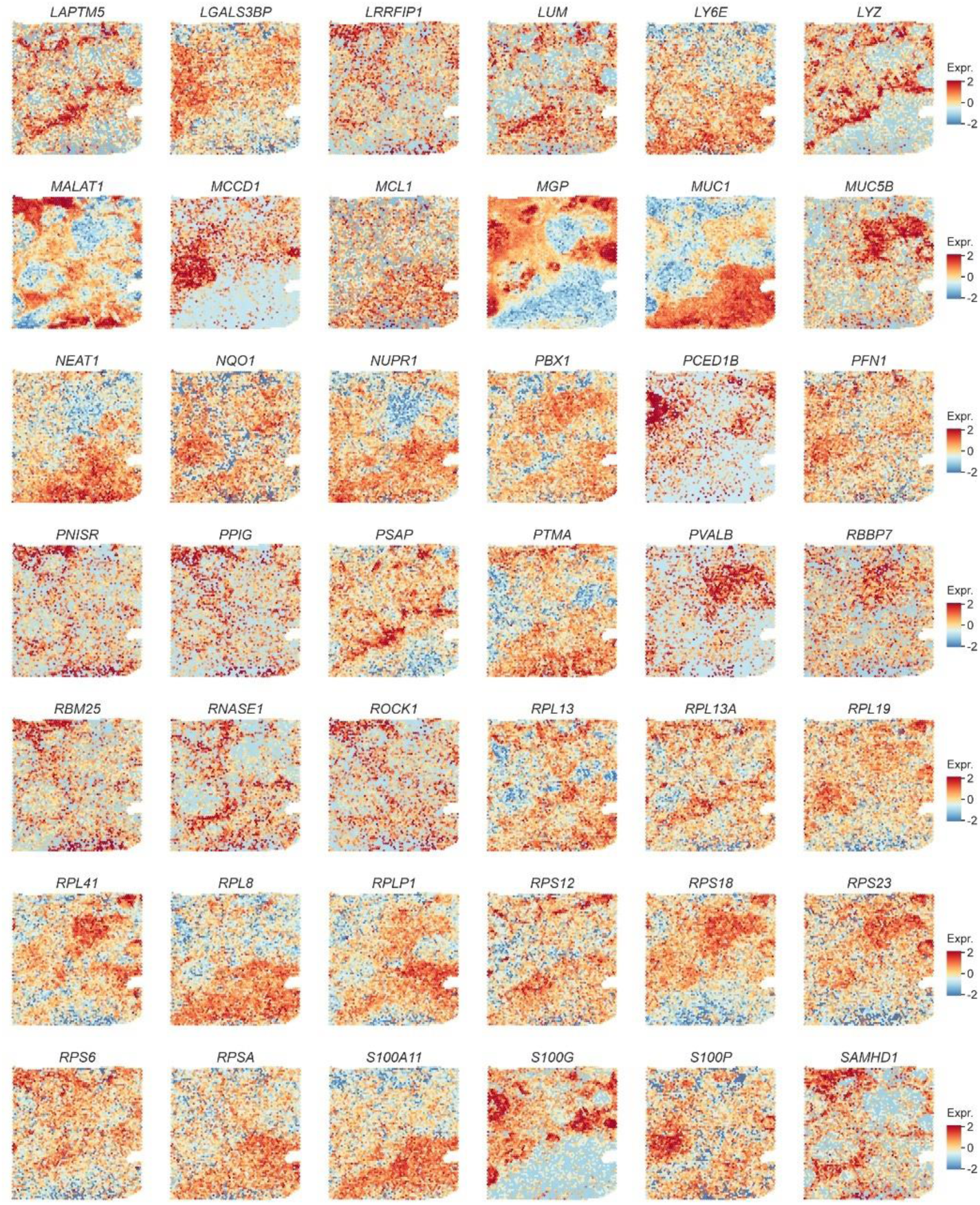
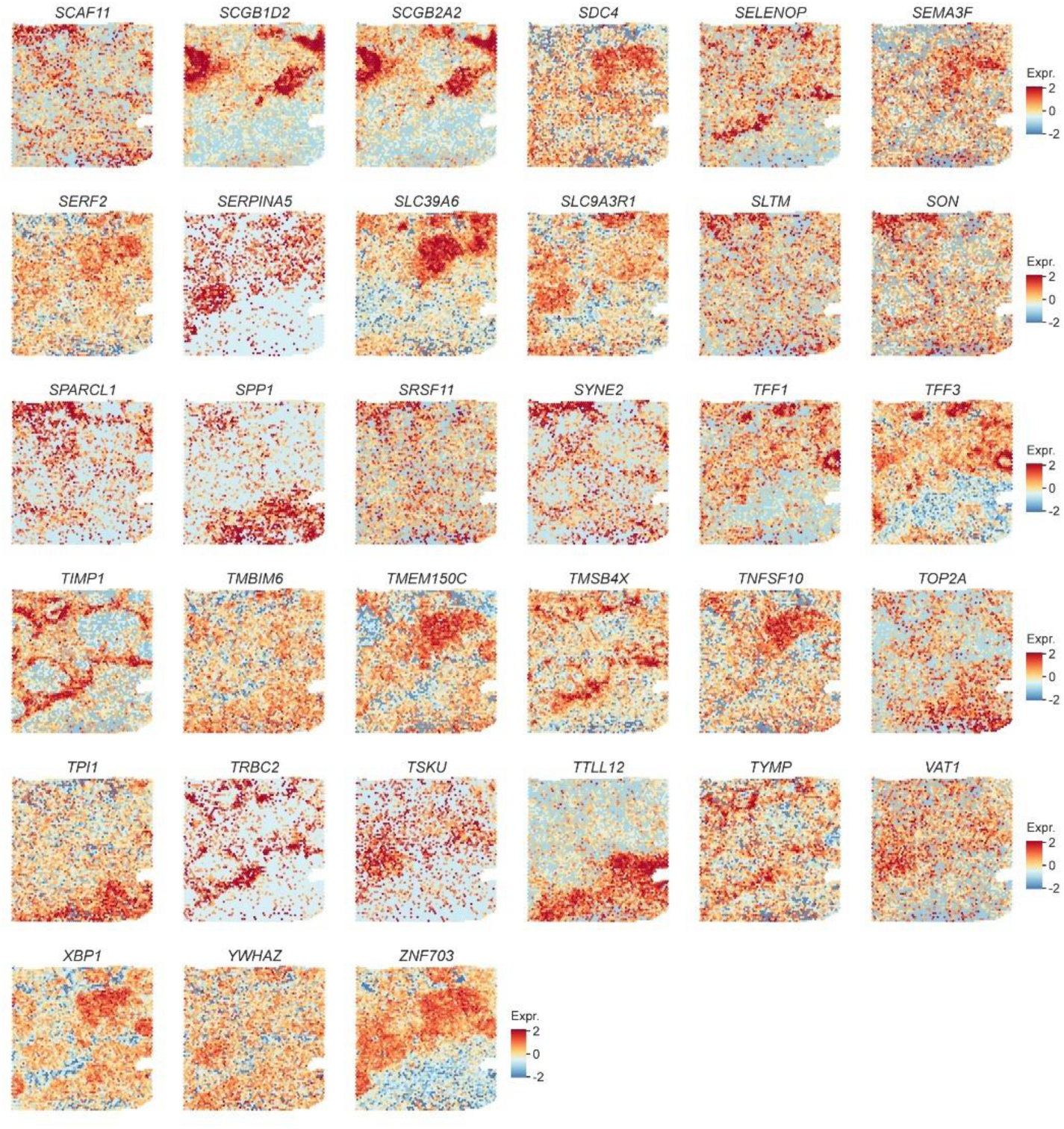
Visualization of all SpaSEG-detected SVGs (n=159) for the 10x Visium IDC dataset.

**Supplementary Fig. 29.**
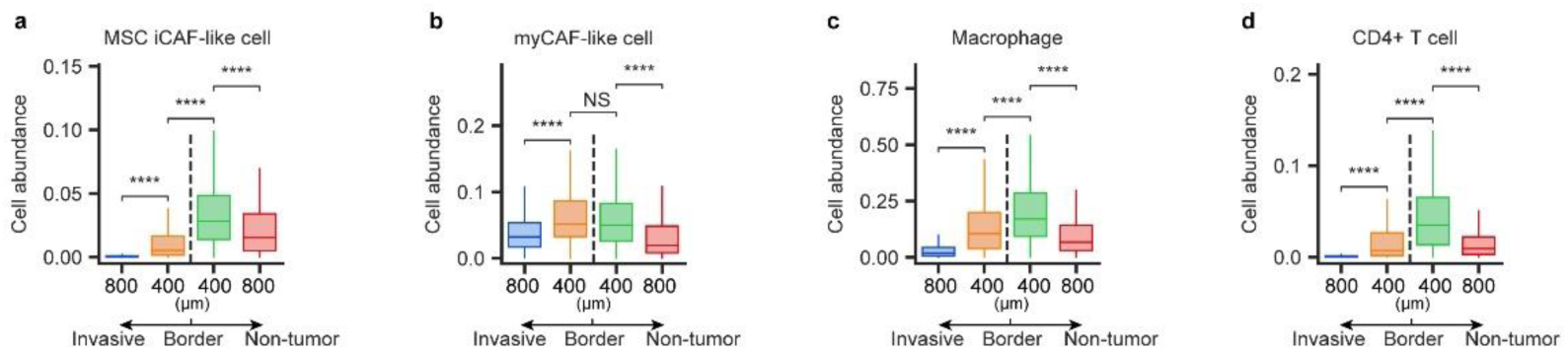
Cell abundance in each 400 μm-width margin area around the borderline. **a**, MSC iCAF-like cell abundance. **b,** myCAF-like cell abundance. **c**, Macrophage abundance. **d**, CD4+ T cell abundance.

**Supplementary Fig. 30.**
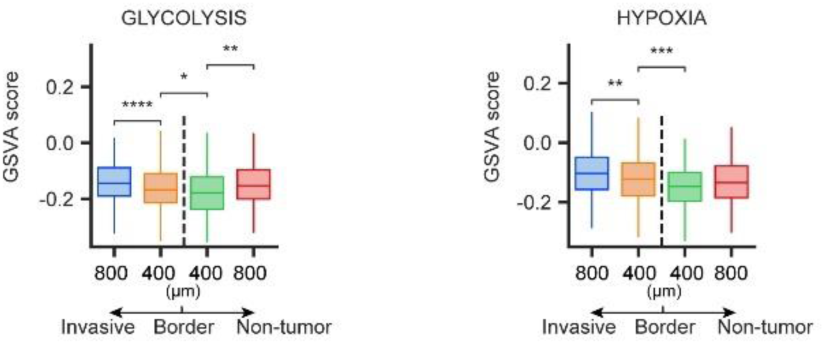
Boxplots showing GSVA scores of glycolysis and hypoxia in each 400 μm-width margin area around the borderline.

**Supplementary Fig. 31.**
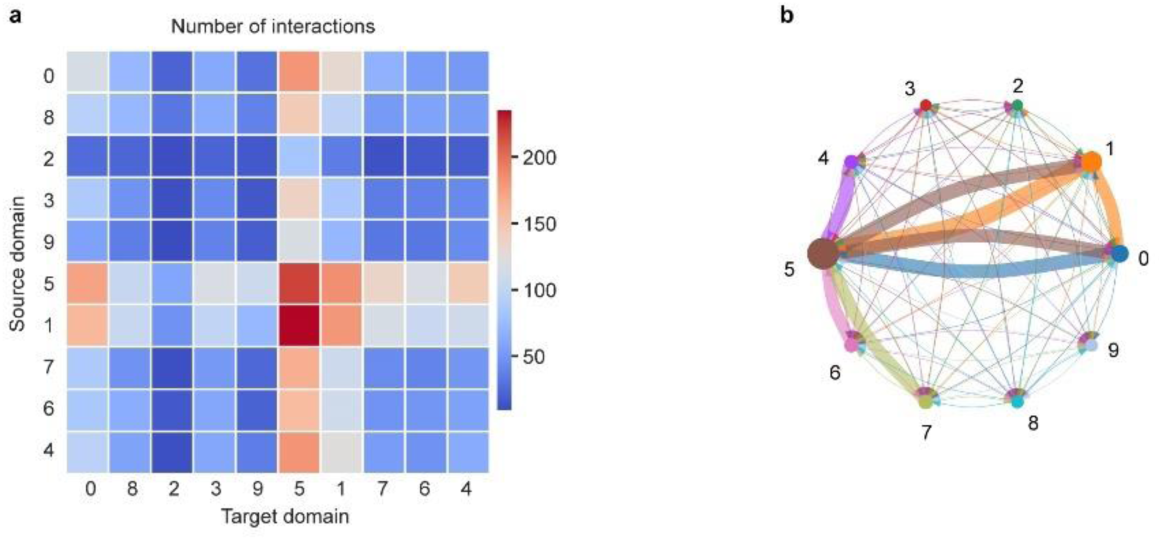
Ligand-receptor interactions in the 10x Visium IDC dataset. **a,** Interaction heatmap plotting the total number of interactions between the source (y axis) and target (x axis) domains in the 10x Visium IDC dataset. **b,** Chord plot depicting interactions between spatial domains. Size of the colored domain node represents the number of interactions from and to that domain, while edge is colored the same as the from node and its thickness denotes the number of interactions from that domain (at least 90% quantile of the distribution of the interactions are shown).

**Supplementary Fig. 32.**
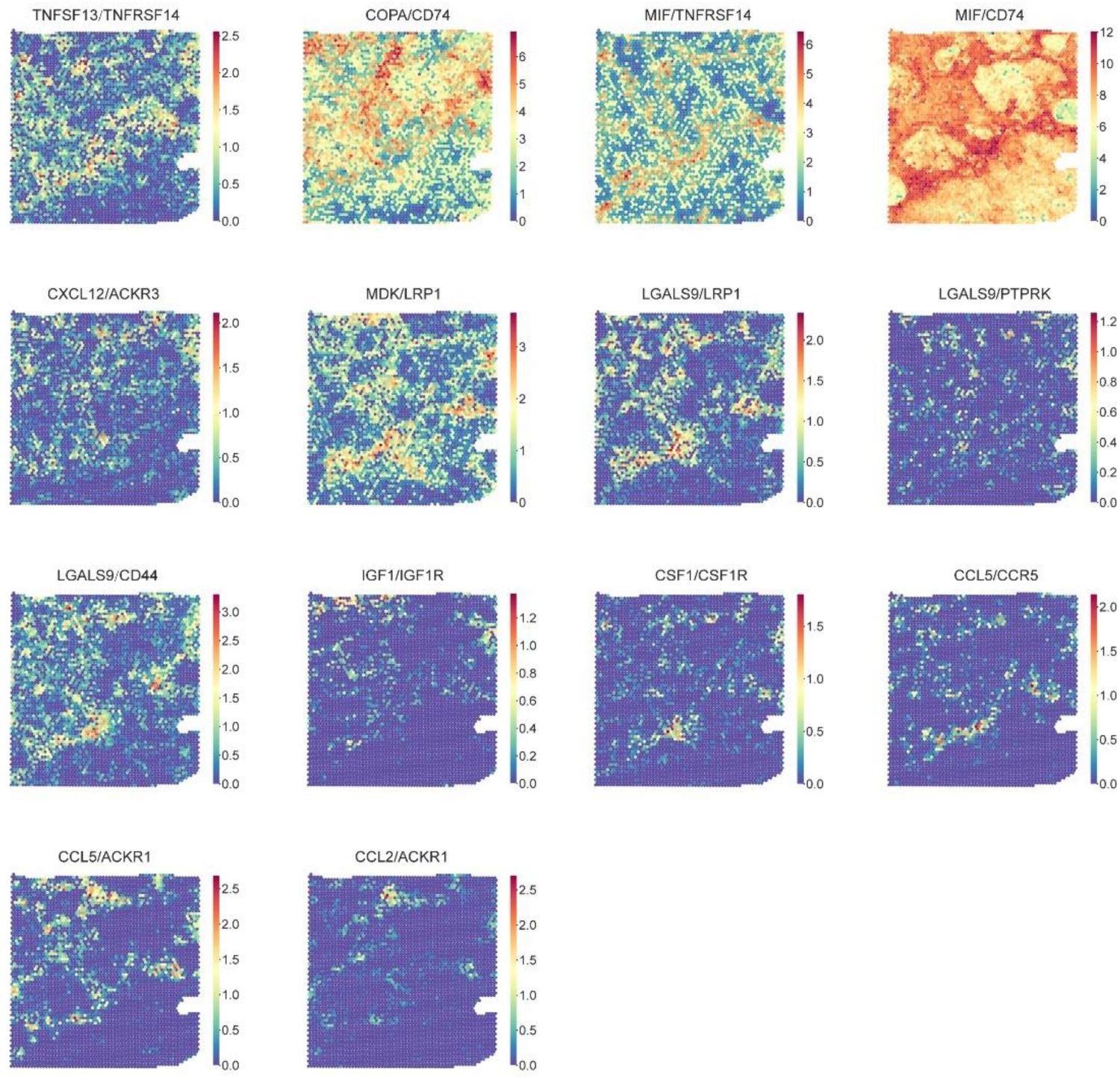
Visualization of additionally selected spot-wise LR score corresponding to those shown in Fig. 6j.

**Supplementary Fig. 33.**
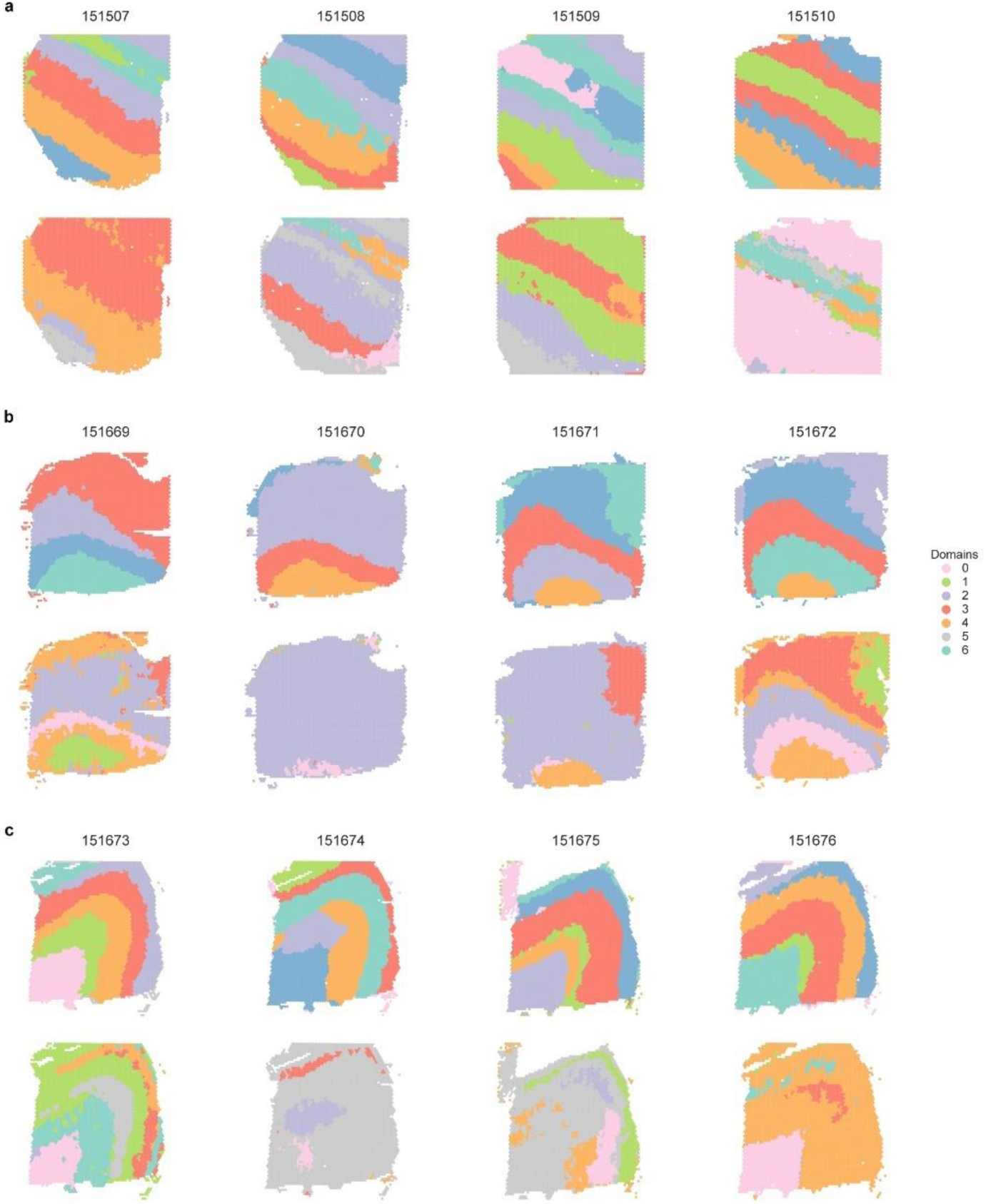
Qualitative evaluation of the identification of spatial domains by SpaSEG in 12 sections in the 10x Visium DLPFC dataset. **a**, Visualization of spatial domains in sections from 151507 to 151510. **b**, Visualization of spatial domains in sections from 151669 to 151672. **c**, Visualization of spatial domains in sections from 151673 to 151676. Each panel has two rows of spatial domain results. In the first row, spatial domains are identified by SpaSEG with all optimal parameters, and in the second row with optimal parameters except for the use of edge strength loss ℒ_edge_ (by setting *β*=0) during training iteration.

**Supplementary Fig. 34.**
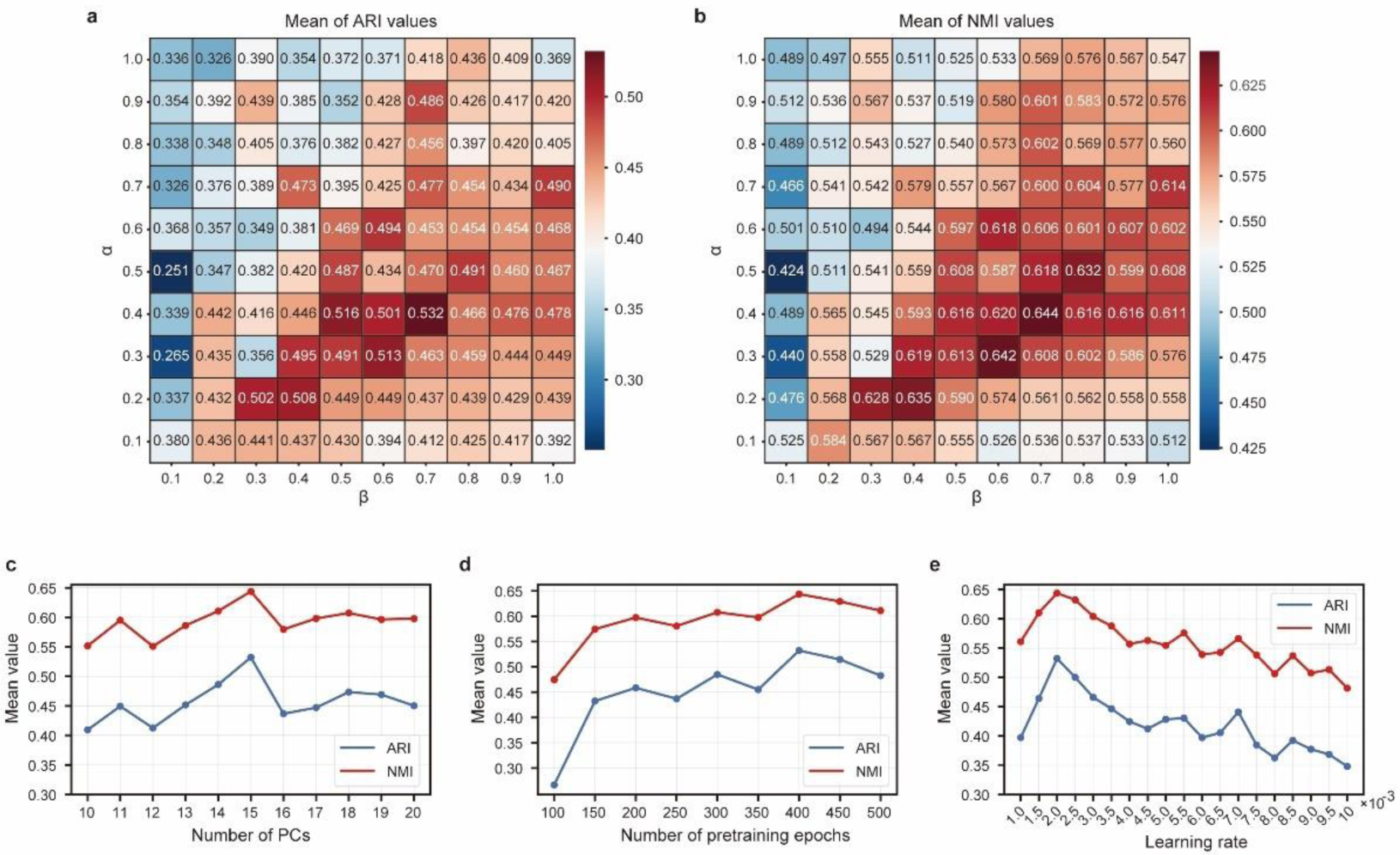
Analysis of SpaSEG hyper-parameters based on the 10x Visium DLPFC dataset. **a-b**, Mean values of ARI (**a**) and NMI (**b**) with respect to the settings of *α* and *β* used in the overall loss function. **c-e**, Mean values of ARI and NMI with respect to the number of PCs (**c**), pretraining epochs (**d**), and learning rate (**e**).

**Supplementary Fig. 35.**
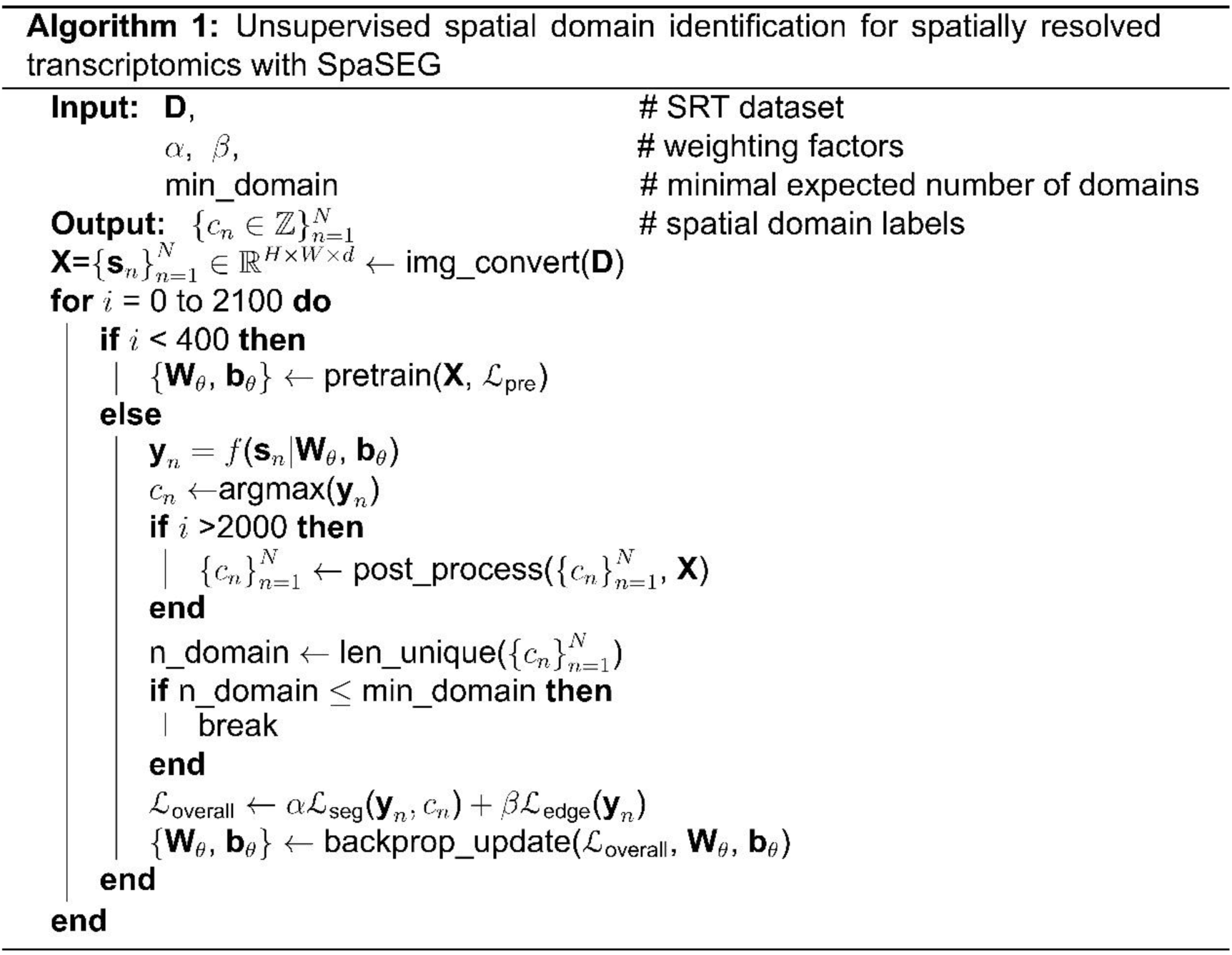
Pseudo-code of SpaSEG for the identification of spatial domains.

**Supplementary Table 1.**
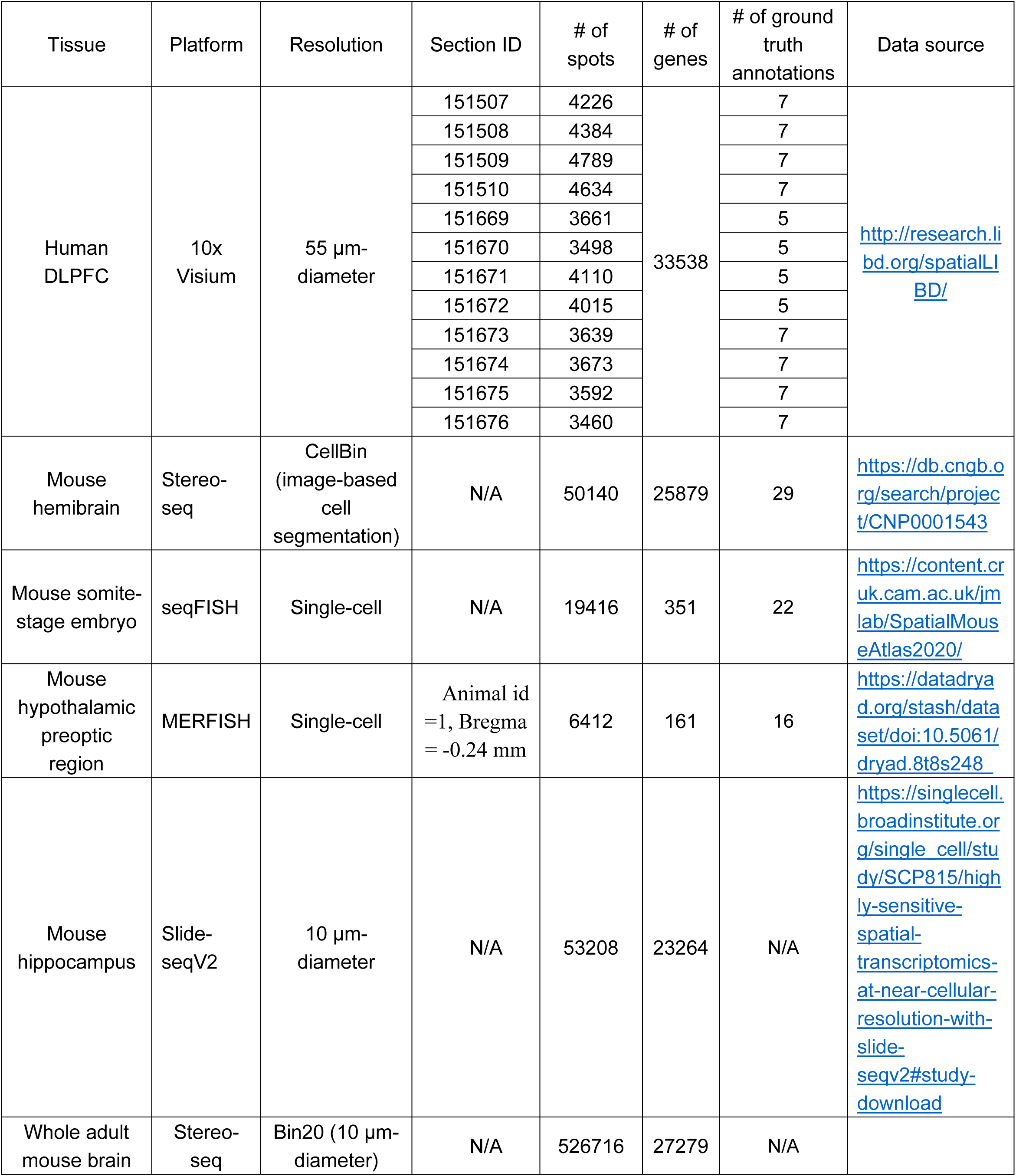
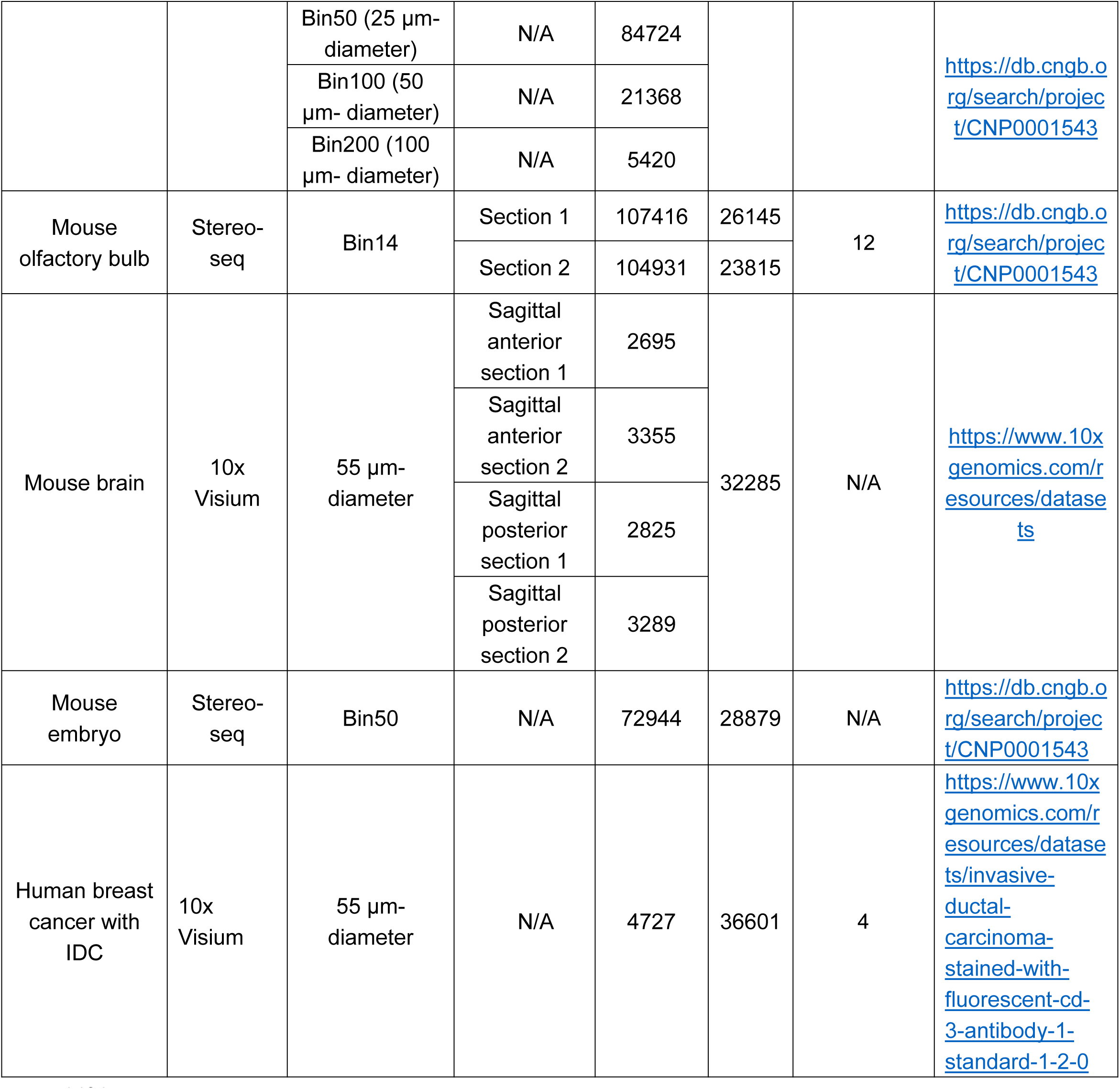
SRT datasets used in our study. N/A: not available

**Supplementary Table 2.** Results of ARI metrics from SpaSEG and competing methods to evaluate the performance of identifying spatial domains in 12 all tissue sections of the 10x Visium DLPFC dataset. The highest ARI values for different sections are highlighted. Std: standard deviation

| Section ID | Leiden | Giotto | stLearn | SEDR | SpaGCN | BayesSpace | SpaSEG |
| --- | --- | --- | --- | --- | --- | --- | --- |
| 151507 | 0.399 | 0.384 | 0.335 | 0.440 | 0.448 | **0.467** | 0.435 |
| 151508 | 0.400 | 0.390 | 0.306 | 0.350 | 0.358 | 0.437 | **0.489** |
| 151509 | 0.393 | 0.337 | 0.431 | 0.446 | 0.373 | 0.397 | **0.468** |
| 151510 | 0.393 | 0.332 | 0.441 | 0.433 | 0.408 | 0.429 | **0.528** |
| 151669 | 0.352 | 0.285 | 0.324 | 0.321 | 0.098 | 0.469 | **0.592** |
| 151670 | 0.331 | 0.274 | 0.208 | 0.406 | 0.375 | 0.431 | **0.665** |
| 151671 | 0.310 | 0.419 | 0.387 | 0.472 | 0.544 | **0.744** | 0.520 |
| 151672 | 0.286 | 0.391 | 0.339 | 0.465 | **0.543** | 0.425 | 0.526 |
| 151673 | 0.335 | 0.291 | 0.306 | 0.522 | 0.457 | 0.546 | **0.554** |
| 151674 | 0.377 | 0.197 | 0.408 | 0.441 | 0.320 | 0.290 | **0.561** |
| 151675 | 0.398 | 0.347 | 0.390 | 0.520 | 0.354 | **0.523** | 0.492 |
| 151676 | 0.322 | 0.358 | 0.364 | 0.464 | 0.342 | 0.364 | **0.555** |
| Median | 0.365 | 0.342 | 0.352 | 0.444 | 0.374 | 0.434 | **0.527** |
| Std | 0.040 | 0.063 | 0.065 | 0.059 | 0.117 | 0.112 | 0.061 |

**Supplementary Table 3.** Results of NMI metric from SpaSEG and competing methods to evaluate performance the of identifying spatial domains in all 12 tissue sections of the 10x Visium DLPFC dataset. The highest NMI values reached by the method for different sections are highlighted. Std: standard deviation.

| Section ID | Leiden | Giotto | stLearn | SEDR | SpaGCN | BayesSpace | SpaSEG |
| --- | --- | --- | --- | --- | --- | --- | --- |
| 151507 | 0.487 | 0.545 | 0.530 | 0.574 | 0.559 | 0.625 | **0.634** |
| 151508 | 0.466 | 0.527 | 0.522 | 0.414 | 0.491 | 0.598 | **0.622** |
| 151509 | 0.519 | 0.504 | 0.612 | 0.593 | 0.538 | 0.592 | **0.643** |
| 151510 | 0.474 | 0.489 | 0.592 | 0.567 | 0.540 | 0.554 | **0.651** |
| 151669 | 0.490 | 0.403 | 0.495 | 0.514 | 0.292 | 0.605 | **0.643** |
| 151670 | 0.411 | 0.408 | 0.394 | 0.531 | 0.487 | 0.551 | **0.602** |
| 151671 | 0.451 | 0.507 | 0.546 | 0.590 | 0.657 | **0.703** | 0.631 |
| 151672 | 0.430 | 0.531 | 0.472 | 0.573 | 0.631 | 0.568 | **0.633** |
| 151673 | 0.479 | 0.473 | 0.505 | 0.641 | 0.618 | **0.680** | 0.674 |
| 151674 | 0.489 | 0.333 | 0.566 | 0.612 | 0.484 | 0.478 | **0.671** |
| 151675 | 0.479 | 0.471 | 0.584 | 0.612 | 0.503 | **0.676** | 0.655 |
| 151676 | 0.434 | 0.467 | 0.534 | 0.607 | 0.530 | 0.560 | **0.667** |
| Median | 0.477 | 0.481 | 0.532 | 0.582 | 0.534 | 0.595 | **0.643** |
| Std | 0.031 | 0.062 | 0.059 | 0.060 | 0.094 | 0.064 | 0.021 |

**Supplementary Table 4.** Optimal hyper-parameters of SpaSEG for the single-section analysis.

| Platform | Dataset(s) | # of PCs | Weighting factors in the overall loss function |  | # of expected domains | coordinate scale factor |
| --- | --- | --- | --- | --- | --- | --- |
| | | | $\alpha$ | $\beta$ | | |
| 10x Visium | Human DLPFC sections from 151507 to 151510 | 15 | 0.4 | 0.7 | 7 | N/A |
|  | Human DLPFC sections from 151669 to 151672 | 15 | 0.4 | 0.7 | 5 | N/A |
|  | Human DLPFC sections from 151673 to 151676 | 15 | 0.4 | 0.7 | 7 | N/A |
|  | Human breast cancer with IDC | 15 | 0.4 | 0.7 | 10 | N/A |
| seqFISH | Mouse somite-stage embryo | 100 | 0.4 | 0.7 | 22 | 400 |
| MERFISH | Mouse hypothalamic preoptic region | 50 | 0.4 | 0.7 | 14 | 500 |
| Slide-seqV2 | Mouse hippocampus | 100 | 0.4 | 0.7 | 15 | 500 |
| Stereo-seq | Mouse hemibrain | 100 | 0.8 | 0.3 | 830 | 14 |
|  | Whole adult mouse brain at Bin20 resolution | 100 | 0.5 | 0.8 | 20 | N/A |
|  | Whole adult mouse brain at Bin50 resolution | 100 | 1 | 0.3 | 20 | N/A |
|  | Whole adult mouse brain at Bin100 resolution | 50 | 0.2 | 0.8 | 20 | N/A |
|  | Whole adult mouse brain at Bin200 resolution | 50 | 0.8 | 0.3 | 20 | N/A |
|  | Mouse embryo at Bin50 resolution | 100 | 0.9 | 0.7 | 30 | N/A |

**Supplementary Table 5.**
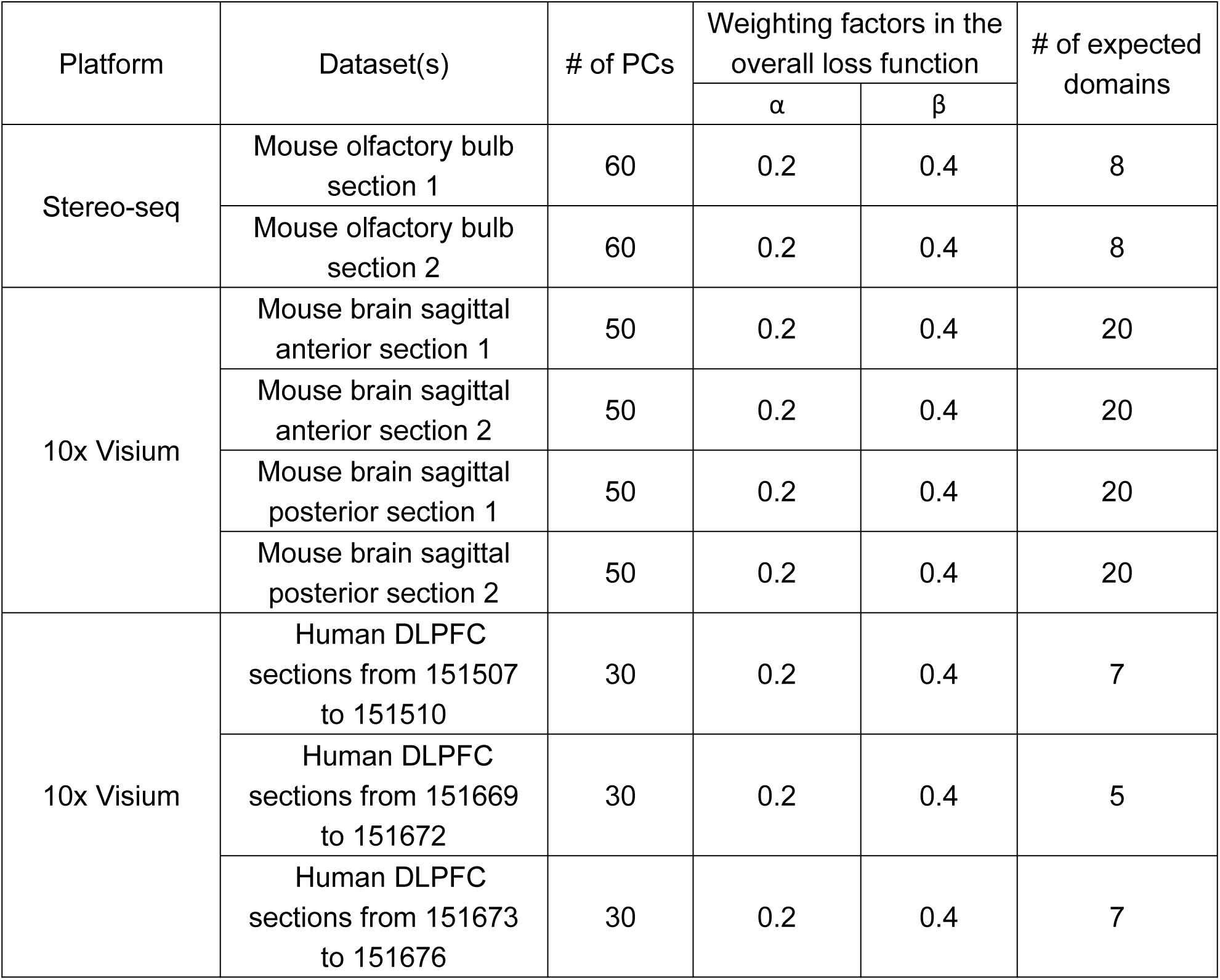
Optimal hyper-parameters of SpaSEG for the integrative analysis of multiple adjacent sections.

| Platform | Dataset(s) | # of PCs | Weighting factors in the overall loss function |  | # of expected domains |
| --- | --- | --- | --- | --- | --- |
| | | | $\alpha$ | $\beta$ | |
| Stereo-seq | Mouse olfactory bulb section 1 | 60 | 0.2 | 0.4 | 8 |
|  | Mouse olfactory bulb section 2 | 60 | 0.2 | 0.4 | 8 |
| 10x Visium | Mouse brain sagittal anterior section 1 | 50 | 0.2 | 0.4 | 20 |
|  | Mouse brain sagittal anterior section 2 | 50 | 0.2 | 0.4 | 20 |
|  | Mouse brain sagittal posterior section 1 | 50 | 0.2 | 0.4 | 20 |
|  | Mouse brain sagittal posterior section 2 | 50 | 0.2 | 0.4 | 20 |
| 10x Visium | Human DLPFC sections from 151507 to 151510 | 30 | 0.2 | 0.4 | 7 |
|  | Human DLPFC sections from 151669 to 151672 | 30 | 0.2 | 0.4 | 5 |
|  | Human DLPFC sections from 151673 to 151676 | 30 | 0.2 | 0.4 | 7 |

